# Stochastic variation in surface protein expression diversifies *Trypanosoma cruzi* infection

**DOI:** 10.1101/2025.04.07.647584

**Authors:** Lissa Cruz Saavedra, Mira Loock, Luiza Berenguer Antunes, Igor Cestari

## Abstract

*Trypanosoma cruzi* possesses hundreds of genes associated with its pathogenesis. The extent and organization of this diverse gene repertoire, expression, and role in infection remain unclear. Using accurate long-read sequencing and chromatin conformation capture, *T. cruzi* chromosomes were assembled from telomere-to-telomere. The genome revealed multigene families of virulence genes accounting for ∼70% of some chromosomes, organized in clusters or scattered through housekeeping genes. Quantitative proteomics identified stage-specific proteins and numerous trans-sialidases upregulated in trypomastigotes. Notably, the expression of virulence gene families changed stochastically in trypomastigotes, conferring *T. cruzi* fitness to cardiomyocyte infection while diversifying the invasion to multiple tissues. A *T. cruzi* genome-wide yeast surface display screen against Chagas disease patients’ antibodies revealed virulence genes expressed during human infections. However, limited conservation in their antibody-binding sites suggests their sequence diversity and variation help parasites avert antibody recognition. The data point to a role for virulence multigene families in infection persistence.

## Introduction

Many organisms evolved large gene families for specialized survival functions or to adapt to environmental changes. This is the case for odorant receptors in mammals (1), ubiquitin-like proteins in some bacteria (2), or pathogens’ variant surface antigens for immune evasion (3). Their expansion results from gene duplication and recombination events, with their chromosome organization typically influencing their expression (4, 5). The pathogen *Trypanosoma cruzi*, a single-celled protozoan parasite, has six prominent multigene family (MGF) groups encoding dozens to hundreds of virulent genes, including trans-sialidases (TS), mucins, mucin-associated surface proteins (MASPs), dispersed gene family 1 (DGF-1), retrotransposon hot spot (RHS) and the zinc metalloprotease gp63 (GP63), accounting for ∼20% of the genes in the genome (6-8). MGFs are associated with *T. cruzi* pathogenesis (9), but their function and control of expression are poorly understood, partly due to their repetitive and highly variable sequences, complex genome organization, and diversity among strains.

*T. cruzi* causes Chagas disease, affecting ∼8 million people primarily in South and Central America, but has spread to the United States, Canada, Europe, and Asia (10). A reduviid insect vector transmits the parasite from small mammal reservoirs to humans; however, it can also spread through congenital transmission, blood transfusion, or food contamination. Chagas disease has a short acute stage that can evolve into a decades-long chronic disease. The host immune response against *T. cruzi* entails parasite-specific antibody-mediated attacks and CD8^+^- and CD4^+^-T cell responses (11-13). While the immune response controls parasitemia, it does not eliminate parasites that persist in certain tissues. This leads to chronic disease, with approximately ∼30% of cases progressing to cardiac or gastrointestinal complications, often resulting in patient disability or death (14). Nifurtimox and benznidazole are drugs used for treatment, but they are mainly effective in the acute stage (15).

The mechanisms underlying *T. cruzi*’s ability to infect multiple tissues and cause persistent infections are unclear. *T. cruzi* surface is heterogeneous, with multiple glycosylated proteins such as TSs, mucins, and MASPs usually attached to the cell surface by glycosylphosphatidylinositol anchors (16). Some of these proteins are virulence surface factors involved in immune evasion and host cell infection, and others are highly immunogenic and shed in the host bloodstream (17-22). Active TSs, for example, transfer host sialic acid to the parasite surface (22, 23), contributing to complement evasion (24), cell invasion (17) and cell egress (17). Some enzymatically inactive TSs and mucins play a role in host cell receptor recognition and invasion (25, 26), modulation of host cell apoptosis (27) and the immune system responses (28, 29). It is still unclear how the surface protein repertoire of *T. cruzi* changes in various developmental stages and throughout infection. Given MGFs’ extensive repertoire and amino acid sequence diversity, a plausible conjecture is that variation in the genes expressed helps the parasite invade multiple tissues or evade host immune responses, contributing to infection persistence.

To gain insights into the organization of MGFs encoding *T. cruzi* virulence genes, we generated a high-resolution genome assembling complete chromosomes of the *T. cruzi* Sylvio X10 strain. The genome defined the extent and locations of MGF genes, typically organized as gene clusters spread throughout the chromosomes. Quantitative proteomics revealed stage-specific MGF subsets with an upregulation of TSs in trypomastigotes. A temporal nanopore-based MGF-seq and proteomic analysis in trypomastigotes revealed stochastic expression variation of MGFs with changes occurring every round of cell infection. The changes concurred with variable rates of *T. cruzi* invasion in multiple tissues, and increased fitness in cardiomyocytes. A *T. cruzi* genome-wide yeast surface display (YSD) screen against antibodies from Chagas disease patients identified MGFs expressed during human infections. Notably, their antibody-recognition sites exhibited limited conservation, implying that sequence diversity and expression variation might contribute to antibody evasion. The data revealed the genomic organization of MGFs, their stage-specific expression, and stochastic variation in trypomastigotes, indicating that their sequence diversity and variation contribute to parasite immune evasion and diversify tissue infection.

## Results

### High-resolution genome assembly reveals chromosome-level multigene family organization

To understand the organization and number of MGF genes in *T. cruzi*, we sequenced and *de novo* assembled the genome of the TcI strain Sylvio X10. TcI is widespread from South to North America and is typically associated with cardiovascular disease (30). The genome was assembled using PacBio HiFi sequencing and chromatin conformation capture with nanopore sequencing (Pore-C) (Fig. 1A). Additional nanopore sequencing was used to validate the assembly. The assembled nuclear genome was 38.1 Mb, with an N50 of 1.30 Mb and the largest contig being 2.63 Mb. It included 13,798 genes, of which 2,934 encoded MGFs (Fig. 1A, Table 1). Thirty-one chromosomes were assembled, of which 27 included end-to-end telomeres and four had one telomere (Fig. 1A). Ploidy analysis by depth (average 397x) revealed 29 diploid chromosomes, with chromosome 22 having one copy per haplotype (Fig. 1B, Fig. S1), one of which includes an insertion of ∼126 kb containing L1Tc genes followed by TSs. In contrast, chromosomes 1 and 30 were haploid and triploid, respectively. The mitochondrial maxicircle genome was hexaploid, with a complete set of genes and repeat sequences (Fig. S2). Sequence depth confirmed chromosome copy numbers and indicated segmental aneuploidy in chromosome 1 (Fig. S1). Repeat sequences higher than 70% were observed in chromosomes 9, 12, 29, and 31 (Fig. 1A), all rich in MGF sequences. There were 39 short-length scaffolds ranging from 6 to 68 kb. Synteny analysis revealed they represented segmental aneuploidy or haplotype differences, including non-syntenic insertions and MGF duplications (Fig. 1C, Fig. S3).

**Table 1.**
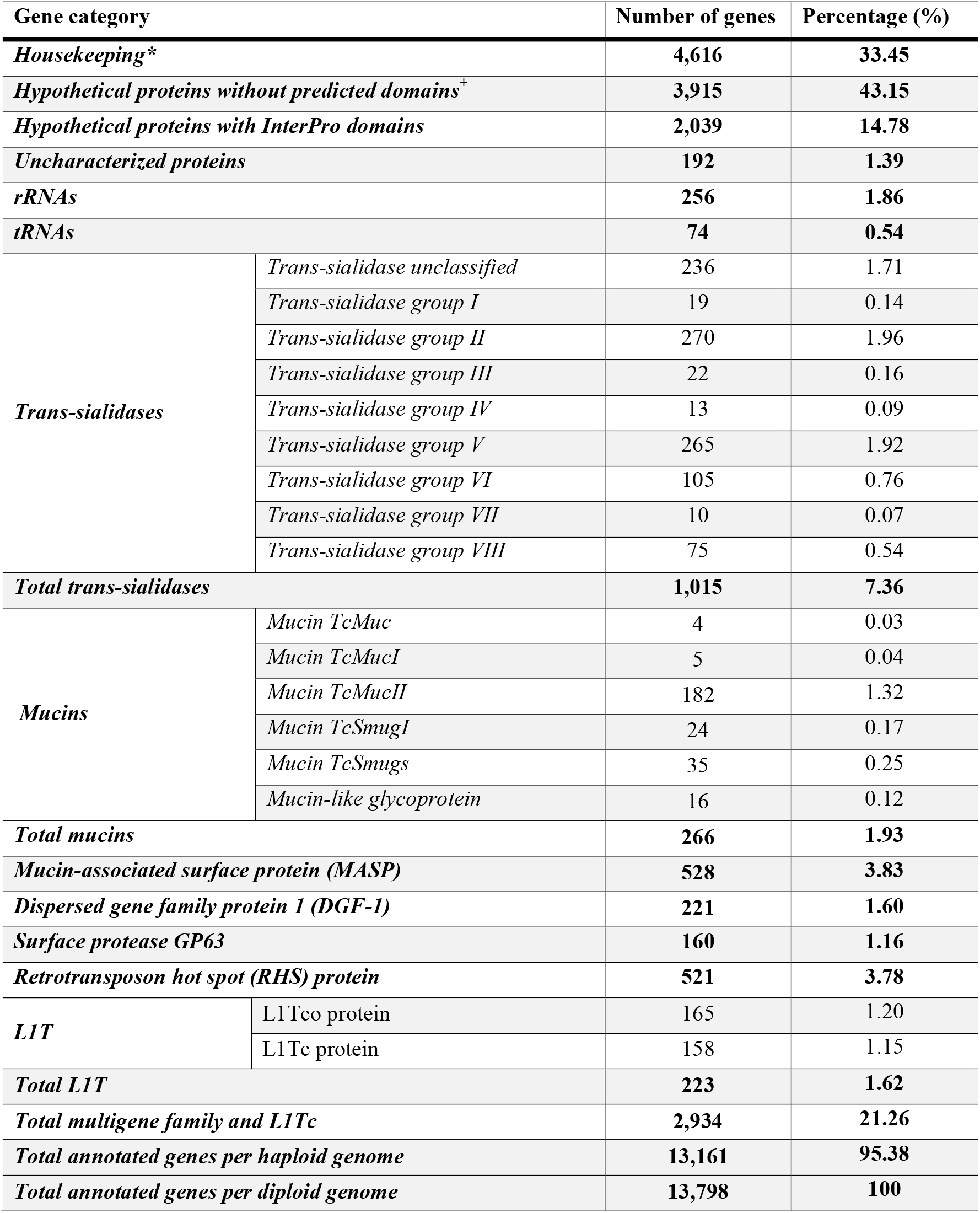
Distribution of annotated genes in *T. cruzi* Sylvio X10 genome. The haplotype genome, which includes chromosome 22, has 13,161 annotated genes. Chromosome 22b contains an additional 637 genes, resulting in 13,798 annotated genes in the genome. * Includes all protein-coding genes except hypothetical proteins and multigene families. ^+^ The total number of genes encoding hypothetical proteins is 5,954.

**Figure 1.**
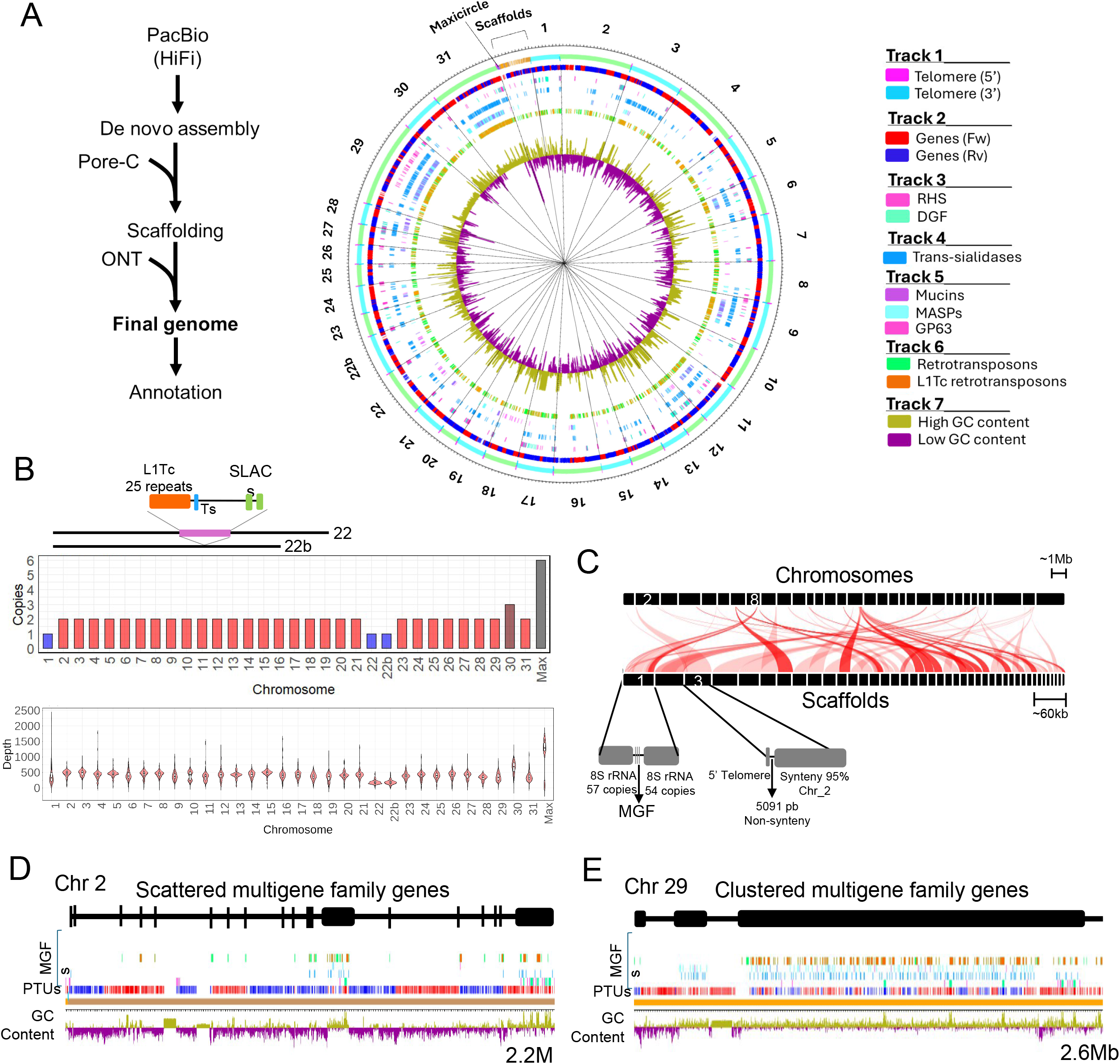
Assembly of *T. cruzi* Sylvio X10 genome. A) The diagram (left) shows the steps to assembling the *T. cruzi* Sylvio X10 genome. The assembled genome’s circular map (right) shows the distribution of genes, MGFs, telomeres, and GC content throughout the 31 chromosomes, 39 scaffolds, and maxicircles. B) Quantification of chromosome copies (top) and sequence depth per chromosome. The diagram indicates differences between chromosome 22 haplotypes designated as 22 and 22b. C) Synteny analysis of scaffolds against all chromosomes. D-E) The diagrams show chromosome 2 with scattered (D) and chromosome 29 (E) with an expanded and clustered MGF gene distribution. The diagrams follow the same colour code indicated in the circular plot legend in A.

The chromosomes were primarily organized into clusters containing housekeeping and hypothetical (HK) genes, alternating with clusters of MGF genes (Fig. 1A, D-E). There were ∼70 genes per HK cluster and ∼18 genes per MGF cluster, and HK genes were syntenic across multiple *T. cruzi* strains (Fig. S4). MGFs and L1Tc-co retrotransposon accounted for ∼21% of the annotated genes (Table 1). The most abundant members were the TSs (1,015 genes), particularly those from groups II and V. These were followed by MASPs (528 genes), RHS (521 genes), mucins (266 genes) – with TcMucII being the most abundant group – and L1Tc-co retrotransposons (223 genes). Chromosomes 6, 9, 11, 12, 29, and 31 contained over 70% of their sequences as repeats and ∼60% of the genes encoded MGFs, with notable disruptions in the organization of forward (red) and reverse (blue) strands (Fig. 1A and E, Fig. S1). Other chromosomes, such as chromosome 2, contained primarily HK genes (Fig. 1D). Mucins, MASPs, TSs, and GP63 were distributed in MGF clusters or scattered throughout multiple chromosomes (Fig. 1D-E). In contrast, DGF-1 and RHS were enriched in subtelomeric regions (Fig. 1A, D-E), indicating potential differences in gene expression regulation. MGF sequences correlated with a high GC content and were rich in LINE/I-L1Tc retrotransposons and RHS (Fig. 1A, D-E). The data provide a high-resolution genome assembly and insights into MGF’s widespread chromosomal organization. The alternating clusters of MGF and HK genes likely reflect an evolutionary pressure to maintain MGF diversity and expression.

### Tandem mass tag quantitative proteomics reveals stage-specific multigene family expression

We performed multiplexed quantitative proteomics to determine differences in protein expression among *T. cruzi* stages and identify expressed MGF proteins. We used tandem mass tags (TMT) to quantify proteins of each parasite life cycle stage, i.e., non-infectious epimastigotes (EP), column-purified infectious metacyclic trypomastigotes (MT), amastigotes (AM), and tissue culture-derived trypomastigotes (CT), representing the main developmental stages (Fig. 2A). Proteins were extracted, trypsin-digested, and peptides labeled with isobaric (^15^N, ^13^C, and ^2^H) but molecularly identical tags containing a reactive group, a space arm for mass normalization, and a mass reporter (from 126–135 Da) followed by LC-MS/MS to identify peptides and quantify the abundances of reporter ions (Fig 2B). Principal component analysis (PCA) revealed differences among each parasite life stage and consistency within biological replicates (Fig. 2C). The dataset comprised 42,348 peptides and 5,993 proteins with quantifiable differences among various parasite stages (Table S1). Hundreds of proteins in various cellular processes were up- and down-regulated throughout the life stages, defining stage-specific and common proteomes (Fig. 2D, Fig. S5). They included known stage-specific markers, such as GP82 and GP90 in MTs (21), trans-sialidases in CTs (22, 23), amastin in AMs (31), proteins involved in energy metabolism in EPs (Fig. 2D-F), and known vaccine candidates (e.g., Tc24) and diagnostic targets (e.g., HSP70 and histones) (32) (Fig. 2E, Table S1). The differentially expressed proteins between infectious and non-infectious forms (Fig. 2F, Fig. S5) revealed ∼600 potential virulence factors, as well as metabolic, morphological, and surface coat proteins associated with infectious stages.

**Figure 2.**
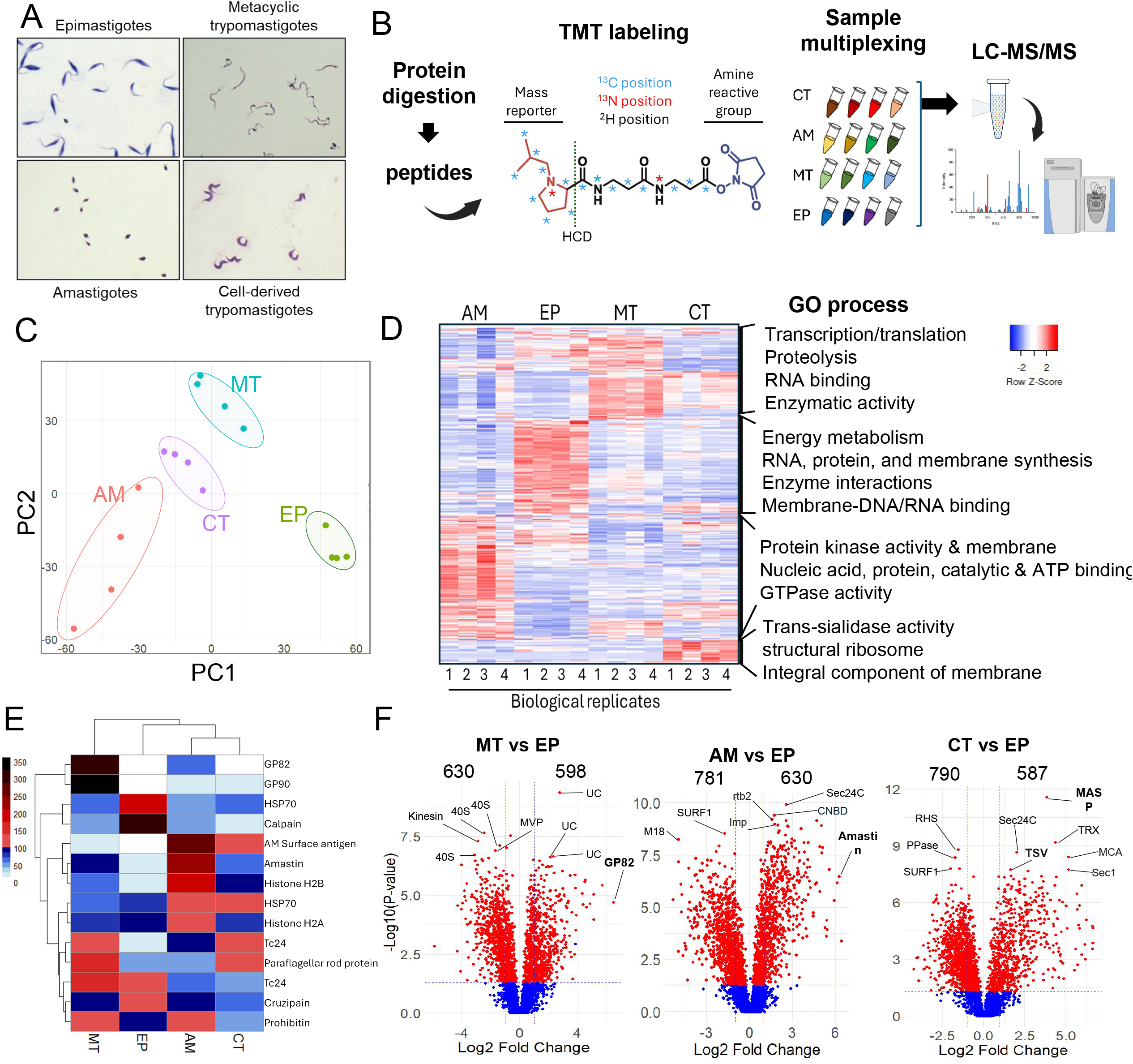
Quantitative proteomics of *T. cruzi* life cycle stages. A) Giemsa staining of the four main stages of *T. cruzi* used for proteomics. MT and AMs were purified using DEAE-Sephadex (46). B) Diagram of TMT-labeling and mass spectrometry. Proteins from each life stage were extracted (four biological replicates), trypsin digested, and peptides labelled with a 16-plex isobaric molecule with an amine-reactive group and mass reporter. After labelling, samples were mixed and co-analyzed by mass spectrometry. C) PCA plot of the mass spectrometry results of each parasite stage. Each dot represents a biological replicate. D) Heatmap of total protein expression change; it does not include MGFs. The most represented gene ontology (GO) processes are indicated on the right. E) The heatmap shows the expression of selected known stage-specific proteins, vaccine candidates, and diagnostic targets. F) Volcano plots show a comparison of protein expression between infectious stages (MT, CT, AM) and the non-infectious stage (EP). Red dots are significant changes, i.e., fold-change ≥ 2 and *p-*value ≤0.05, whereas blue dots are non-significant. Stage-specific markers are indicated in bold (GP82, Amastin, MASP, TS). The data show the results of four biological replicates.

We identified a stage-specific pattern of MGF protein expression (Fig. 3A), characterized by variations in protein abundance across each stage. TSs, mucins, and MAPSs were significantly upregulated in CTs with many TS proteins expressed (Fig. 3A-B, Fig. S6). In contrast, DGF-1 and RHS were upregulated in AMs, MTs, and EPs, whereas GP63 was up-regulated in AMs and CTs (Fig. S6). Analysis of TS subsets showed an enrichment of all TS groups in CTs compared to other stages (Fig. 3C). The MGF proteins expressed in CTs were encoded by genes distributed across multiple chromosomes rather than a single locus (Fig. 3D-E). However, chromosomes 6, 9, 29, and 31, enriched in MGFs, contributed to a large proportion and exhibited higher MGF abundance in CTs compared to other stages (Fig. 3D, Fig. S7). RNA-seq and proteomic analysis confirmed the expression of large subsets of MGF genes from multiple chromosome loci (Fig. 3E, Fig. S8). The diversity of MGF proteins expressed likely reflects heterogeneity within the parasite population. Analysis of mRNA abundance in CTs showed a bimodal distribution of TSs (Fig. 3F), distinct from other MGF and housekeeping proteins (Fig. S9), with RNA levels around 1.2 and 5 logs (Fig. 3F). The high abundance of TS mRNAs correlated with the distribution of TS proteins expressed, suggesting a threshold accumulation in mRNA levels for their translation. The data revealed stage-specific *T. cruzi* proteome, identifying stage-specific subsets of MGF proteins and an upregulation of TSs in CTs. It also indicates that MGFs are expressed from multiple locations on various chromosomes.

**Figure 3.**
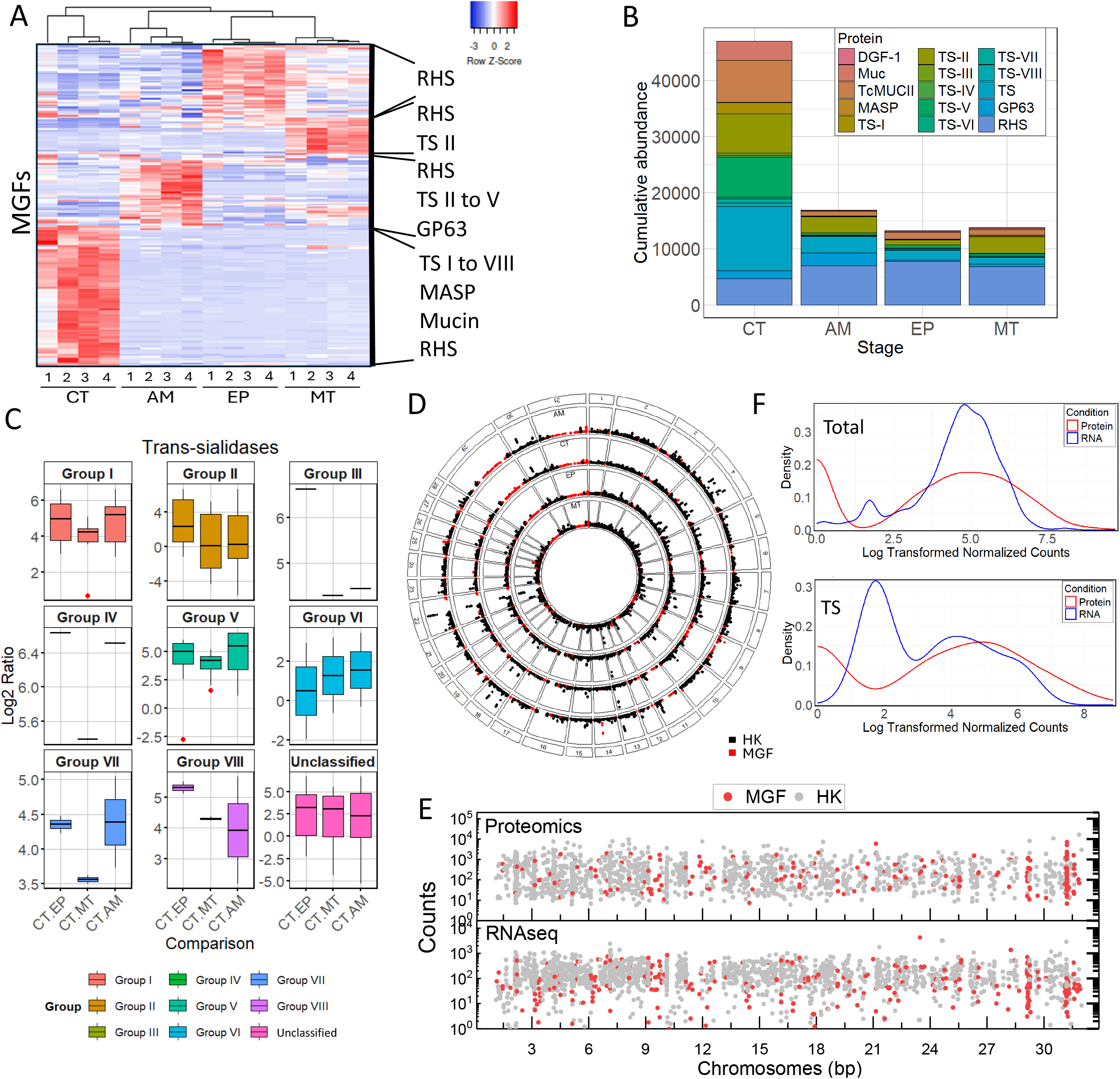
MGF proteins are expressed from various chromosome loci. A) Heatmap of MGF protein expression by TMT-labeling proteomics. B) Cumulative abundance of MGF proteins quantified by TMT-labeling proteomics in each life cycle stage. C) Quantitative comparison of TS protein subgroups between CT vs EP, CT vs MT, and CT vs AM. The graphs display a 25-75% distribution (box), mean (horizontal lines), and standard deviation (vertical lines) for four biological replicates. All TSs within a group were combined for the comparisons. D) The circular plot shows the chromosome location associated with MGF (red) and HK (black) protein expression (mean of four biological replicates) at each stage of the parasite. E) Mean RNA and protein abundance counts in all chromosomes. F) Distribution of total cell transcriptome (top) and TSs (bottom) in CTs. The data show the results of four biological replicates.

### Stochastic variation in the multigene family’s expression diversifies *T. cruzi* cell invasion

We postulated that MGF gene expression varies temporarily in CTs after every round of cell infection, resulting in heterogeneous parasite populations. We developed an MGF-seq approach to select all expressed MGF mRNAs and determine their expression changes in CTs. We designed sets of primers targeting the 3’ conserved ends of MGF genes and the splice leader sequence (Table S2), attached to the 5’ UTRs of all trypanosome mRNAs (33), followed by nanopore sequencing. We infected H9-C2 cardiomyocyte cells with MTs and collected the generated CTs (Fig. 4A). Then, we performed four rounds of host cell infection with CTs accompanied by MGF-seq at each round to investigate MGF expression changes (Fig. 4A). The approach identified 2,465 MGF transcripts, ∼91% of all MGF genes annotated in the genome, and 2,029 (∼82%) transcripts varied throughout CT generations (Fig. 4B). Some transcripts appeared more often than others, although still varied in expression (Fig. 4B brackets). There were twice as many MGF genes expressed in CTs (1,301 ± 436 genes) as in MTs (686 genes) (Fig. 4B, Fig. S10), with 154 TSs expressed at any time in the CT population, representing ∼15% of the TS repertoire. However, TS expression levels varied by up to ∼1,700-fold and changed throughout CT generations or biological replicates, with all subgroups represented (Fig. 4B, Fig. S10), indicating TS heterogeneity and stochastic variation within the parasite population. TMT-labeling proteomics over CT generations confirmed MGF expression changes at the protein level (Fig. 4C), confirming diversity and variation in MGF expression. It is likely that individual parasites express only subsets of the MGFs and vary their expression throughout the course of infection.

**Figure 4.**
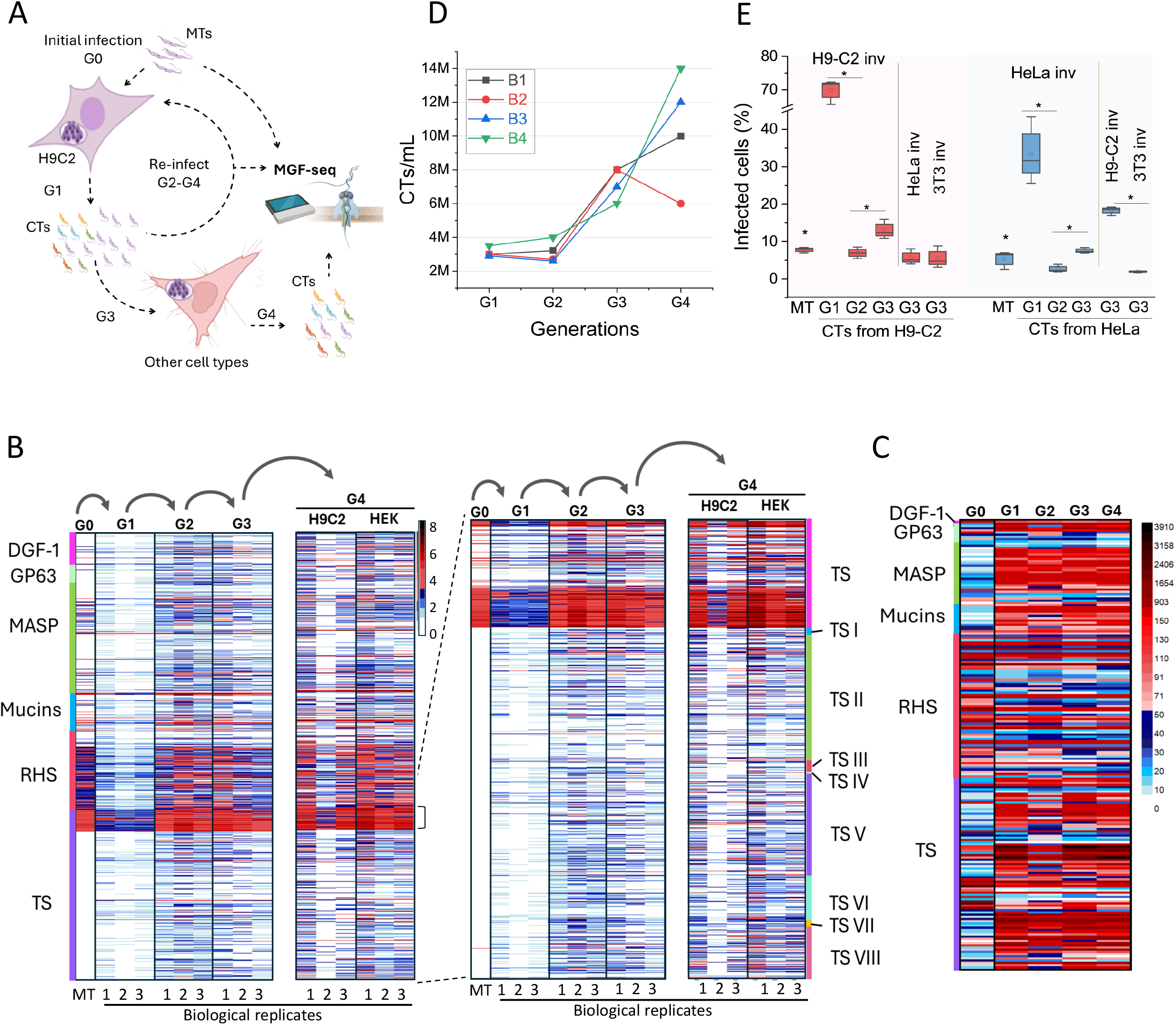
Stochastic variation in MGF gene expression in trypomastigotes. A) Diagram of MGF-seq experimental design. H9-C2 cardiomyocyte cells were infected with MTs, and CTs were collected from the supernatant and used to re-infect H9-C2 cells. The process was repeated four times, and CTs were collected for MGF-seq each time. After the third round of infection, HEK293T cells were infected, and CTs were collected for MGF-seq. G, generation. B) Heatmap of MGF-seq from MTs and CTs. Each round of infection (G) and biological replicate is shown. On the right is an analysis of TS expression by subgroups. Unclassified TSs are labelled TS. C) Quantification of MGF proteins from MTs (G0) CTs (G1-G4) by TMT-labeling and LC-MS/MS at multiple generations as described in A. D) Quantification of CTs produced and released into the host cell supernatant each round of infection from B. In each round, 30,000 H9-C2 cells were infected (1:10, cells to parasites), and CTs were collected every 6 days post-infection. M, millions. E) Invasion assay with H9-C2, HeLa, or 3T3 cells and *T. cruzi* MTs or CTs (produced at multiple generations in H9-C2 or HeLa cells). 30,000 cells were incubated with 10 parasites per cell for 3h. The graphs display a 25-75% distribution (box), mean (horizontal lines), and standard deviation (vertical lines) for three biological replicates. * *p*-value ≤ 0.05, comparing between subsequent generations.

To determine if the stochastic variation in MGF expression affects host cell infection, we quantified the production of CTs every round of H9-C2 infection. Moreover, we performed invasion assays using H9-C2 (cardiac), HeLa (epithelial), and 3T3 (fibroblast) cells with MTs or CTs obtained from H9-C2 or HeLa cells. Interestingly, there was an increase in CT production with each sequential round of H9-C2 infection (Fig. 4D). In contrast, invasion assays showed variations in the rates of host cell invasion between MTs and CTs, or amongst CT generations and host cell types (Fig. 4E). The variation in invasion rates occurred with CTs obtained from H9-C2 or HeLa cells, implying it is independent of the host cell used to generate the CTs. The data indicate that stochastic variations in *T. cruzi* surface protein expression may diversify host cell infection, rather than selecting for a specific cell type. This may be due to changes in parasite surface proteins allowing them to engage with different host cell receptors for invasion. The increased production of CTs after consecutive H9-C2 infections (Fig. 4D) may be due to increased AM proliferation fitness rather than selection of CTs for invasion. There was a notably high rate of invasion with the first generation of CTs (G1) (Fig. 4E), which correlated with their lower diversity in MGFs expressed (Fig. 4B, Fig. S10). This suggests a more homogeneous parasite population at G1 and increased diversification in subsequent generations. The data show stochastic variation in MGF expression at both the RNA and protein levels in CTs, suggesting that this variation may diversify *T. cruzi* host cell infection.

### Diversity and immunogenicity of MGF proteins during human infections

To gain insights into MGF protein immunogenicity and expression variation during human infections, we screened a *T. cruzi* genome-wide library for yeast surface display (YSD) (33) against antibodies from patients with Chagas disease or healthy individuals (Fig. 5A). The library size corresponds to ∼30-fold the parasite genome, including all genes, and has ∼4 million clones with ∼270 clones per gene (33). The library expression is induced by galactose, resulting in parasite proteins or fragments thereof expressed on the yeast surface via the agglutinin mating system Aga2p-Aga1p (34) (Fig. 5A). The YSD library was incubated with pooled sera from chronic-stage Chagas disease patients (ChD) and compared to those from healthy individuals. Although the strains infecting the patients are unknown, MGF genes are present in all *T. cruzi* strains (35). We performed three rounds of library enrichment, followed by nanopore sequencing to identify the MGF proteins reacting with antibodies (Fig. 5A). PCA showed distinct differences in MGF proteins enriched with antibodies from patients with Chagas disease compared to healthy individuals or the non-enriched library (Fig. 5B). Comparing the screens from Chagas disease patients to healthy individuals showed sets of MGF proteins reacting specifically to Chagas disease patients’ antibodies, with a significant enrichment of TS proteins (Fig. 5C), indicating multiple MGFs expressed during human infections. The large number of TSs and MASPs is consistent with their upregulation in CTs, especially TSs (Fig. 3A-C).

**Figure 5.**
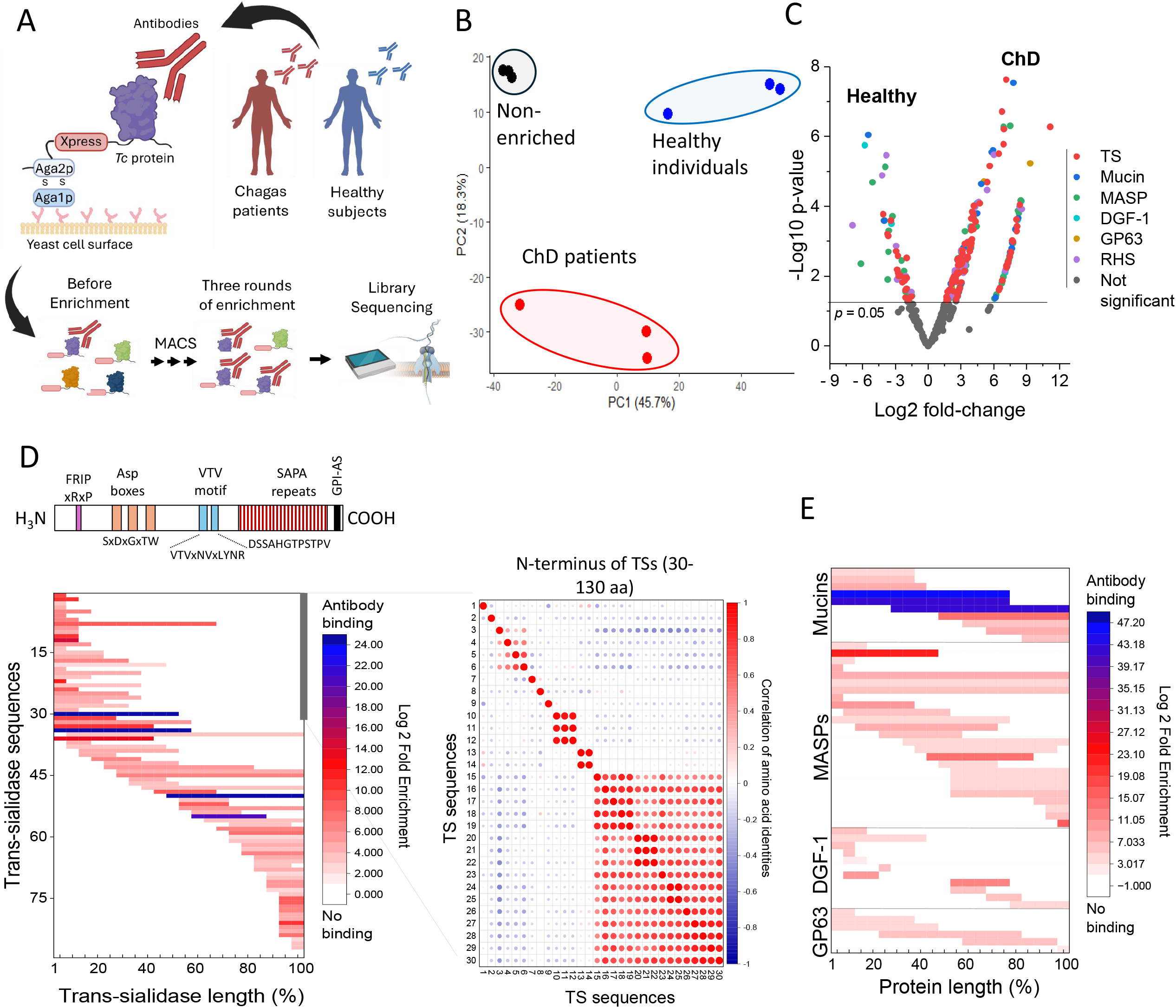
Variable MGF sequences targeted by antibodies during human infections. A) Diagram of the YSD screen with sera from Chagas disease patients or healthy individuals. Sera was incubated with the YSD library expressing *T. cruzi* antigens on the surface and enriched using magnetic-activated cell sorting (MACS). After three rounds of enrichment, the library DNAs were sequenced using Oxford nanopore technology to identify enriched sequences compared to the initial non-enriched library or the library enriched with sera from healthy individuals. B) PCA plot of each sequencing replicate (dots) by each enrichment condition. The experiment was performed in three biological replicates. C) Volcano plot shows YSD screen analysis comparing MGF proteins enriched with ChD patients’ or healthy individuals’ antibodies. ChD or healthy enriched libraries were first compared to non-enriched libraries, and non-significant hits were removed. Only MGF proteins are shown. D) Diagram of *T. cruzi* TS on top. Not all TSs have FRIP, Asp boxes, or SAPA repeats. All contain VTV motifs. Each motif’s amino acid (aa) sequences are indicated, and x means any aa. The heatmap (bottom graph) displays the location of antibody binding to the TSs identified in the YSD screen, as mapped using nanopore sequencing. TS sequence lengths were normalized from 1-100 for visualization. On the right, sequence comparison of 30 TSs with antibodies reacting at their N-terminus (grey bar). Sequence alignment of aa 30 to 130. The first 30 aa represent predicted signal peptides removed for the analysis. The percentage of identity between all 30 sequences was obtained, and their correlation was analyzed and plotted. E) The heat map displays the locations of antibody binding sites on mucins, MASPs, DGF-1, and GP63, which were identified in the YSD screen and mapped using nanopore sequencing. Protein sequences were normalized to a range of 1-100 for visualization. Experiments were performed in three biological replicates.

We analyzed the YSD library screened with antibodies from Chagas disease patients to identify antibody binding sites within the MGF proteins. Because we generated a fragment library, the antibody binding regions are enriched and can be identified by mining the enrichment of nanopore reads. Our *T. cruzi* genome contains 1,015 TSs, encompassing all eight subfamilies and an unclassified group (Table 1). TSs are highly variable in sequence (36). Those expressed in CTs may contain a FRIP motif, one to three Asp boxes, and a VTV motif, and some subfamilies contain the shed acute-phase antigen (SAPA) repeats at the C-terminus (Fig. 5D). Antibodies reacted to various regions in different TS sequences with no bias to a specific protein site (Fig. 5D). Analysis of the N-terminus of 30 TSs (100 amino acids) reacting with antibodies showed significant divergence among the antibody targeted sequences (Fig. 5D, Fig. S11), with half of the sequences showing lower than 20% identity (Fig. 5D, low correlation – blue dots) and the other half with up to 38% identify (Fig. 5D, moderate correlation – red dots), indicating a limited overlap in antibody binding sites. The same occurs in other parts of the protein.

Mucins and MASPs were more immunogenic than DGF-1 and GP63 (Fig. 5E), consistent with their upregulation in CTs (Fig. 3A-C, Fig. S6), with antibodies recognizing various protein sites. Analogous to TSs, their sequence diversity and variable expression suggest limited antibody cross-reaction. Hence, patient antibodies reacted to multiple MGFs, revealing an enrichment in TSs, and implying their expression during human infections. Their sequence diversity and variable expression might help parasites evade antibody recognition during infection.

## Discussion

*T. cruzi* expanded its large gene families encoding virulence factors to ∼3,000 genes; however, the function and evolutionary significance of this diversification remained unknown. The families’ size, repetitive sequences, and sequence similarities posed challenges in understanding their genomic organization. We generated a high-resolution genome of the *T. cruzi* Sylvio X10 strain, assembling all chromosomes and assigning precise numbers and genomic locations to MGF genes. Unlike other protozoan pathogens such as *T. brucei spp*. and *Plasmodium sp*., which evolved variant antigen families in large subtelomeric arrays (3), *T. cruzi* MGF genes are typically dispersed throughout the genome in clusters that alternate with housekeeping genes. This is evident for TSs, mucins, MASPs, and GP63, and it may facilitate their expression, as this chromosome organization may position them near other transcribed sequences. Exceptions include the DGF-1 and RHS families, which are enriched in subtelomeric regions. Notably, ∼70% of some chromosomes consisted of repeat sequences containing MGF genes, suggesting their expansion in these chromosomes.

Our proteomic analysis revealed subsets of MGF proteins that were expressed at each stage of the parasite’s life cycle. Moreover, MGFs were expressed from various loci across all chromosomes, as detected by RNA-seq and proteomics, indicating that there is no specific genomic site for their expression. The MGF genomic organization may facilitate transcription by RNA polymerase II, resulting in constitutive polycistronic transcription and the production of numerous transcripts within a cell. However, post-transcriptional levels of expression control may determine which transcripts are translated. This is consistent with the 3’-UTRs of mucins and TSs regulating their expression (37, 38). The presence of chromatin three-dimensional hubs (39) could also select subsets of gene clusters for transcription and mRNAs for processing at nuclear bodies, thereby favoring the expression of different MGF genes throughout various life stages and infection.

Although our proteomics data represent a population analysis, single-cell transcriptomics suggest that multiple MGF genes are transcribed in a single parasite (40), and an upregulation of TSs in CTs. The upregulation of TSs in CTs is also evident from our proteomic analysis and YSD screen with antibodies from Chagas disease patients. TSs function to transfer sialic acid from the host to the *T. cruzi* surface (22) helping in host cell infection and immune evasion (22). Still, not all TS genes are functional enzymes. Some TSs are involved in cell invasion (21, 22, 25), cellular egress (17) and perhaps other functions. The mechanisms regulating TS protein (and other MGFs) expression remain unknown. We found a bimodal distribution in TS transcripts, low- and high-abundance mRNAs, suggesting a threshold-level model of expression consistent with transcript abundance regulation (37, 38). In this model, the accumulation of transcripts may increase the likelihood of translation, possibly by mRNA processing and competition for the translational machinery. This aligns with data showing that most, if not all, TSs are transcribed; however, differences in mRNA production or stability could lead to their accumulation and, consequently, translation.

Our data suggest a stochastic variation in MGF mRNA and protein expression in CTs. MGF expression regulation likely resets with the parasite developmental transition from AMs to CTs, contributing to different sets of MGF expressed every round of cell infection. *T. cruzi* infects multiple cell types, interacting with host surface receptors (21, 25, 26). Diversifying surface antigens may facilitate the infection of various tissues and/or evade intracellular host cell defenses. The variation may result in parasites expressing different surface proteins within a population, thereby enhancing their ability to engage with various types of host cell receptors and invade multiple tissues. Notably, multiple rounds of H9-C2 infection by CTs resulted in increased parasite fitness, i.e., increased CT production in every subsequent round of infection, suggesting epigenetic regulation of genes that optimize parasite proliferation in H9-C2 cells, possibly by evading intracellular host cell responses or facilitating replication.

The MGF diversity and variable expression may also impact host immune recognition. The YSD screening revealed ∼150 MGF antigens recognized by human antibodies. The small subset of antigens detected (∼7% of the MGF repertoire) may reflect differences between the strain used to generate the YSD library and those infecting patients, or the immunodominance of some MGF proteins. Interestingly, analysis of antibody binding sites revealed various immunogenic regions in different TSs, indicating the absence of a specific immunogenic region. However, some regions exhibited a high antibody signal, possibly due to the presence of multiple antibodies or a high antibody titer against particular sequences. The combination of TS sequence diversity and variation in their expression could help the parasite to escape host antibody recognition. Experiments in mice or analysis of T-cells from Chagas disease patients identified different TS CD8+ T-cell epitopes (41-43). The inability of CD8+ T-cells to clear *T. cruzi* infection could also result from the variable expression of TSs and perhaps other MGFs during infection. Analysis of other MGFs revealed a high antibody signal for mucins, and a similar antibody distribution for MASPs, as observed for TSs. The diversity and temporal changes in MGF protein expression, particularly TSs in CTs, may help parasites evade antibody or T-cell recognition during infection, thereby contributing to parasite persistence. In this scenario, the MGF variation in *T. cruzi* may not be distinct from that in *T. brucei*, which is highly immunogenic and switches to evade host antibody detection (44). However, *T. cruzi* MGF expression is stochastic and likely multigenic, with multiple family members and high genome complexity, adapted for both intracellular and extracellular parasite survival. Its repertoire, in part, might have evolved to facilitate host cell invasion and evade cellular and humoral immune responses, aiding the infection of multiple tissues and promoting parasite persistence.

## Material and methods

### *T. cruzi* and mammalian cell culture

Epimastigotes (EP) of TcI *Trypanosoma cruzi* strain Sylvio X10 (ATCC 50823) were cultured in Liver Infusion Tryptose (LIT) medium, supplemented with 10% fetal bovine serum and 1% penicillin-streptomycin (45). Metacyclic trypomastigotes (MTs) were purified from a stationary 15 mL culture of EPs grown for 12 days using DEAE-Sephadex resin (46). For amastigotes (AM), H9C2 cells were infected with cell culture-derived trypomastigotes (CT) at a 1:10 ratio. The CTs were collected from the supernatant of H9C2 (ATCC CRL-1446) after 5-6 days of *T. cruzi* infection. After 10-12 days of infection, a mix of cell culture-derived trypomastigotes and amastigotes was collected from culture supernatant, and the AM were purified using DEAE-Sephadex resin (46). H9C2, HEK293-T, 3T3, and HeLa cells were cultured in DMEM medium supplemented with 10% fetal bovine serum (FBS) and 1% penicillin and streptomycin at 37°C with 5% CO[.

### Genome sequencing, assembly, and repeat calculation

For genomic DNA extraction, 3×10^7^ mid-logarithmic growth phase parasites were centrifuged at 4000 x g for 10 minutes. The resulting pellet was washed twice with 1X PBS (37 mM NaCl, 2.7 mM KCl, 4.3 mM Na2HPO4, 1.47 mM KH2PO4), resuspended in 200 μL of lysis buffer from NEB Monarch HMW DNA Extraction (NEB #T3050, Massachusetts, USA), and processed following the manufacturer’s protocol. The DNA integrity was verified by 1% agarose gel in TAE buffer (Tris 40 mM, acetic acid 20 mM, EDTA 1 mM) and quantified by Thermo Scientific™ NanoDrop™ 2000/2000c. The extracted DNA was sequenced using Pacific Bioscience (PacBio) HiFi using Single Molecule Real-Time Technology (SMRT) at Genome Quebec. HiFiasm v0.19.5 was used for the initial *de novo* assembly using HiFi PacBio sequences (47). Pore-C reads were mapped to the assembled genome using minimap2, and the resulting BAM files were converted to BED format using the bedtools package)(48). The scaffolding was performed using Salsa with the assembled genome and Pore-C interaction data, with parameters to remove misassembled scaffolds (49) (scripts available at https://github.com/cestari-lab/). We screened the scaffolds for telomeric repeats and evaluated repeat content to identify chromosomes. Telomeric repeats [(TTAGGG)n] were analyzed using Integrated Genome Browser (IGV, Broad Institute) motif tools and JBrowse. To assess repeats, a repeat database was built from the scaffolded genome using RepeatMasker (50). Then, RepeatModeler was used to identify repeat families and create a *de novo* repeat library, followed by RepeatMasker to annotate the assembly. The masked file was used to calculate the percentage of non-masked versus masked nucleotides (the latter annotated as repeats) using a custom script (https://github.com/cestari-lab/) to determine repeat content per scaffold. Finally, the scaffolds were sorted and aligned based on the *T. cruzi* Brazil A4 and Dm25 TcI reference strains (7, 8) using the AssemblyReferenceSorter tool in the Next Generation Sequencing Experience Platform (NGSEP) v4.3.1 (51), resulting in the final set of chromosomes and scaffolds. Synteny analysis was performed to compare the *T. cruzi* Sylvio genome with other TcI strains, including Brazil A4 v.68 (downloaded from TriTrypDB repository), Dm25 (downloaded from NCBI), CL-Brenner Esmeraldo like v.68 (downloaded from TriTrypDB repository), and the previous version of Sylvio X10 v.68 (52). Genomic alignments were generated using minimap2, producing PAF files visualized in JBrowse2 or plotted in R software using the Circlize package to assess structural rearrangements and conserved regions. Genome-wide Average Nucleotide Identity (ANI) was calculated with FastANI to quantify the similarity between genomes. Chromosome-level and scaffold-level were also compared using the same approach.

### Chromatin conformation capture and nanopore sequencing (Pore-C)

*T. cruzi* Sylvio X10 EPs at mid-logarithmic growth (1.8×10^9^ parasites total) were fixed in 1% paraformaldehyde for 10 minutes, rotating at 150 rpm, then quenched with 0.2 M glycine. Cells were centrifuged at 4,000xg and washed three times in PBS. Pellets were resuspended in 1 mL hic-lysis buffer (10 mM Tri-HCl pH 8, 10 M NaCl, 1X Roche protease inhibitor cocktail) and rotated for 30 min at 4°C. They were then pelleted at 2,500xg for 5 minutes at 4°C. The pellet was then resuspended in 0.5% SDS at 62 °C for 10 minutes, followed by adding 1.5% Triton X-100 and incubating at 37 °C for 15 minutes. The sample was digested with 100 units of Hind III HF (New England Biolabs) at 37°C overnight. Samples were incubated for 20 minutes for heat inactivation. DNA ends were filled in with 10mM biotin-dCTP (Jena Bioscience) and non-biotinylated 10 mM dATP, dGTP, and dTTP with 50 units of DNA polymerase I, large Klenow fragment (New England Biolabs) for 16°C at 80 rpm overnight. DNAs were extracted with phenol:chloroform:isoamyl-alcohol (25:24:1, pH 6.7) and precipitated with 100% cold isopropanol for 30 minutes at 14,000 xg at 4°C. The pellet was washed with 75% ethanol and resuspended in 10 mM Tris pH 8. Streptavidin magnetic beads (New England Biolabs) were added and rotated at 150 rpm for 45 minutes at room temperature (RT), then washed three times in 10 mM Tris pH 8 at 55°C for 2 minutes. DNAs were separated using a magnetic rack, eluted in RNA-free water, and incubated with 40 μg of Proteinase K at 55°C overnight and shaking at 150 rpm. DNAs were separated on a magnetic rack and then extracted with phenol:chloroform:isoamyl alcohol (25:24:1, pH 6.7), as above. Eluted DNAs in RNAase free water were used for Oxford nanopore library preparation and DNA sequencing, as previously described (53). The output .fastq files were mapped to the genome using minimap2 (54) and DNA contacts were analyzed using pairtools (55). The scripts used for analysis are available at https://github.com/cestari-lab.

### Chromosomal numbers determination by depth and allelic frequency

The *T. cruzi* Sylvio X10 strain assembled genome (this work) was used as an index for minimap2 alignment with PacBio HiFi reads (54). For ploidy determination, the standardized protocol by Schwabl et al. was followed (56). The Samtools depth tool (v0.1.18) was used to calculate the average read depth for 1 kb bins across each chromosome from the .bam files generated by minimap2. The median of the average depths was then calculated for all bins. Somy was estimated by dividing the average depth by the median at the 60th percentile and multiplying the result by two.

### Genome annotation

The *T. cruzi* Sylvio X10 genome annotation was performed using the GenSAS platform (Fig. S12). Annotation included EST data from *T. cruzi* sequencing libraries available in TrytripDB (https://tritrypdb.org/), companion gene predictions based on the CL-Brenner reference, amino acid FASTA file containing all available *T. cruzi* proteins from TrytripDB, nucleotide FASTA for multigene family proteins from TrytripDB, a GFF file of repeats obtained from RepeatMasker, and paired-end RNA-seq reads from trypomastigotes (sequencing read archive identification: SRR9202394). The reads were mapped to the assembly using HISAT v2.2.1. Initial alignments against the NCBI RefSeq Protozoa nucleotide database were performed using BLASTn v2.12 with specific parameters (Expect value 1e-50, 85% identity, maximum of 10 hits per region, Gap Open of 5, and Gap Extend of 2), along with BLAT v2.5 with a minimum identity of 95%. Alignments were also conducted against the EST evidence and multigene family FASTA files using the same BLASTn and BLAT parameters. PASA v2.4.1 was used to perform spliced alignments of expressed transcript sequences to aid in modeling gene structures. For structural annotation, we used Augustus v3.4.0 for gene prediction, incorporating data from NCBI RefSeq Protozoa and multigene families, supported by gene structures generated through PASA, and protein databases, and RNAseq alignments. Additional ab initio gene prediction was performed using GeneMarkES v4.48, and gene identification was further refined using SNAP v2017-05-17. The results were integrated with EvidenceModeler, which combines ab initio predictions with protein and transcript alignments into a consensus gene structure and refined using PASA. For functional annotation, BLASTp was employed with LOSUM62 matrix, Expect value 1e-8, Word Size 3, Gap Open 11, Gap Extend 1, and a maximum HSP distance of 30,000. DIAMOND v2.0.11 was used to compare against the NCBI RefSeq Protozoa protein dataset and the protein sequences from TrytripDB. InterProScan was run to identify protein signatures from the InterPro consortium, while the Pfam database was used to assign protein families. SignalP predicted the presence and location of signal peptides, and TargetP predicted subcellular localization for eukaryotic proteins. The outputs from the tools were manually inspected and merged to produce the final annotated genome. To annotate subgroups of TSs, annotated TSs were blasted against the TS subgroups from Dm28c genome.

### Proteomic sample preparation

For EPs, a fresh 80 mL culture at a concentration of 1 × 10□ parasites/mL was maintained for 72 hours (logarithmic phase), and the absence of MTs verified under a microscope. The culture was centrifuged at 4000 g for 5 minutes. Parasites were washed twice with 1X PBS, transferred to Eppendorf tubes, centrifuged once at 4000 g for 5 minutes, and the pellet (EPs) collected. MTs, AMs, and CTs were purified as described above (see *T. cruzi and mammalian cell culture*), washed twice in 1X PBS, and pellets collected. Parasite pellets were snap-frozen in liquid nitrogen and stored at -80°C until protein extraction. Protein extraction was performed by resuspending cell pellets in lysis buffer (60 mM Triethylammonium bicarbonate buffer (TAEB), 8 M urea, 1 mM Ethylenediaminetetraacetic acid (EDTA), and 2x protease inhibitors). Samples were sonicated in Covaris M220 ultra sonicator (75 peak power, 10% duty cycle, 200 cycles, and a 100-second duration). Afterward, lysates were transferred to Eppendorf tubes and centrifuged at 10,000 xg for 10 minutes. Extracted proteins (supernatants) were collected and stored at -80°C. Aliquots were collected for SDS-PAGE and proteins quantified using a BCA kit (Life Technologies) according to the manufacturer’s instructions. Four biological replicates were generated for each parasite stage.

### TMT-labeling and mass spectrometry

A 100 μg of extracted parasite proteins (EP, MT, AM, and CT) was adjusted to 100 μL with 100mM TEAB. Samples were reduced with 5 mM TCEP at 55°C for 1 h and alkylated with 18.75 mM iodoacetamide for 30 min at room temperature in the dark. Proteins were precipitated overnight with six volumes of pre-chilled acetone at - 20°C, pelleted by centrifugation, and resuspended in 100mM TEAB pH 8.5. Digestion was performed overnight at 37°C using trypsin at a 1:40 (w/w) enzyme-to-protein ratio. The peptides were treated with TMT-16plex reagents (ThermoFisher Scientific) according to the manufacturer’s instructions. The labeled peptides were fractionated using the Pierce™ High pH Reversed-Phase Peptide Fractionation Kit into 8 fractions. Each fraction was re-solubilized in 0.1% aqueous formic acid, and 2 micrograms of each fraction were loaded onto a Thermo Acclaim Pepmap precolumn (75 μm ID x 2 cm, C18, 3 μm beads) and then onto an Acclaim Pepmap Easyspray analytical column (75 μm x 15 cm, 2 μm C18 beads) for separation using a Dionex Ultimate 3000 uHPLC system at a flow rate of 250 nL/min. A gradient of 2-35% organic solvent (0.1% formic acid in acetonitrile) was applied over three hours, using default settings for MS3-level SPS TMT quantitation (57) on an Orbitrap Fusion mass spectrometer (ThermoFisher Scientific) operating in DDA-MS3 mode. Briefly, MS1 scans were acquired at a resolution of 120,000, scanning from 375-1500 m/z, with ions collected for 50 ms or until an AGC target of 4e5 was reached. Precursors with a charge state of 2-5 were selected for MS2 analysis, using an isolation window of 0.7 m/z. Ions were collected for up to 50 ms or until an AGC target of 1e4 was reached, and fragmented using CID at 35% energy. MS2 spectra were read out in the linear ion trap in rapid mode. From the MS2 spectra, the top 10 precursor notches (based on signal height) were selected for MS3 quantitative TMT reporter ion analysis. These were isolated with a 2 m/z window, fragmented with HCD at 65% energy, and the resulting fragments were read in the Orbitrap at a resolution of 60,000, with a maximum injection time of 105 ms or until an AGC target of 1e5 was reached. Raw .raw files were processed using Proteome Discoverer 2.2 (ThermoFisher Scientific) to identify proteins and quantify TMT reporter ion intensities. Standard TMT quantification workflows were used, with Trypsin set as the enzyme specificity. Spectra were matched against a strain-specific *T. cruzi* database. Dynamic modifications were set as oxidation of methionine (M) and acetylation at protein N-termini. Cysteine carbamidomethylation and TMT tags at both peptide N-termini and lysine residues were set as static modifications. The results were filtered to a 1% FDR to ensure high confidence in protein identification.

### Universal MGF primer design

To evaluate expression changes in MGFs, we designed universal primers to amplify all genes from a gene family. Available sequences of MGFs (Mucins, MASPs, TSs, RHS proteins, DGF-1, and GP63) were obtained from strains Dm28c (2018), Brazil A4, Y C6, CL Brener Esmeraldo-like, CL Brener non-Esmeraldo-like, available in the TriTrypDB database, and Sylvio X10 assembly genome (this work). For each group of MGFs, fasta sequences were aligned using MAFFT, and primers were designed based on conserved regions located at the 3’ conserved ends of the sequences. Due to the high genetic variability within some MGFs, multiple primers were developed to ensure comprehensive coverage. For TSs, primers were designed for each subgroup (I-VIII) using sequences from Dm28c and Sylvio X10 genome (this work) and assessed for compatibility with all annotated TSs in the other genomes. Primer designs were evaluated for secondary structure formation, specificity, and melting temperature (TM) compatibility with *T. cruzi* spliced leader sequences. The final set of primers (Table S2) was tested using DNA from *T. cruzi* EPs to confirm amplification efficiency and reliability.

### MGF-Seq

MTs were used to infect H9-C2 cells (ratio 1:10) and incubated in DMEM medium supplemented with 10% FBS and 1% penicillin and streptomycin at 37°C with 5% CO□. After 10 days, the CTs produced were collected and quantified to re-infect H9-C2 cells, and the process was repeated to produce a total of four consecutive CT generations. After the third generation, CTs were used to infect HEK293T cells. After six days CTs were collected. The procedure was repeated to obtain four biological replicates. MTs and CTs were washed twice in 1X PBS, and RNAs were extracted using the RNeasy® Plus Mini Kit, which removes DNA. cDNAs were synthesized using the LunaScript® RT SuperMix Kit (NEB) according to the manufacturer’s instructions. cDNAs were diluted 10X in water and MGF genes amplified using Taq DNA Polymerase with ThermoPol® Buffer (NEB), according to manufacturer’s protocol, using a splice leader primer and universal 3’ primers for MGF genes (Table S2) with an initial denaturation of 95°C for 5 minutes, and 22 cycles of 95°C for 35 seconds, 59°C for 45 seconds, and 68°C for 2 minutes, and final extension of 68°C for 5 minutes. PCR products were purified using the NucleoMag® NGS Clean-up and Size Select kit (Takara), retaining amplicons larger than 500 bp. DNA fragments were prepared for Oxford Nanopore sequencing using the Ligation Sequencing Kit (SQK-LSK114, Oxford Nanopore Technologies) and the PCR Barcoding Expansion Kit (EXP-PBC096) following the manufacturer’s instructions, with cDNA from biological replicates and infection generations receiving different barcode sequences (26 samples total) and multiplexed. Fifty femtomoles (fmol) of pooled and barcoded libraries were loaded onto a MinION sequencing device with an R10 flow cell (FLO-MIN114, Oxford Nanopore Technologies) and sequenced for 72 hours, yielding ∼15 Gb of data. Total RNA-seq data of CTs were obtained from the Sequence Read Archive (SRA) accession SRR9202394.

### Quantifying *T. cruzi* invasion and infection

A total of 30,000 H9-C2, HEK, HeLa, and 3T3 cells were incubated at 37°C with 5% CO□ in RPMI medium supplemented with 10% FBS and 1% penicillin and streptomycin for 4 hours in 24-well plates to allow adherence. Cells were washed twice with 1X PBS and subsequently infected with Sylvio X10 *T. cruzi* MT or CT at a parasite-to-host cell ratio of 1:10. After 3 hours of incubation, the cells were washed to remove unbound parasites from the supernatant and fixed using 4% formaldehyde. Cells were mounted using fluoromount with DAPI. Stained DNAs of mammalian cells and parasites were used to quantify infections using a Citation 5 Imaging System under a 40x objective.

### YSD screen using Chagas disease patient antibodies

A genome-wide yeast surface display was constructed for *T. cruzi* Sylvio X10 and transformed into EBY100 *Saccharomyces cerevisiae* (33, 53). To express *T. cruzi* proteins on the yeast surface, 1.2×10^7^ yeast cells were grown in 5 ml YPD media at 30°C and 225 rpm until OD600 1 (∼2 hours). Cells were collected by centrifugation at 3000 xg for 5 minutes and washed three times in 10 ml sterile MilliQ water by centrifugation at 3000 xg for 5 minutes. Cells were resuspended in 50ml SD/-trp media with 2% dextrose (SD/-trp+dex) and incubated at 30°C and 225 rpm until OD600 reaches 1 (∼16h). Cells were collected by centrifugation and washed in water, as indicated above. Cells were resuspended in 50 ml SD/-trp with 2% raffinose and incubated at 30°C and 225 rpm for 2 h. Cells were collected by centrifugation, resuspended in 50 ml SD/-trp + 2% raffinose + 1% galactose for protein expression, and incubated at 30°C and 225 rpm for 16h. For binding assays, yeasts were collected by centrifugation and washed three times in ice-cold PBS (as indicated above) and incubated in 1:1000 pooled sera of five chronic-stage Chagas disease patients diluted in PBS or 1:1000 pooled sera of two healthy individuals. Sera was kindly provided by Dr. Momar Ndao (McGill University Health Centre). Yeast and sera mix were incubated, rotating for 2 hours at 4°C. Cells were collected by centrifugation and washed three times in PBS. Cells were resuspended in 400 μL of 10 mM PBS and 100 μL magnetic Protein G beads and incubated at 4°C rotating for 30 minutes. The 500 μL yeast and magnetic bead mixture was loaded onto a Miltenyi Biotech column and passed through by gravity on a magnetic stand. The bound yeasts were washed in the column three times in PBS to remove unspecific binders. Antibody-bound yeasts were removed from the magnetic rack and re-cultured to expand the enriched (binders) population in SD/-trp + 1% dextrose overnight. This process was repeated three times for both conditions to enrich antibody-binding populations. To identify antibody-binding proteins, the yeast cells from expanded populations were collected and washed three times to remove media and plasmid DNAs were extracted using an adapted protocol from (58), where DNA is purified using NGS magnetic beads at 0.7x ratio rather than isopropanol precipitation. Oxford nanopore library and sequencing were prepared as previously described (59). Scripts used for computational analysis can be found at https://github.com/cestari-lab/YSD. Briefly, Oxford nanopore sequence .fasta files were aligned to the genome using minimap2. Sequence Alignment Map (.sam) files were converted to Binary Alignment Map (.bam) files using the samtools package, filtering supplementary and secondary reads and any reads with a MAPQ score < 1. The coverage was analyzed using plotCoverage from deeptools. The resulting .bed file was visualized using Integrative Genomic Visualizer (IGV, Broad Institute). Read counts were obtained using featureCounts, and fold change comparisons between groups were performed using edgeR and enriched regions with macs3. Heatmaps were generated from edgeR enrichment analysis.

### Data Sharing

The assembled genome of the *T. cruzi* Sylvio X10 strain is available in the National Center for Biotechnology Information (NCBI) with BioProject number PRJNA1237338 and at the DNA DataBank of Japan (DDBJ), the European Nucleotide Archive (ENA), and GenBank at NCBI under the accession JBMETK000000000. PacBio HiFi sequences and Pore-C data are available in the Sequence Read Archive (SRA) with the BioProject identification PRJNA1236874. The mass spectrometry proteomics data have been deposited to the ProteomeXchange Consortium via the PRIDE partner repository with the dataset identifier PXD061891. Codes used for data analysis are available at https://github.com/cestari-lab/.

## Author Contributions

LBA and ML extracted *T. cruzi* DNA for genome sequencing. LCS designed a genome assembly pipeline and performed computational analysis. LCS performed *T. cruzi* infections, purifications, and sample preparation for mass spectrometry. LCS analyzed mass spectrometry data. IC and LCS designed MGF-seq, and LCS performed MGF-seq and computational analysis. ML and LBA transformed *T. cruzi* genome-wide in yeast, and ML performed the YSD screen and computational analysis. IC designed the research, analyzed the results, secured research funding, supervised the study, and wrote the manuscript. All authors read and revised the manuscript.

## Acknowledgments

This research was partly enabled by computational resources provided by Calcul Quebec (https://www.calculquebec.ca/en/) and the Digital Research Alliance of Canada (alliancecan.ca).

## Supplementary figures legends

**Figure S1. Chromosomal depth and repeat content in *T. cruzi* TcI strain Sylvio X10**. A) The circos plot displays chromosomal depth calculated in 10,000 bp windows. Chromosome 30 shows an increase in depth, consistent with a trisomy structure, while chromosome 1 and both haplotypes of chromosome 22 exhibit decreased depth. Chromosomes 10 and 17 show evidence of segmental aneuploidy, indicated by a decrease in depth in the initial regions of these chromosomes. B) The percentage of repeat content is shown across chromosomes, with high repeat content (>70%) observed on chromosomes 9, 12, 29 and 30 (indicated with asterisks), which are enriched in multigene family (MGF) sequences.

**Figure S2. Synteny analysis of *T. cruzi* scaffolds and chromosomes**. Comparison of assembled short-length scaffolds (<60 kb) reveals segmental aneuploidy and/or haplotype differences. The chromosomes are ordered from the highest to the lowest number of scaffold matches. Chromosome 16 shows synteny with eight scaffolds, chromosome 30 with six scaffolds, and chromosome 26 with five scaffolds. All 39 scaffolds display synteny with some chromosomes.

**Figure S3. Mitochondrial genome structure**. A) The circos plot displays a map of the assembled *T. cruzi* mitochondrial maxicircles, highlighting the locations of unedited genes, long repeat regions, and short repeat regions.

**Figure S4. Synteny between regions of *T. cruzi* genomes**. The synteny graph displays a synteny comparison between the genomes of *T. cruzi* strain X10 (this work) and the genomes of the strains DM25, Brazil A4, Sylvio X10-2018 assembly, and CL-Brener Esmeraldo-like.

**Figure S5. Differentially expressed proteins in the *T. cruzi* life cycle**. A. Comparison between protein expression changes in non-replicative and replicative stages. MT vs AM, MT vs EP, CT vs AM, and CT vs EP. The top 10 differentially expressed proteins are listed. B) Comparison between CT vs MT. C) Comparison between EP vs AP. Blue bars indicate the top 10 down-regulated proteins, while red bars represent the top 10 up-regulated proteins.

**Figure S6. Protein expression pattern of MGFs across *T. cruzi* life stages**. Cumulative abundance of all dispersed gene family 1 (DGF-1), GP63, mucin-associated surface protein (MASP), mucin, retrotransposon hotspot protein (RHS), and trans-sialidases. Trans-sialidases are organized into their respective groups I to VIII and the unclassified group. TcMUCII is a mucin group. The abundance of all proteins within a group was summed to represent the cumulative abundance.

**Figure S7. Distribution and expression of MGFs and housekeeping (Other Genes) proteins in all chromosomes across *T. cruzi* life stages**. The X-axis represents chromosome length in base pairs, while the Y-axis shows the log□protein abundance, calculated as the mean of four biological replicates. The top plot corresponds to MGF and the bottom plot to other genes. Chromosomes enriched in multigene families (MGFs) —specifically chromosomes 6, 9, 29, and 31 — show increased MGF expression in CTs compared to other life cycle stages, as well as compared to non-MGF proteins.

**Figure S8. Transcriptomic and proteomic analysis of MGF gene expression in *T. cruzi***. RNA-seq and proteomic data confirm MGF gene expression across multiple chromosomes. The circos plot displays all 31 chromosomes, showing protein and RNA abundance by genomic coordinates. Red dots represent the log2 abundance of MGF genes, while black dots indicate non-MGF genes, i.e., housekeeping genes (HK).

**Figure S9. Distribution of protein and RNA expression of MGF and housekeeping genes**. The density plots show the distribution of log2-transformed expression of MASP, mucin, RHS, DGF-1, GP63, and housekeeping genes in CTs.

**Figure S10. MGF expression across multiple rounds of cell infection**. A) Volcano plots show differentially expressed transcripts across multiple generations of CTs infections in H9-C2 or CTs from infection in HEK293T cells. The X-axis represents log□fold change, while the Y-axis represents -log□□(p-value). Transcripts showing significant expression (log□ ≥ 1, *p*-value ≤ .05) are highlighted in red. B) The bar plot displays the most abundant MGF transcripts across all samples. Their expression is predominantly observed in G4 of CTs from H9-C2 or CTs derived from HEK293T cells. C) The graph represents the number of MGF transcripts expressed per generation after each round of infection. The X-axis represents experimental groups (G0–G4), while the Y-axis indicates the number of MGF genes expressed. G0 is MT, G1-G4 are CTs.

**Figure S11. Sequence alignment of the N-terminus of 30 TSs recognized by Chagas disease patients’ antibodies**. Alignment of the amino acids 30-100 of 30 TS proteins reveals significant sequence divergence and a few conserved amino acids. Conserved amino acids are highlighted in color. The first 30 amino acids were removed to eliminate signal peptides.

**Figure S12. Annotation pipeline used in the *T. cruzi* Sylvio X10 strain genome**. The *T. cruzi* Sylvio X10 genome was annotated using GenSAS, which integrates expressed sequence tags (ESTs), protein and nucleotide sequences, and RepeatMasker data. RNA-seq reads (Sequence Read Archive accession SRR9202394) were also utilized. Reads were mapped with HISAT2, and alignments were performed using BLASTn and BLAT. Structural annotation combined Augustus, PASA, GeneMarkES, and SNAP, with EvidenceModeler integrating predictions. Functional annotation was performed using BLASTp, DIAMOND, InterProScan, and Pfam, while SignalP and TargetP predicted protein features. TS subgroups were classified using BLAST against *T. cruzi* Dm28c strain. The final annotation was manually curated.

**Table S1. Quantitative proteomics of *T. cruzi* life cycle stages**. The data show abundance and differential expression among the four stages (EP, MT, AM, CT).

**Table S2. Primers designed for nanopore multigene family sequencing**. SL, splice leader sequence. Primers were designed to amplify all annotated MGF sequences.

## References

1. S. K. McKenzie, D. J. C. Kronauer, The genomic architecture and molecular evolution of ant odorant receptors. Genome Res 28, 1757–1765 (2018).

2. R. Pushker, A. Mira, F. Rodriguez-Valera, Comparative genomics of gene-family size in closely related bacteria. Genome Biol 5, R27 (2004).

3. K. W. Deitsch, S. A. Lukehart, J. R. Stringer, Common strategies for antigenic variation by bacterial, fungal and protozoan pathogens. Nat Rev Microbiol 7, 493–503 (2009).

4. S. B. Cannon, A. Mitra, A. Baumgarten, N. D. Young, G. May, The roles of segmental and tandem gene duplication in the evolution of large gene families in Arabidopsis thaliana. BMC Plant Biol 4, 10 (2004).

5. V. U. Schwartze et al., Gene expansion shapes genome architecture in the human pathogen Lichtheimia corymbifera: an evolutionary genomics analysis in the ancient terrestrial mucorales (Mucoromycotina). PLoS Genet 10, e1004496 (2014).

6. L. Berna et al., Expanding an expanded genome: long-read sequencing of Trypanosoma cruzi. Microb Genom 4 (2018).

7. M. C. Hoyos Sanchez et al., A phased genome assembly of a Colombian Trypanosoma cruzi TcI strain and the evolution of gene families. Sci Rep 14, 2054 (2024).

8. W. Wang et al., Strain-specific genome evolution in Trypanosoma cruzi, the agent of Chagas disease. PLoS Pathog 17, e1009254 (2021).

9. L. M. De Pablos, A. Osuna, Multigene families in Trypanosoma cruzi and their role in infectivity. Infect Immun 80, 2258–2264 (2012).

10. P. J. Hotez et al., An unfolding tragedy of Chagas disease in North America. PLoS Negl Trop Dis 7, e2300 (2013).

11. I. C. Almeida, M. A. Ferguson, S. Schenkman, L. R. Travassos, Lytic anti-alpha-galactosyl antibodies from patients with chronic Chagas’ disease recognize novel O-linked oligosaccharides on mucin-like glycosyl-phosphatidylinositol-anchored glycoproteins of Trypanosoma cruzi. Biochem J 304 (Pt 3), 793–802 (1994).

12. R. L. Tarleton, CD8+ T cells in Trypanosoma cruzi infection. Semin Immunopathol 37, 233–238 (2015).

13. R. L. Tarleton, B. H. Koller, A. Latour, M. Postan, Susceptibility of beta 2-microglobulin-deficient mice to Trypanosoma cruzi infection. Nature 356, 338–340 (1992).

14. G. J. Medina-Rincon et al., Molecular and Clinical Aspects of Chronic Manifestations in Chagas Disease: A State-of-the-Art Review. Pathogens 10 (2021).

15. F. Altamura, R. Rajesh, C. M. C. Catta-Preta, N. S. Moretti, I. Cestari, The current drug discovery landscape for trypanosomiasis and leishmaniasis: Challenges and strategies to identify drug targets. Drug Dev Res 83, 225–252 (2022).

16. E. S. Nakayasu et al., GPIomics: global analysis of glycosylphosphatidylinositol-anchored molecules of Trypanosoma cruzi. Mol Syst Biol 5, 261 (2009).

17. G. A. Burle-Caldas et al., Disruption of Active Trans-Sialidase Genes Impairs Egress from Mammalian Host Cells and Generates Highly Attenuated Trypanosoma cruzi Parasites. mBio 13, e0347821 (2022).

18. C. A. Buscaglia et al., The surface coat of the mammal-dwelling infective trypomastigote stage of Trypanosoma cruzi is formed by highly diverse immunogenic mucins. J Biol Chem 279, 15860–15869 (2004).

19. M. H. Magdesian et al., Infection by Trypanosoma cruzi. Identification of a parasite ligand and its host cell receptor. J Biol Chem 276, 19382–19389 (2001).

20. G. D. Pollevick et al., Trypanosoma cruzi surface mucins with exposed variant epitopes. J Biol Chem 275, 27671–27680 (2000).

21. F. R. Santori et al., Identification of a domain of Trypanosoma cruzi metacyclic trypomastigote surface molecule gp82 required for attachment and invasion of mammalian cells. Mol Biochem Parasitol 78, 209–216 (1996).

22. S. Schenkman, M. S. Jiang, G. W. Hart, V. Nussenzweig, A novel cell surface trans-sialidase of Trypanosoma cruzi generates a stage-specific epitope required for invasion of mammalian cells. Cell 65, 1117–1125 (1991).

23. S. Schenkman, L. Pontes de Carvalho, V. Nussenzweig, Trypanosoma cruzi trans-sialidase and neuraminidase activities can be mediated by the same enzymes. J Exp Med 175, 567–575 (1992).

24. S. Tomlinson, L. C. Pontes de Carvalho, F. Vandekerckhove, V. Nussenzweig, Role of sialic acid in the resistance of Trypanosoma cruzi trypomastigotes to complement. J Immunol 153, 3141–3147 (1994).

25. J. P. F. Rodrigues et al., Host cell protein LAMP-2 is the receptor for Trypanosoma cruzi surface molecule gp82 that mediates invasion. Cell Microbiol 21, e13003 (2019).

26. T. S. Onofre, L. Loch, J. P. Ferreira Rodrigues, S. Macedo, N. Yoshida, Gp35/50 mucin molecules of Trypanosoma cruzi metacyclic forms that mediate host cell invasion interact with annexin A2. PLoS Negl Trop Dis 16, e0010788 (2022).

27. M. V. Chuenkova, M. PereiraPerrin, Trypanosoma cruzi targets Akt in host cells as an intracellular antiapoptotic strategy. Sci Signal 2, ra74 (2009).

28. J. Mucci et al., Thymocyte depletion in Trypanosoma cruzi infection is mediated by trans-sialidase-induced apoptosis on nurse cells complex. Proc Natl Acad Sci U S A 99, 3896–3901 (2002).

29. M. G. Risso, T. A. Pitcovsky, R. L. Caccuri, O. Campetella, M. S. Leguizamon, Immune system pathogenesis is prevented by the neutralization of the systemic trans-sialidase from Trypanosoma cruzi during severe infections. Parasitology 134, 503–510 (2007).

30. C. Poveda et al., Cytokine profiling in Chagas disease: towards understanding the association with infecting Trypanosoma cruzi discrete typing units (a BENEFIT TRIAL sub-study). PLoS One 9, e91154 (2014).

31. S. M. Teixeira, D. G. Russell, L. V. Kirchhoff, J. E. Donelson, A differentially expressed gene family encoding “amastin,” a surface protein of Trypanosoma cruzi amastigotes. J Biol Chem 269, 20509–20516 (1994).

32. M. J. Pinazo et al., Efficacy and safety of fexinidazole for treatment of chronic indeterminate Chagas disease (FEXI-12): a multicentre, randomised, double-blind, phase 2 trial. Lancet Infect Dis 24, 395–403 (2024).

33. R. Heslop et al., Genome-Wide Libraries for Protozoan Pathogen Drug Target Screening Using Yeast Surface Display. ACS Infect Dis 9, 1078–1091 (2023).

34. E. T. Boder, K. D. Wittrup, Yeast surface display for screening combinatorial polypeptide libraries. Nat Biotechnol 15, 553–557 (1997).

35. J. L. Reis-Cunha et al., Accessing the Variability of Multicopy Genes in Complex Genomes using Unassembled Next-Generation Sequencing Reads: The Case of Trypanosoma cruzi Multigene Families. mBio 13, e0231922 (2022).

36. L. M. Freitas et al., Genomic analyses, gene expression and antigenic profile of the trans-sialidase superfamily of Trypanosoma cruzi reveal an undetected level of complexity. PLoS One 6, e25914 (2011).

37. A. V. Jager, R. P. Muia, O. Campetella, Stage-specific expression of Trypanosoma cruzi trans-sialidase involves highly conserved 3’ untranslated regions. FEMS Microbiol Lett 283, 182–188 (2008).

38. Z. H. Li et al., A 43-nucleotide U-rich element in 3’-untranslated region of large number of Trypanosoma cruzi transcripts is important for mRNA abundance in intracellular amastigotes. J Biol Chem 287, 19058–19069 (2012).

39. A. Tsai, M. R. Alves, J. Crocker, Multi-enhancer transcriptional hubs confer phenotypic robustness. Elife 8 (2019).

40. R. Laidlaw et al., The Trypanosoma cruzi cell atlas; a single-cell resource for understanding parasite population heterogeneity and differentiation. BioRxiv 10.1101/2024.10.01.616042 (2024).

41. C. S. Rosenberg, W. Zhang, J. M. Bustamante, R. L. Tarleton, Long-Term Immunity to Trypanosoma cruzi in the Absence of Immunodominant trans-Sialidase-Specific CD8+ T Cells. Infect Immun 84, 2627–2638 (2016).

42. F. Ferragut, G. R. Acevedo, K. A. Gomez, T Cell Specificity: A Great Challenge in Chagas Disease. Front Immunol 12, 674078 (2021).

43. G. R. Acevedo et al., In Silico Guided Discovery of Novel Class I and II Trypanosoma cruzi Epitopes Recognized by T Cells from Chagas’ Disease Patients. J Immunol 204, 1571–1581 (2020).

44. I. Cestari, K. Stuart, Transcriptional Regulation of Telomeric Expression Sites and Antigenic Variation in Trypanosomes. Curr Genomics 19, 119–132 (2018).

45. M. Sadigursky, C. I. Brodskyn, A new liquid medium without blood and serum for culture of hemoflagellates. Am J Trop Med Hyg 35, 942–944 (1986).

46. L. Cruz-Saavedra et al., Purification of Trypanosoma cruzi metacyclic trypomastigotes by ion exchange chromatography in sepharose-DEAE, a novel methodology for host-pathogen interaction studies. J Microbiol Methods 142, 27–32 (2017).

47. H. Cheng, G. T. Concepcion, X. Feng, H. Zhang, H. Li, Haplotype-resolved de novo assembly using phased assembly graphs with hifiasm. Nat Methods 18, 170–175 (2021).

48. A. R. Quinlan, I. M. Hall, BEDTools: a flexible suite of utilities for comparing genomic features. Bioinformatics 26, 841–842 (2010).

49. J. Ghurye, M. Pop, S. Koren, D. Bickhart, C. S. Chin, Scaffolding of long read assemblies using long range contact information. BMC Genomics 18, 527 (2017).

50. M. Tarailo-Graovac, N. Chen, Using RepeatMasker to identify repetitive elements in genomic sequences. Curr Protoc Bioinformatics Chapter 4, 4 10 11–14 10 14 (2009).

51. D. Tello et al., NGSEP3: accurate variant calling across species and sequencing protocols. Bioinformatics 35, 4716–4723 (2019).

52. J. L. Reis-Cunha et al., Whole genome sequencing of Trypanosoma cruzi field isolates reveals extensive genomic variability and complex aneuploidy patterns within TcII DTU. BMC Genomics 19, 816 (2018).

53. T. Sternlieb, M. Loock, M. Gao, I. Cestari, Efficient Generation of Genome-wide Libraries for Protein-ligand Screens Using Gibson Assembly. Bio Protoc 12 (2022).

54. H. Li, Minimap2: pairwise alignment for nucleotide sequences. Bioinformatics 34, 3094–3100 (2018).

55. Open2C et al., Pairtools: From sequencing data to chromosome contacts. PLoS Comput Biol 20, e1012164 (2024).

56. P. Schwabl et al., Meiotic sex in Chagas disease parasite Trypanosoma cruzi. Nat Commun 10, 3972 (2019).

57. G. C. McAlister et al., MultiNotch MS3 enables accurate, sensitive, and multiplexed detection of differential expression across cancer cell line proteomes. Anal Chem 86, 7150–7158 (2014).

58. J. Sambrook, D. W. Russell, Molecular Cloning: A Laboratory Manual (Cold Spring Harbor Laboratory Press, New York, ed. 3rd, 2001), vol. Vol. 1, pp. 628.

59. T. Sternlieb, M. Loock, M. Gao, I. Cestari, Efficient Generation of Genome-wide Libraries for Protein-ligand Screens Using Gibson Assembly. Bio Protoc 13 (2023).

