## Supplementary material for "Stochastic variation in surface protein expression diversifies *Trypanosoma cruzi* infection": Table S2

**Table 2. Primers designed for nanopore multigene family sequencing.** SL, splice leader sequence. Primers were designed to amplify all annotated MGF sequences.

| **Primer name** | **Sequence** |
| --- | --- |
| SL _T. cruzi | AACGCTATTATTGATACAGTTTC |
| Actin_long | GACTGTTCTTCGTCAGACAT |
| β-Tubulin_long | GAGGAGGAGCAGTACTAG |
| α-Tubulin_long | GATGTGGAGGAGTACTAG |
| GAPDH_long | GTTCGGCAAGGTTGTAG |
| Mucin | CAGCCTCAGCAGCTCTG |
| Mucin-2 | AAGCACCAACAGGGCG |
| TSI | AATGCCGAGGAGATCAAGACCTT |
| TSII | TTTCTGTACAACCGCCCACT |
| TSIII | ATGGCTCTAATTGGTGACAGCA |
| TSIV-V-VI | TTGGGACTGTGGGGGTTTG |
| TSVII-VIII | ATCCACGAGGTGCCGAA |
| TS unclassified | CGCGGAAGTAAACA |
| MASP | ACAGTGACGGCAGCACC |
| MASP-2 | TGCAGCCACCCACTG |
| RHS | TTCGTACCTCCTCTAC |
| RHS_2 | ACTCGCATTGTACACAG |
| DGF | TGCTGCGCGATGACGA |
| DGF-2 | AACCGTGATGGCTCTG |
| GP63-1 | GCAGTTTGACAGCTGCA |
| GP63-2 | TCCGACCGCCGTCA |
| GP63-3 | CGTCGGACCGGTATTC |
| GP63-4 | CTGGACCACTGCTGCC |
