## Supplementary figures and images for "Stochastic variation in surface protein expression diversifies *Trypanosoma cruzi* infection"

### Fig. S1

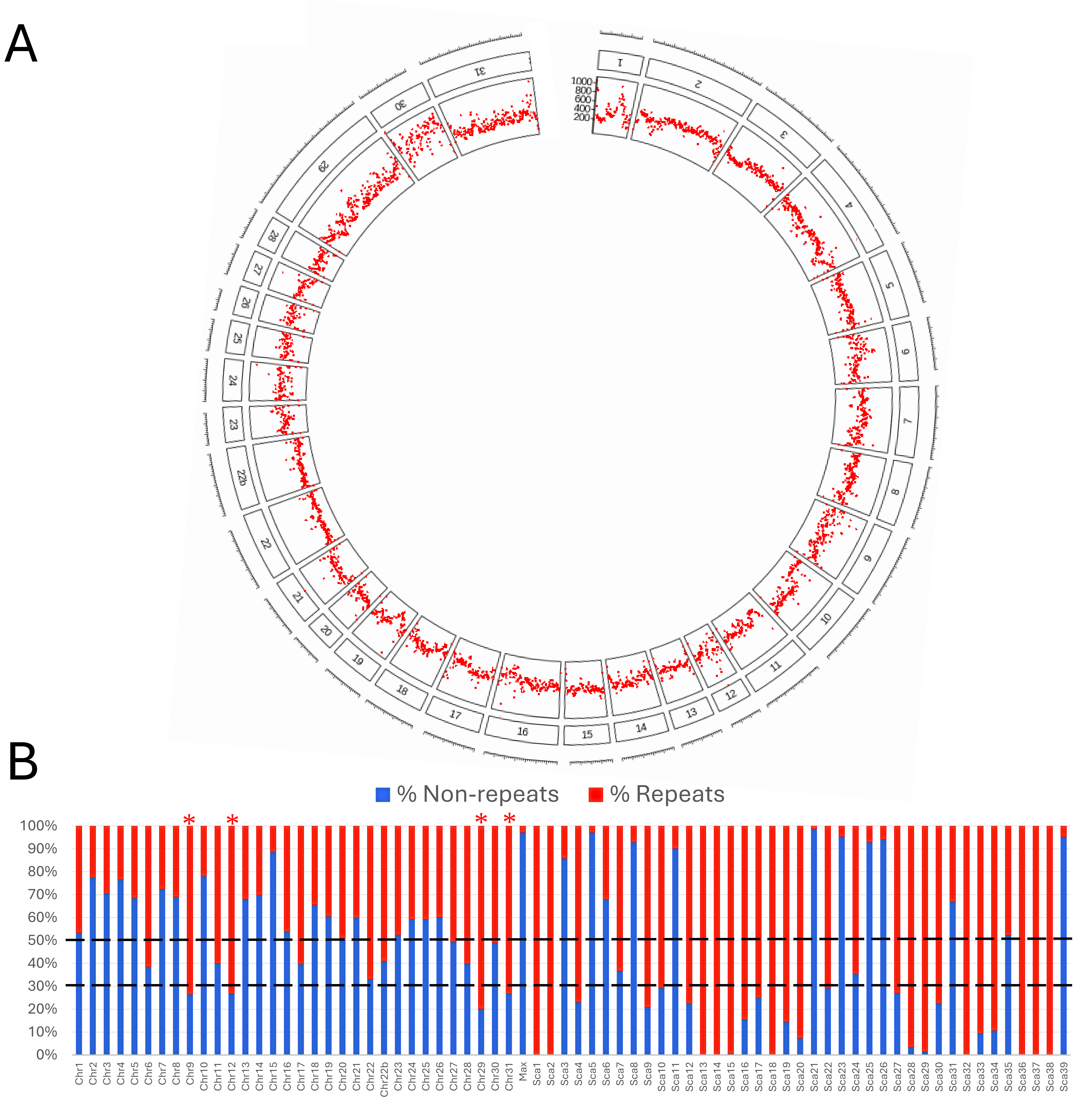

### Fig. S3

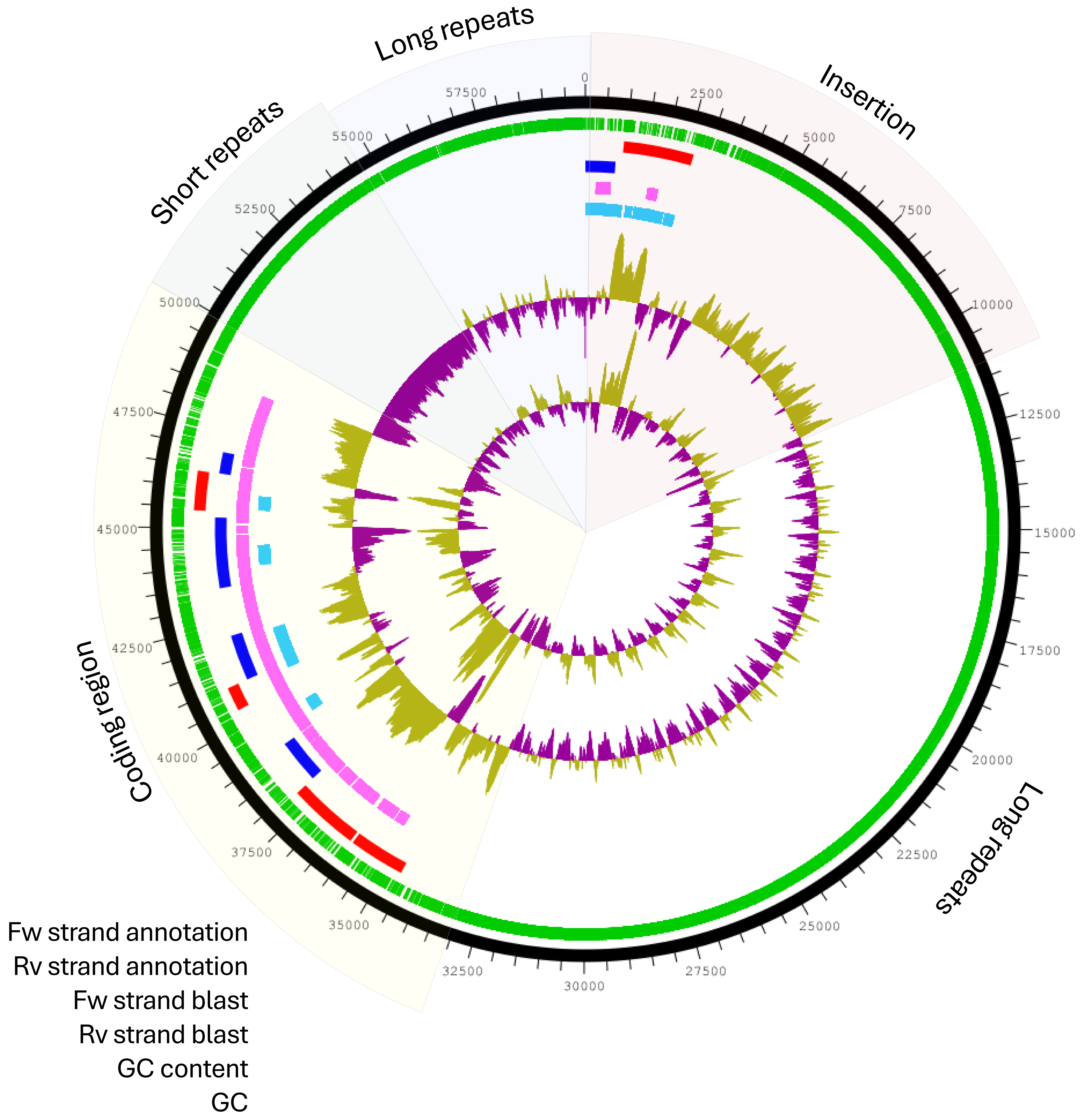

### Fig. S5

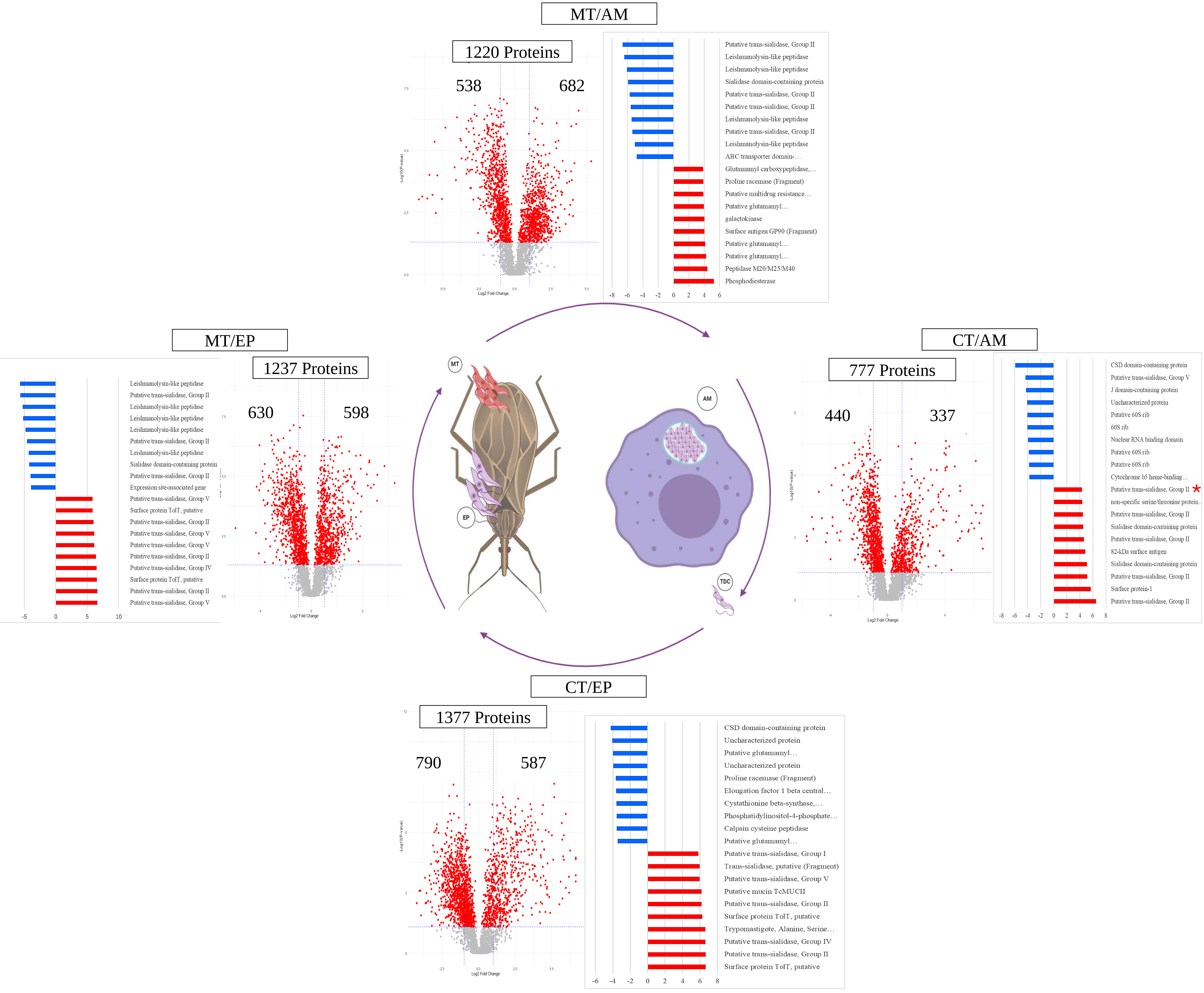

B

#### Trypomastigotes Infective stages

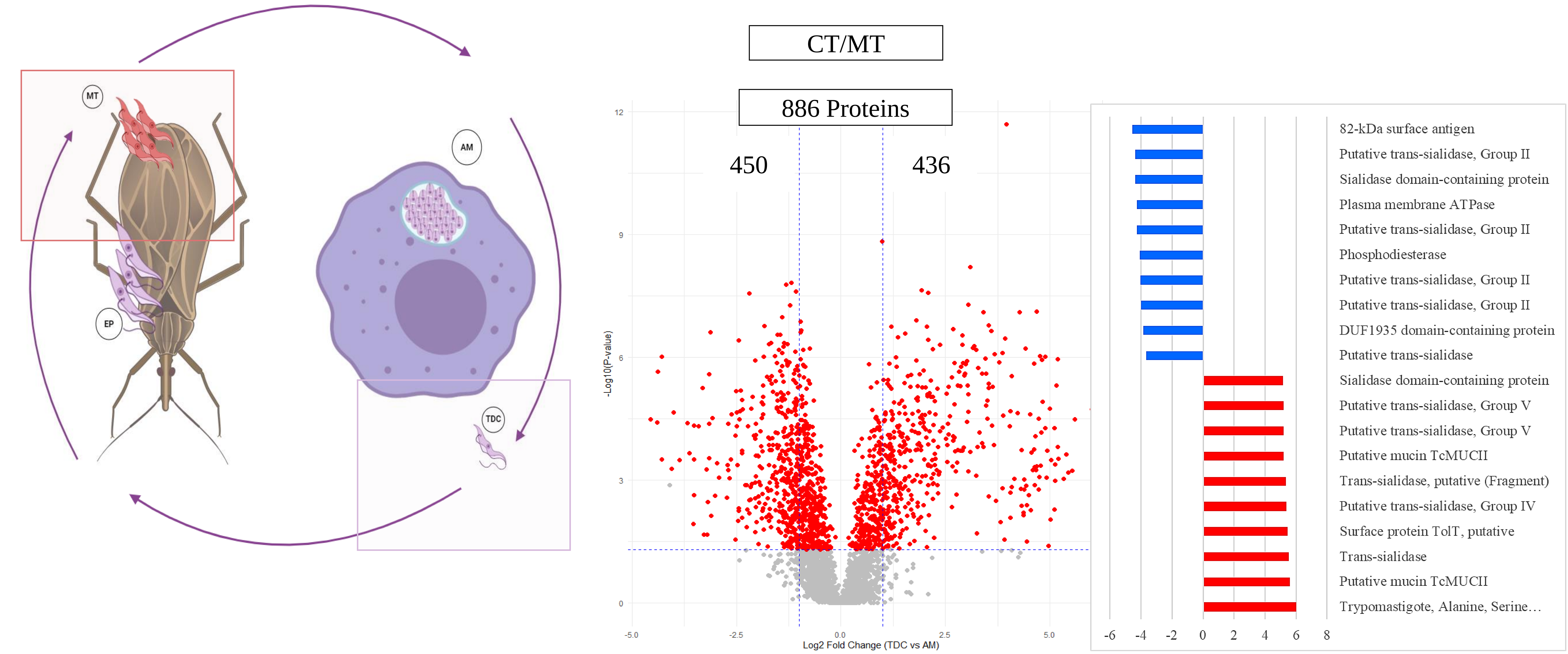


C

#### Replicative stages

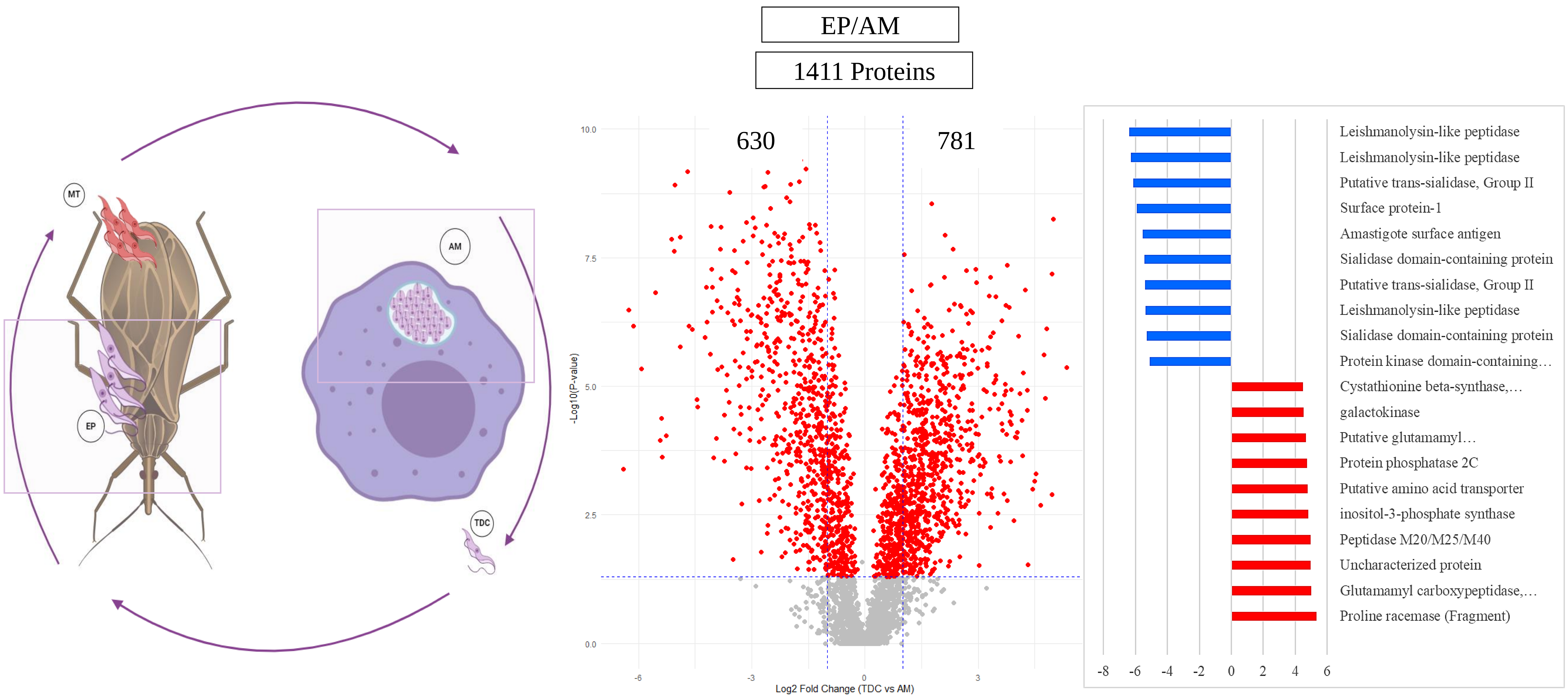

### Fig. S6

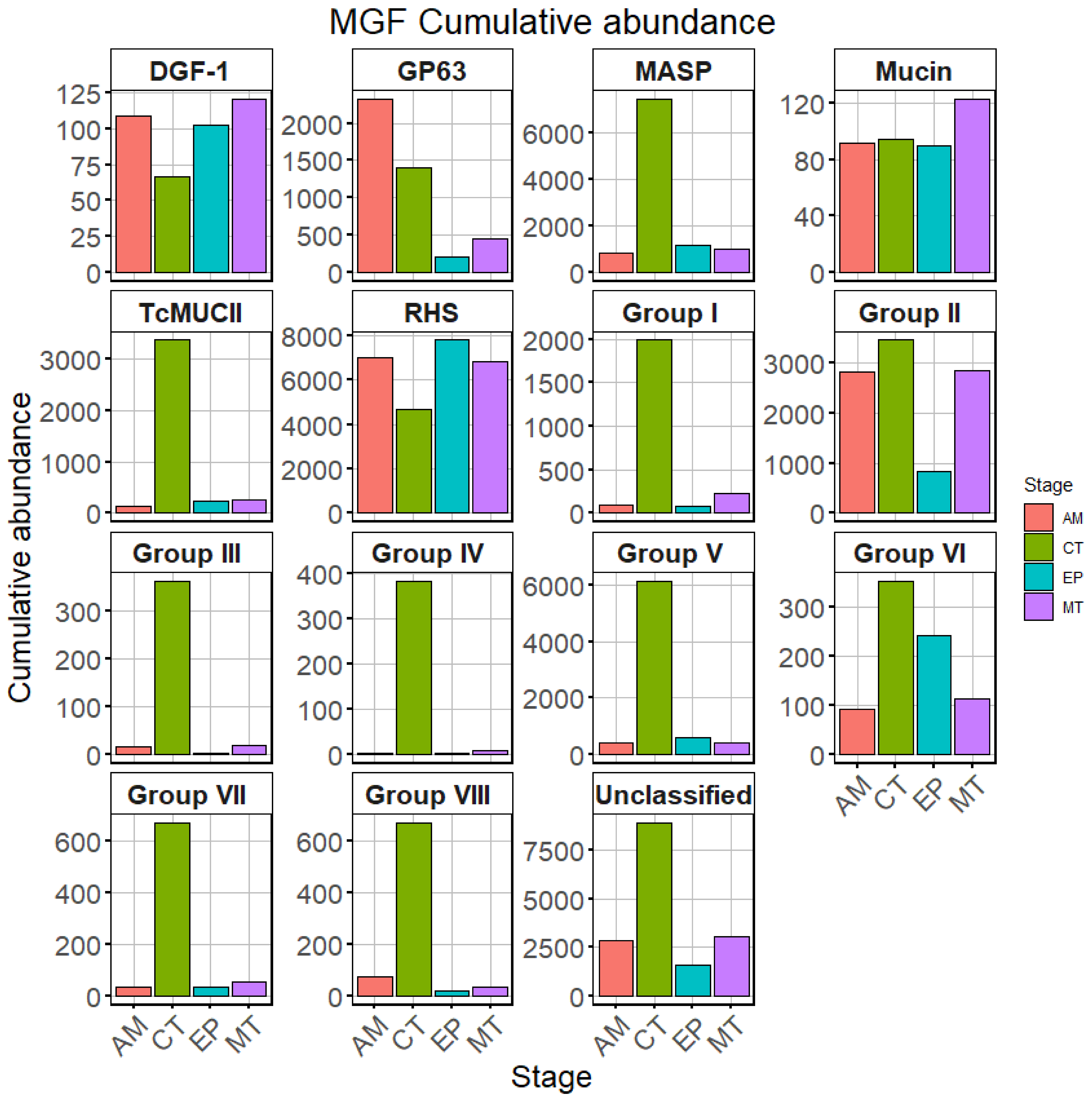

### Fig. S8

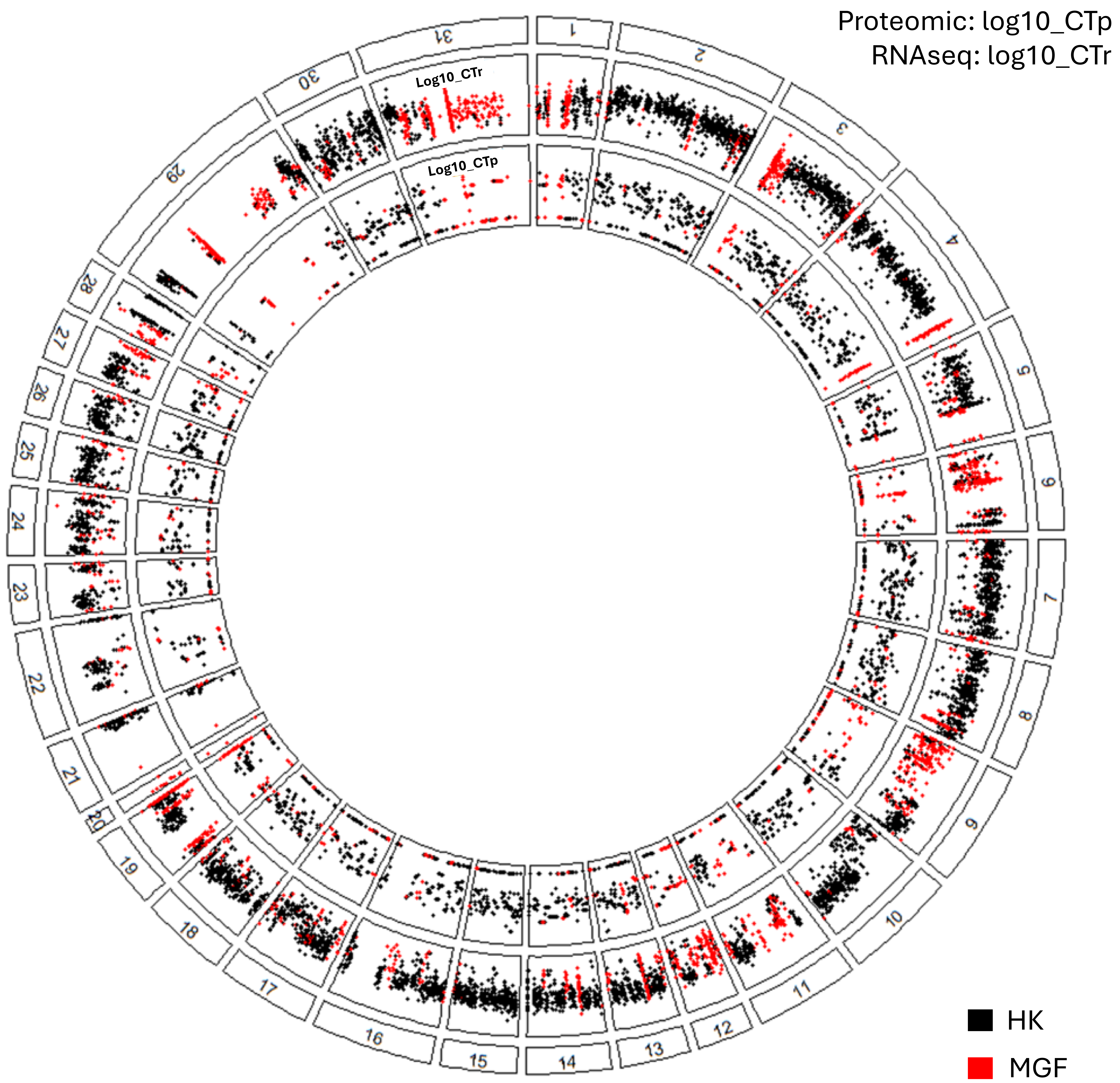

### Fig. S9

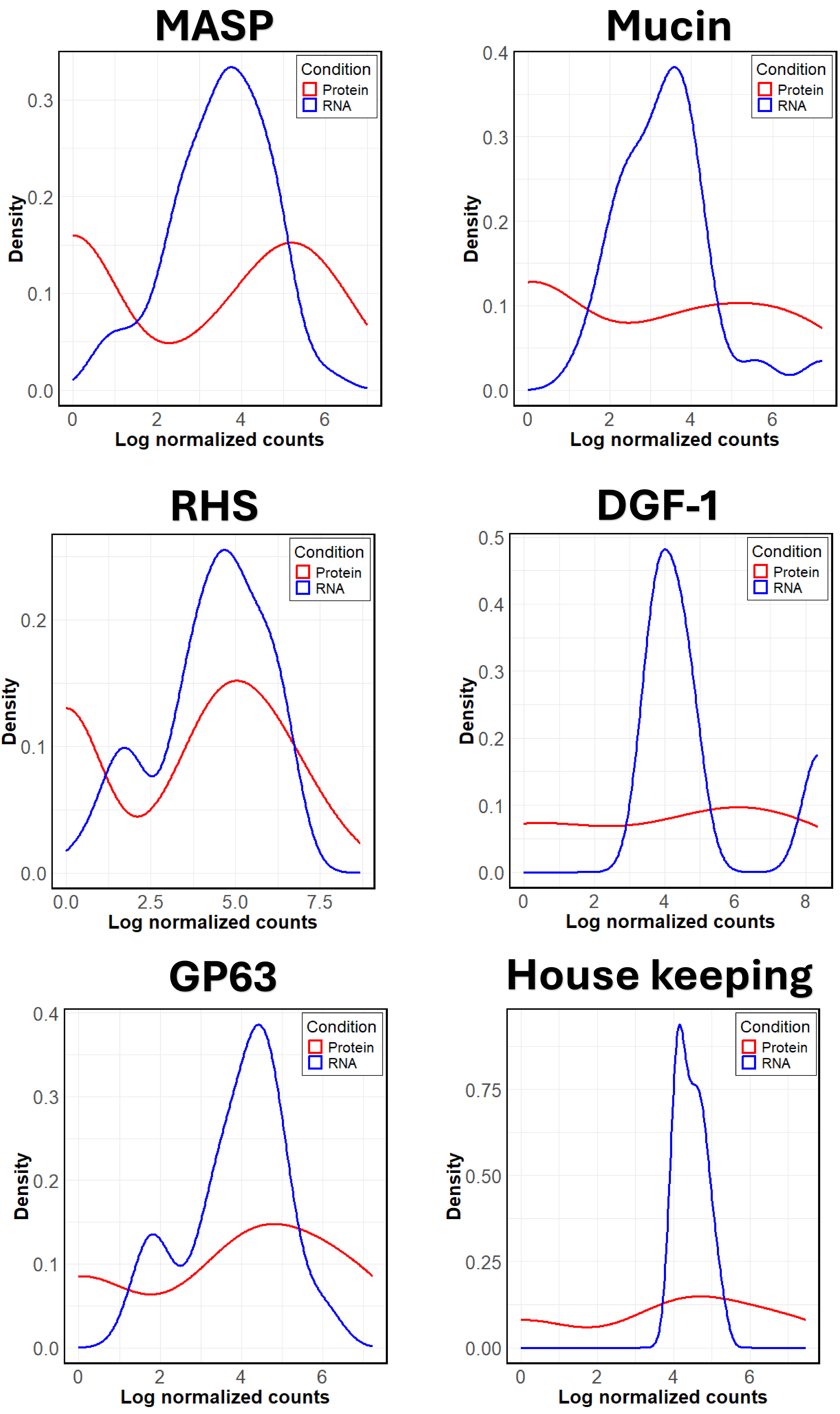

### Fig. S10

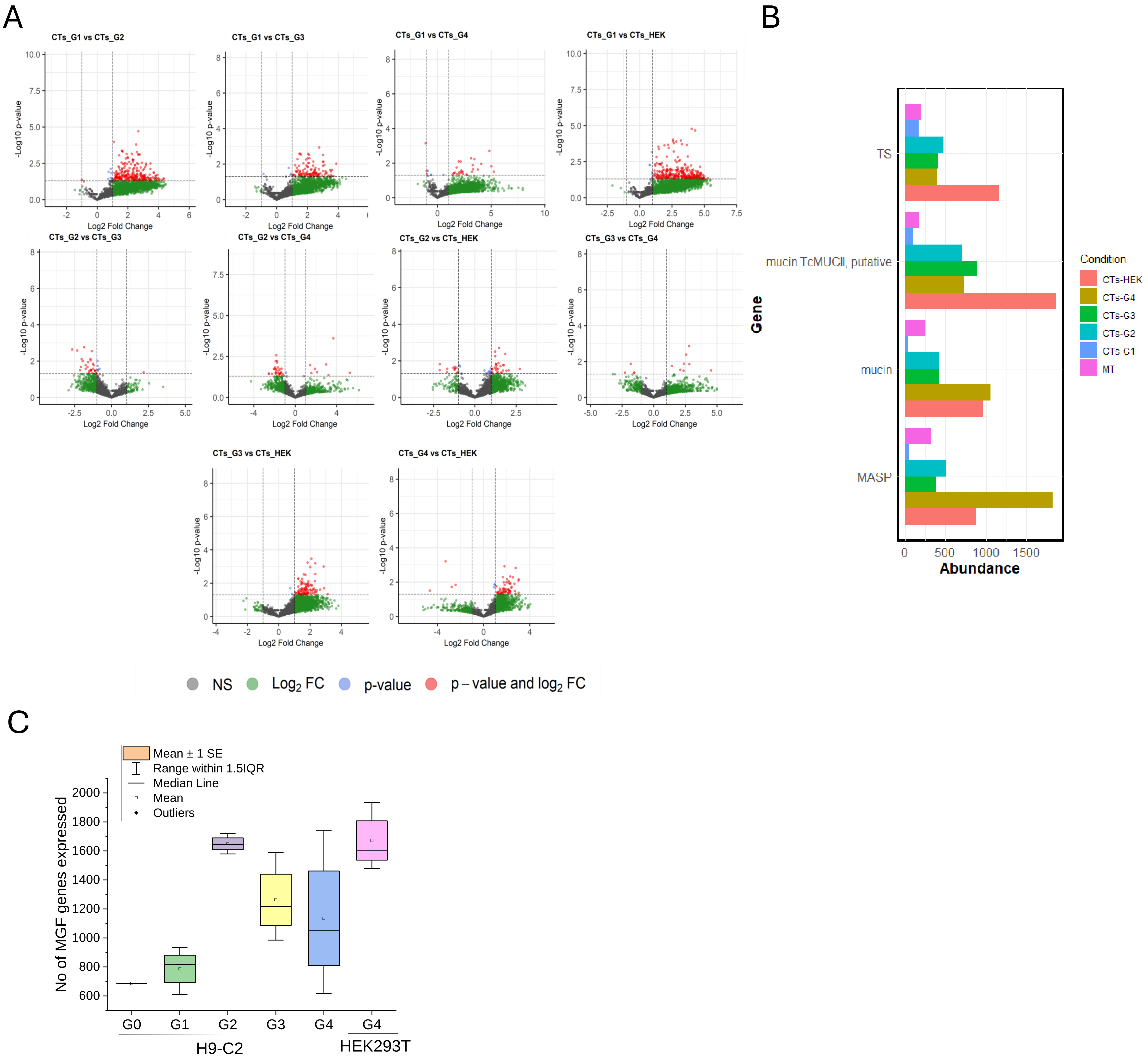

### Fig. S11

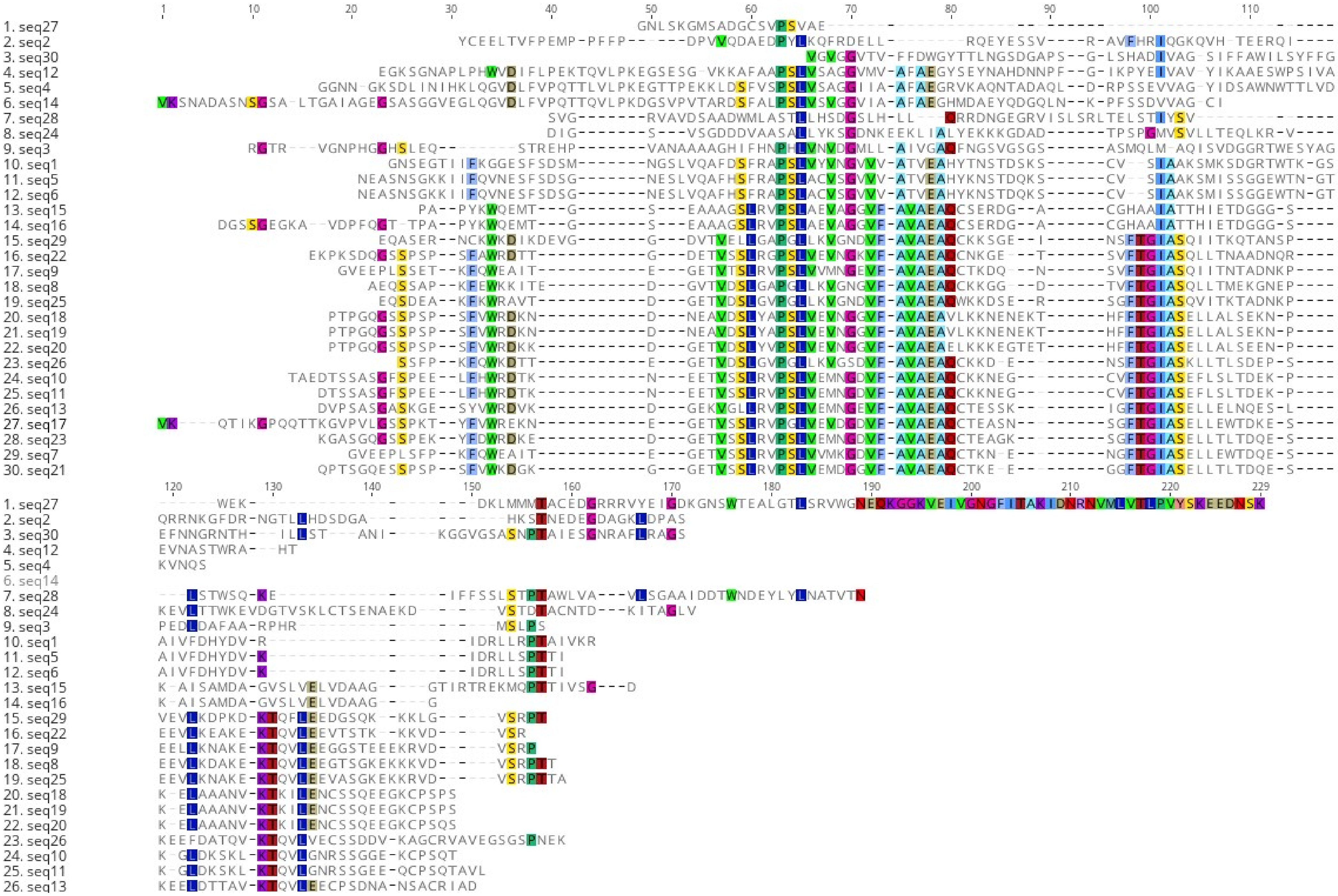

### Fig. S12

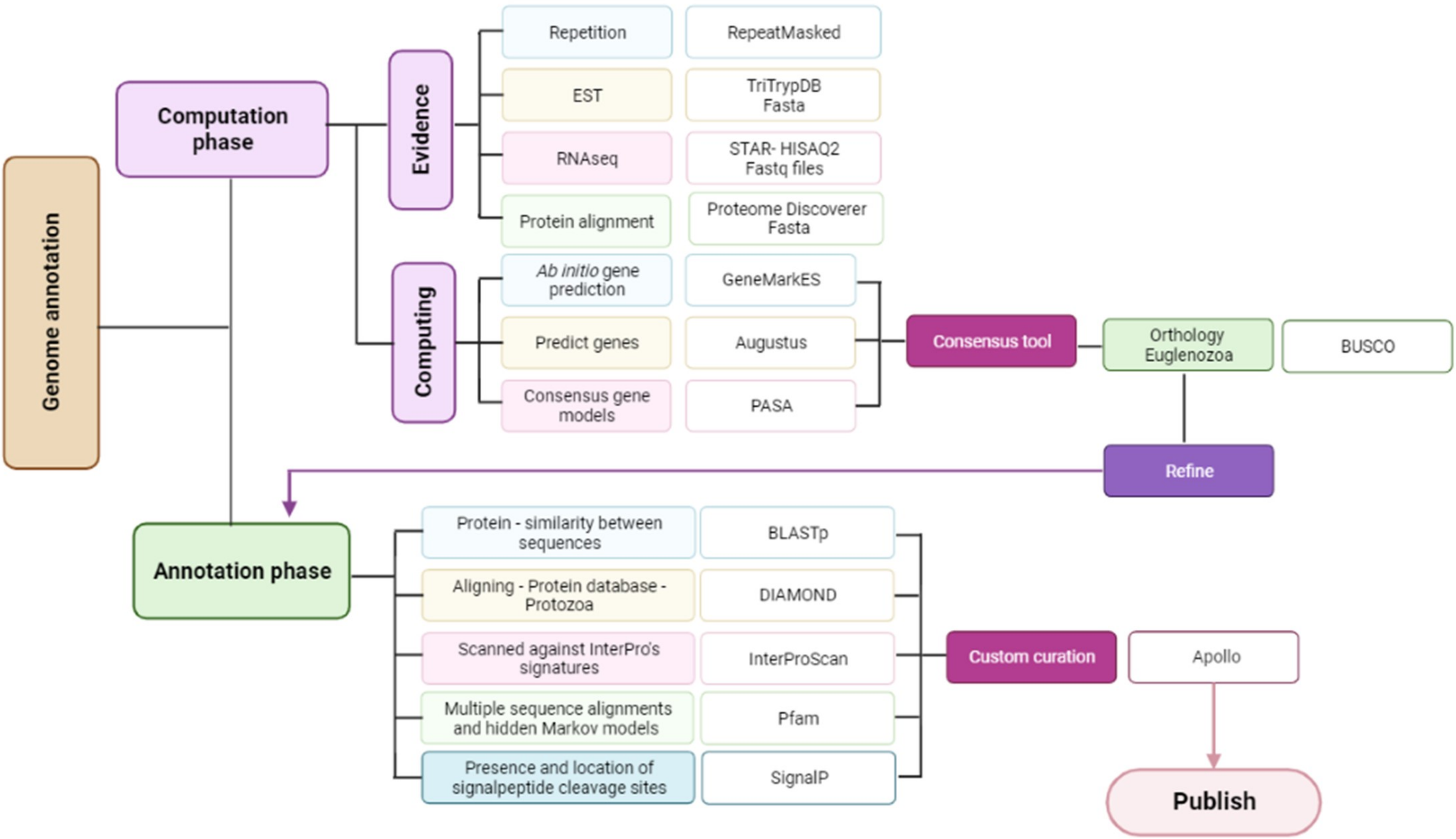
