## Supplementary material for "Stochastic variation in surface protein expression diversifies *Trypanosoma cruzi* infection": Fig. S2

### Chr_16: 8 scaffolds synteny

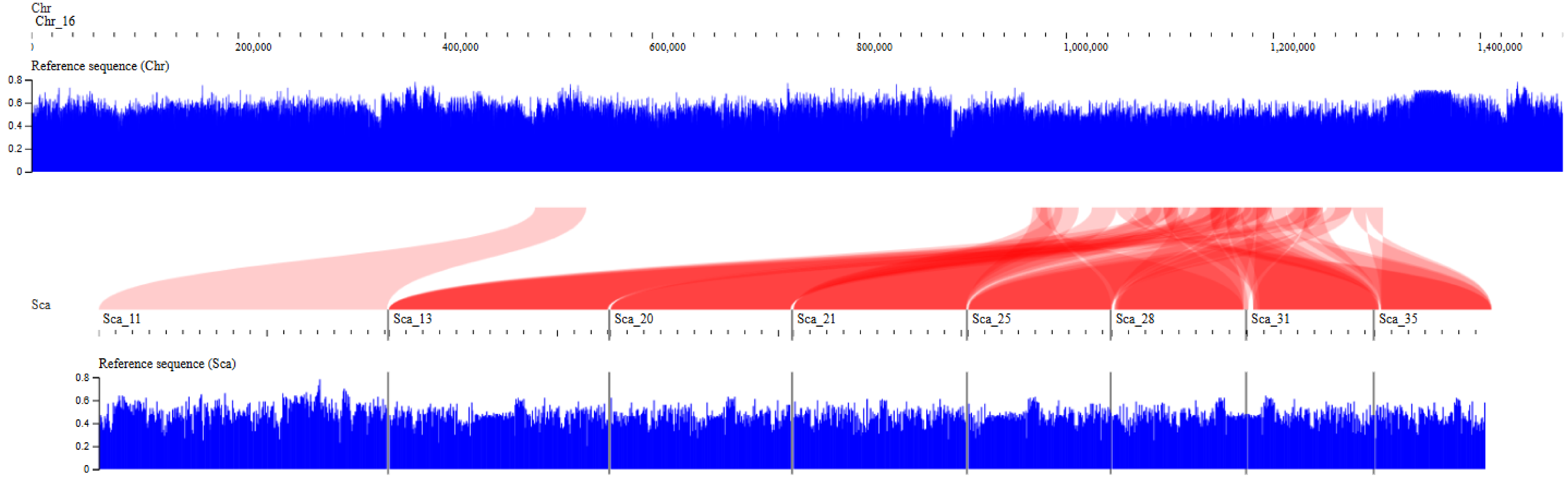

### Chr_30: 6 scaffolds synteny

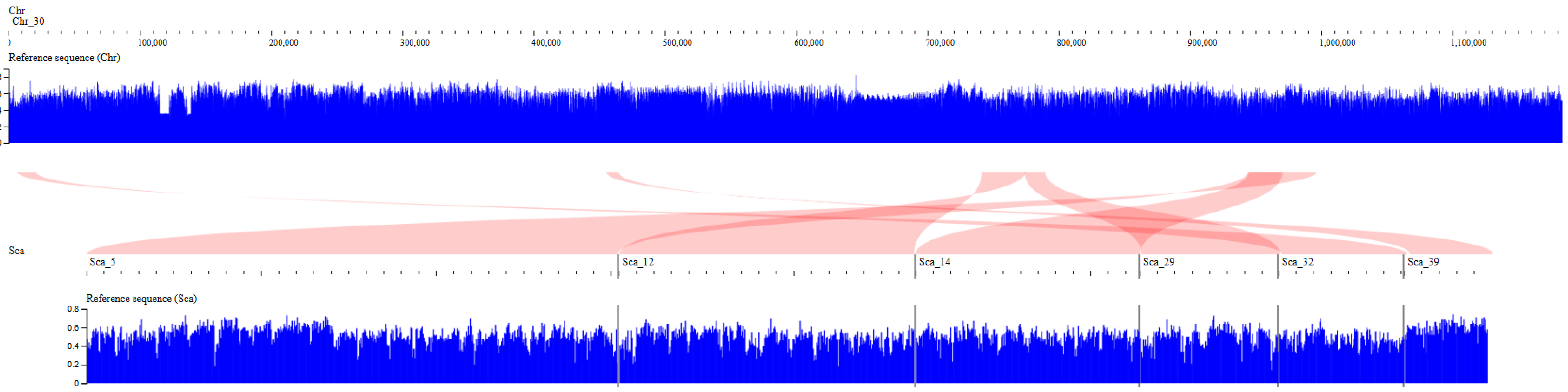

### Chr_26: 4 scaffolds synteny

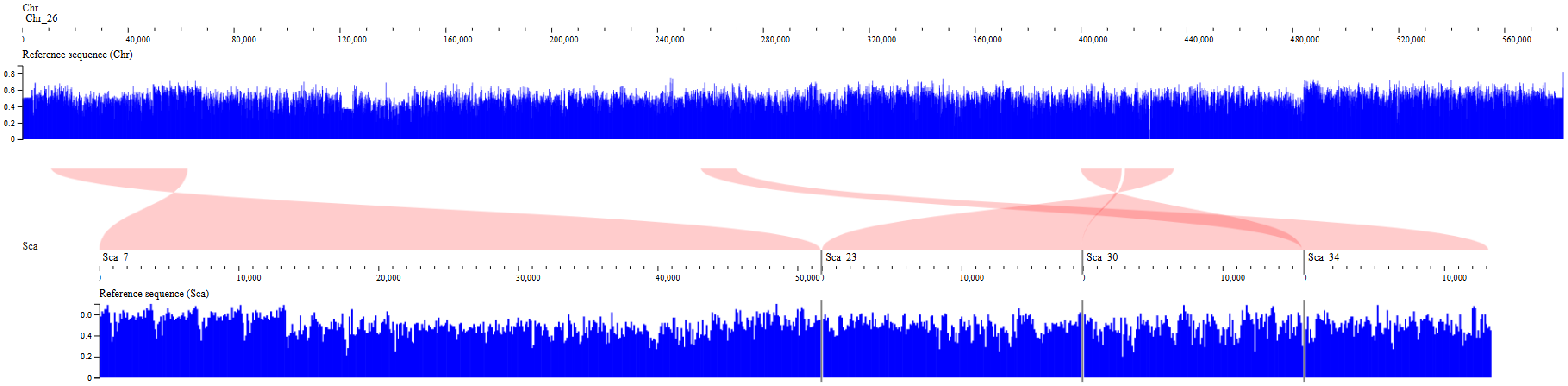

### Chr_7: 3 scaffolds synteny

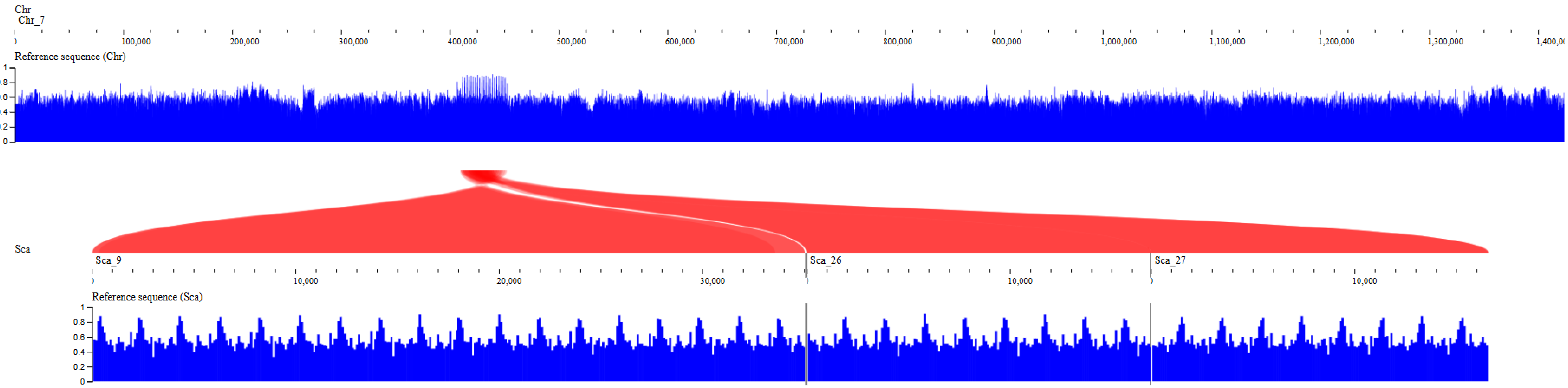

### Chr22: 2 scaffolds synteny

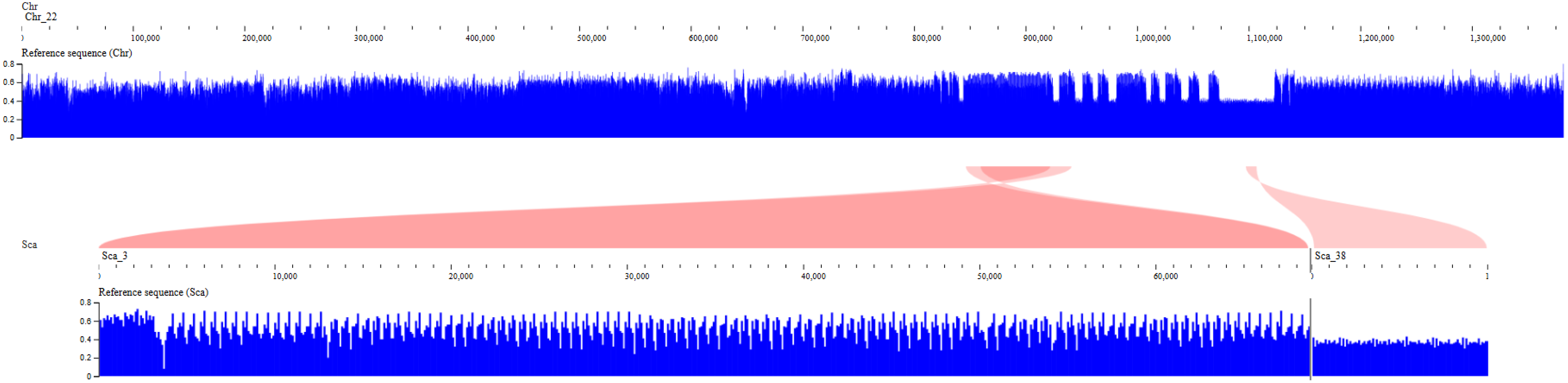

### Chr2: 2 scaffolds synteny

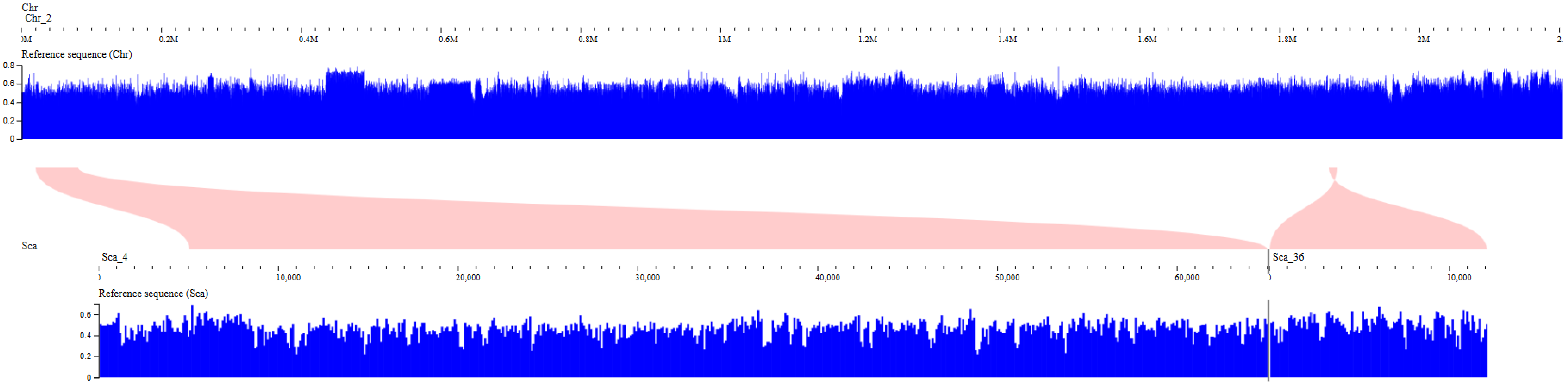

### Chr4: 2 scaffolds synteny

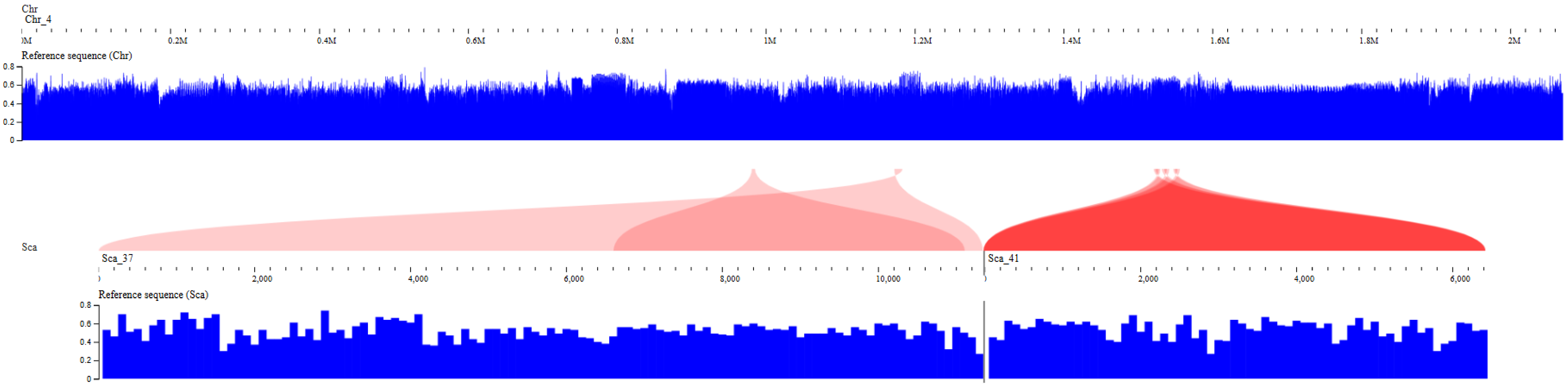

### Chr8: 2 scaffolds synteny

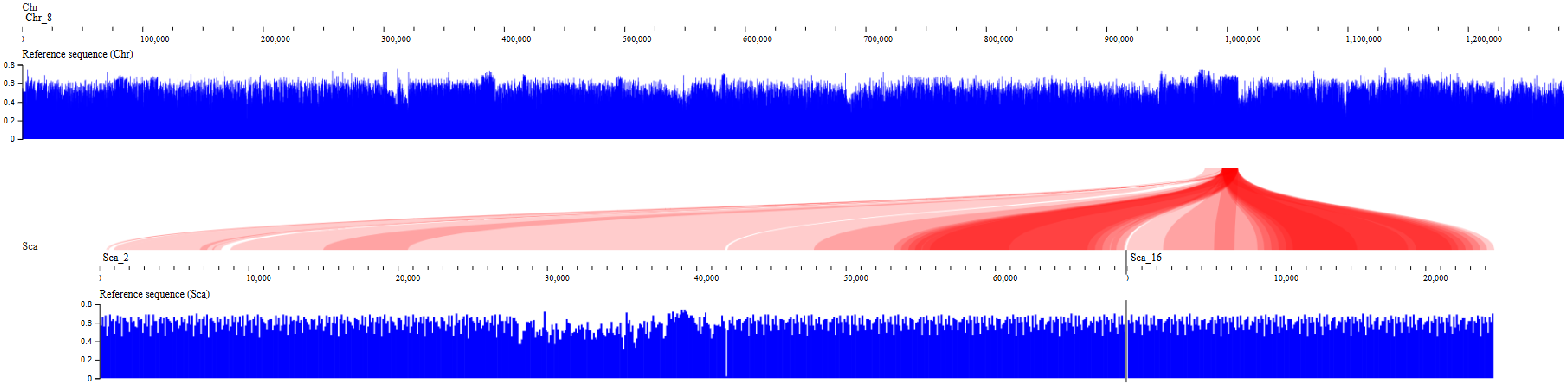

### Chr10: 2 scaffolds synteny

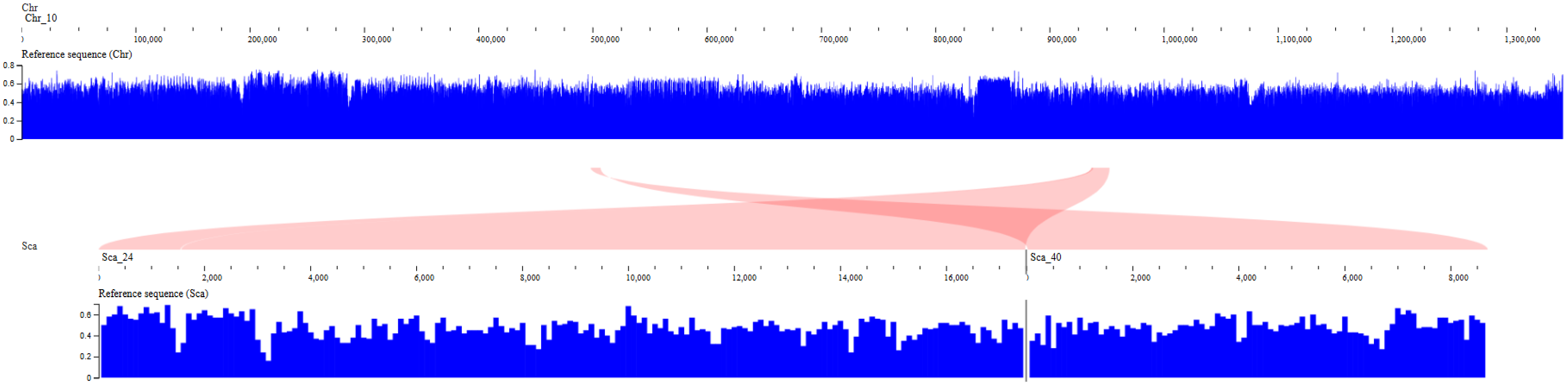

### Chr12: 2 scaffolds synteny

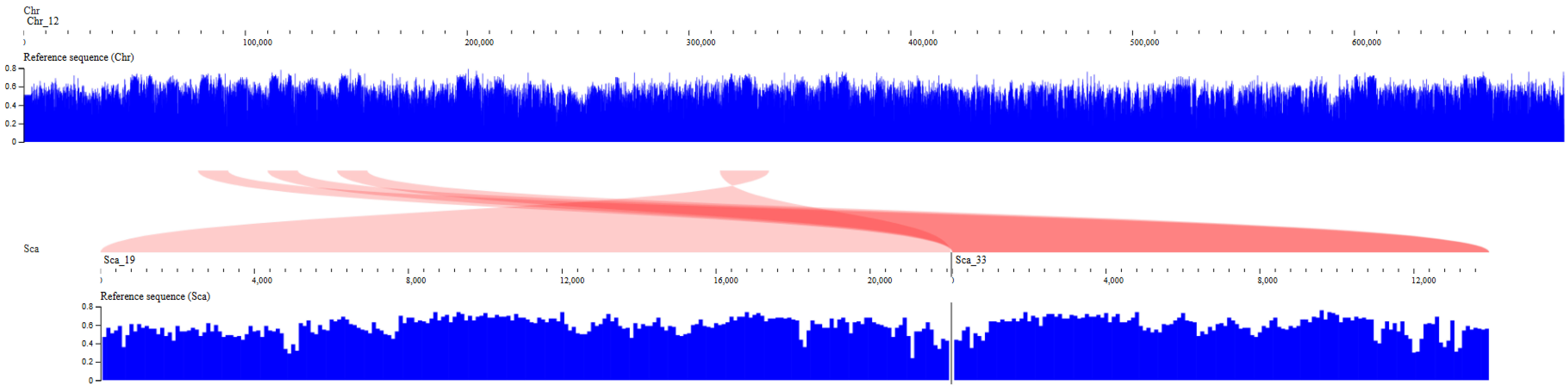

### Chr15: 1 scaffold synteny

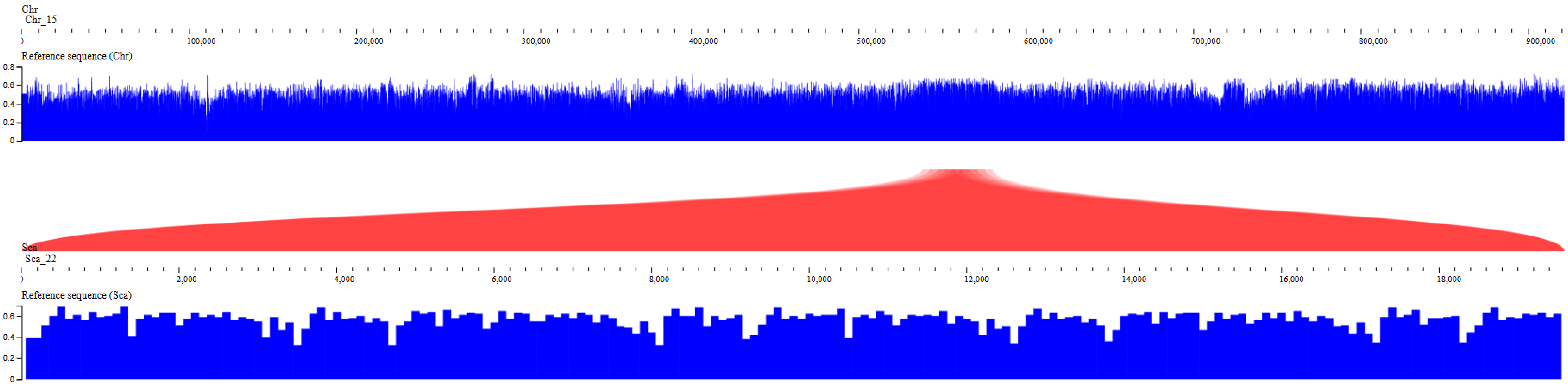

### Chr25: 1 scaffold synteny

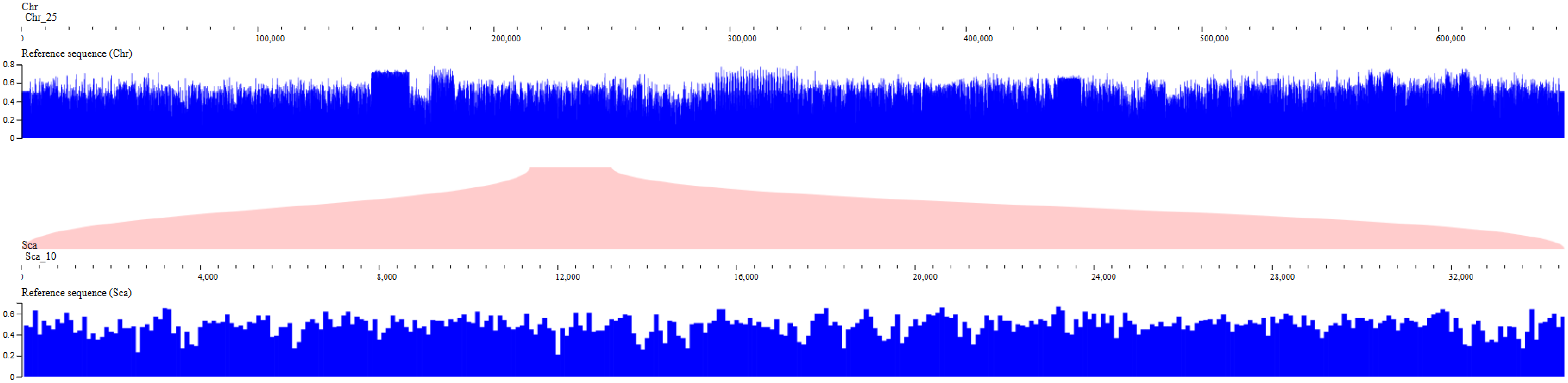

### Chr9: 1 scaffolds synteny

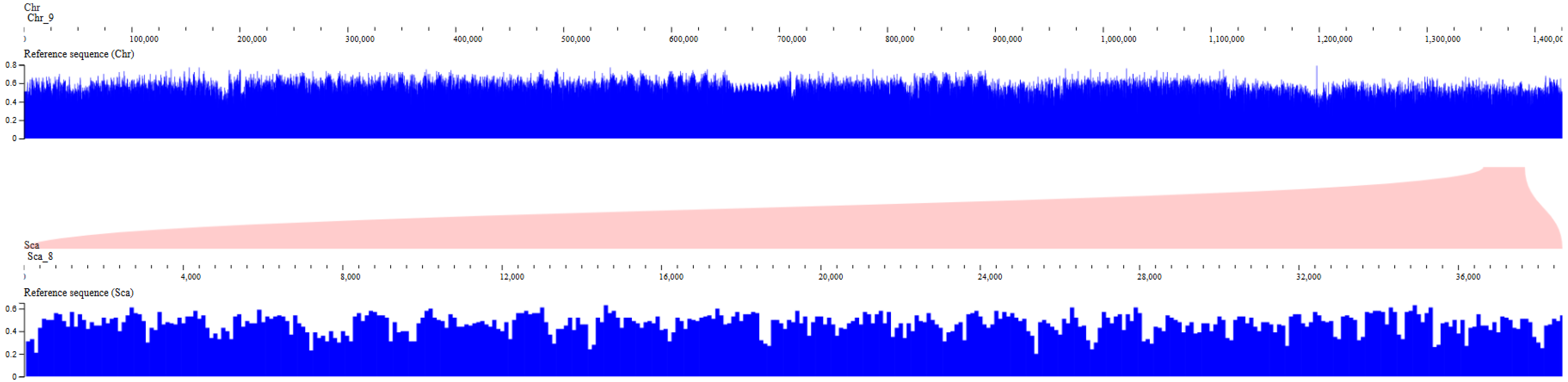

### Chr3-11: 1 scaffolds synteny

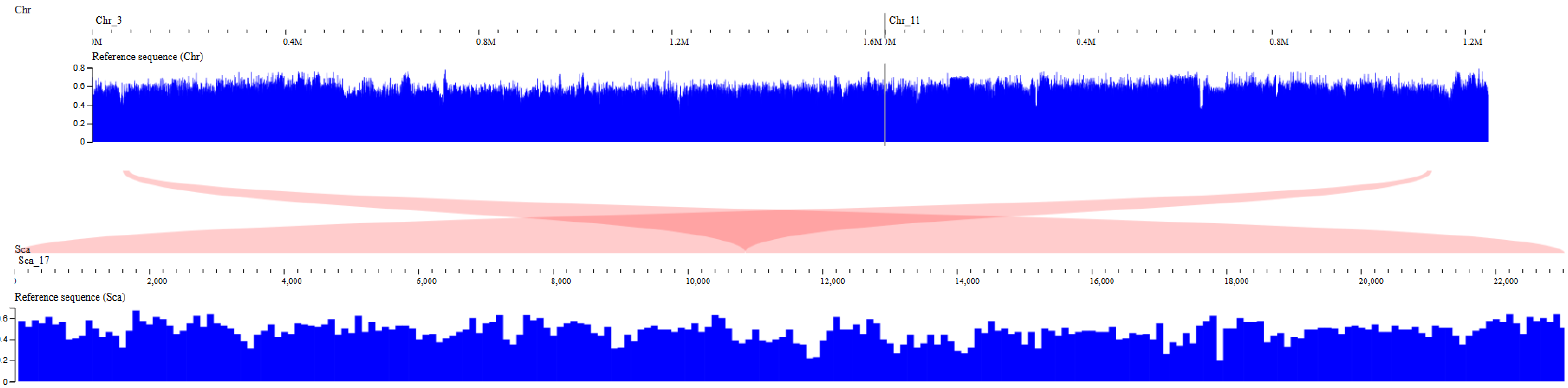

### Chr7-28: 1 scaffolds synteny

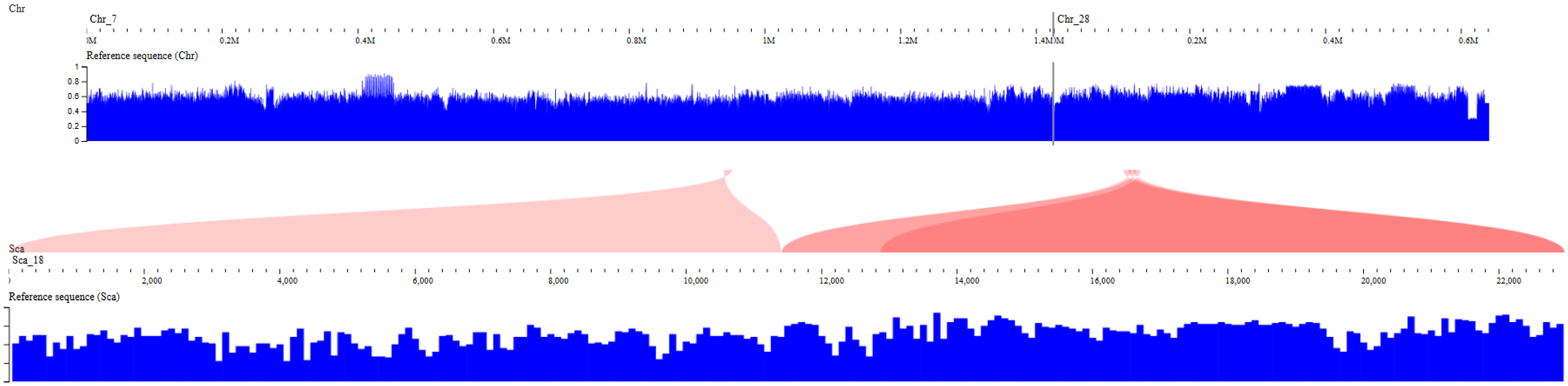
