## Supplementary material for "Stochastic variation in surface protein expression diversifies *Trypanosoma cruzi* infection": Fig. S4

***T. cruzi* Sylvio X10**

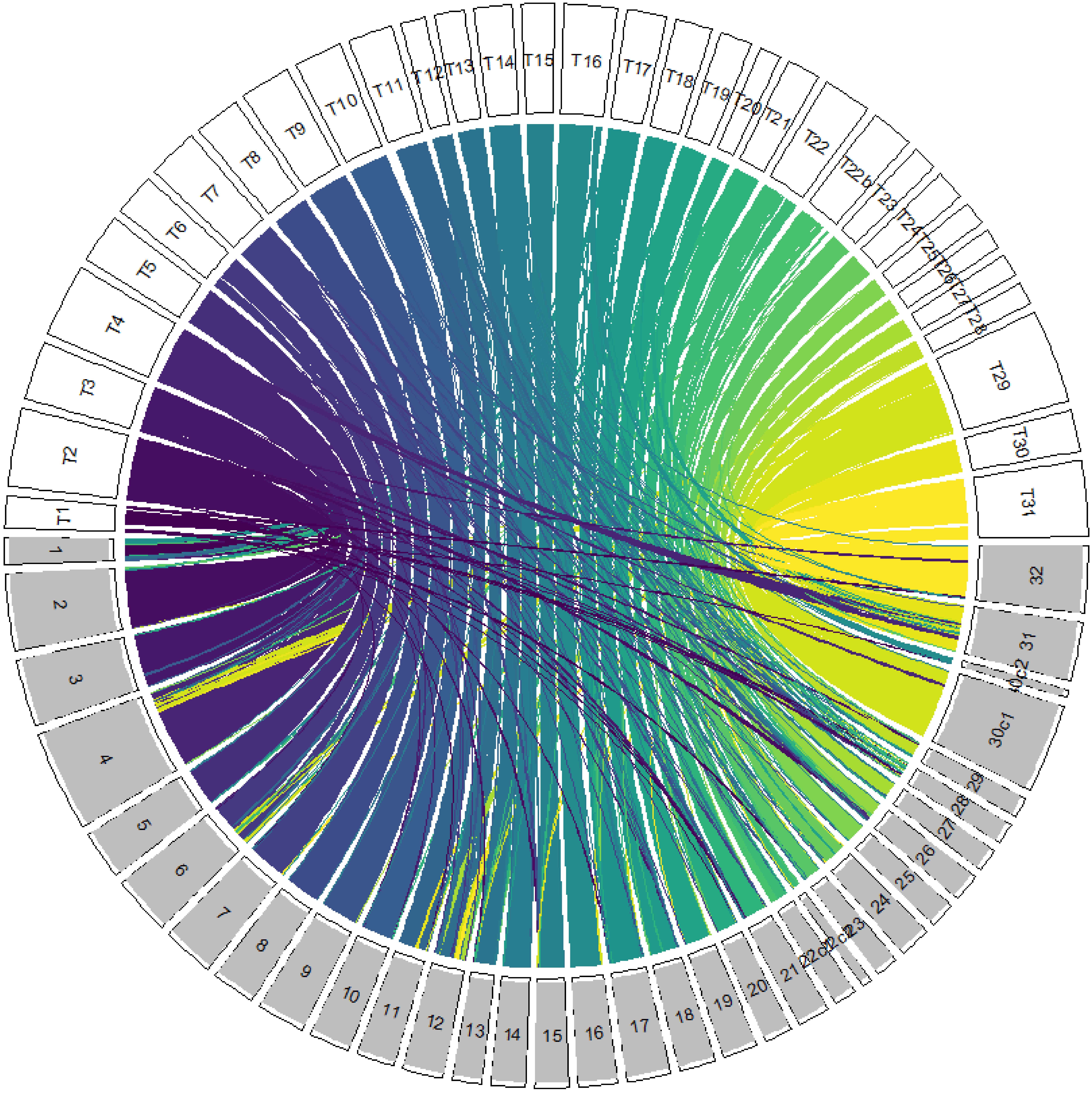


***T. cruzi* DM25**

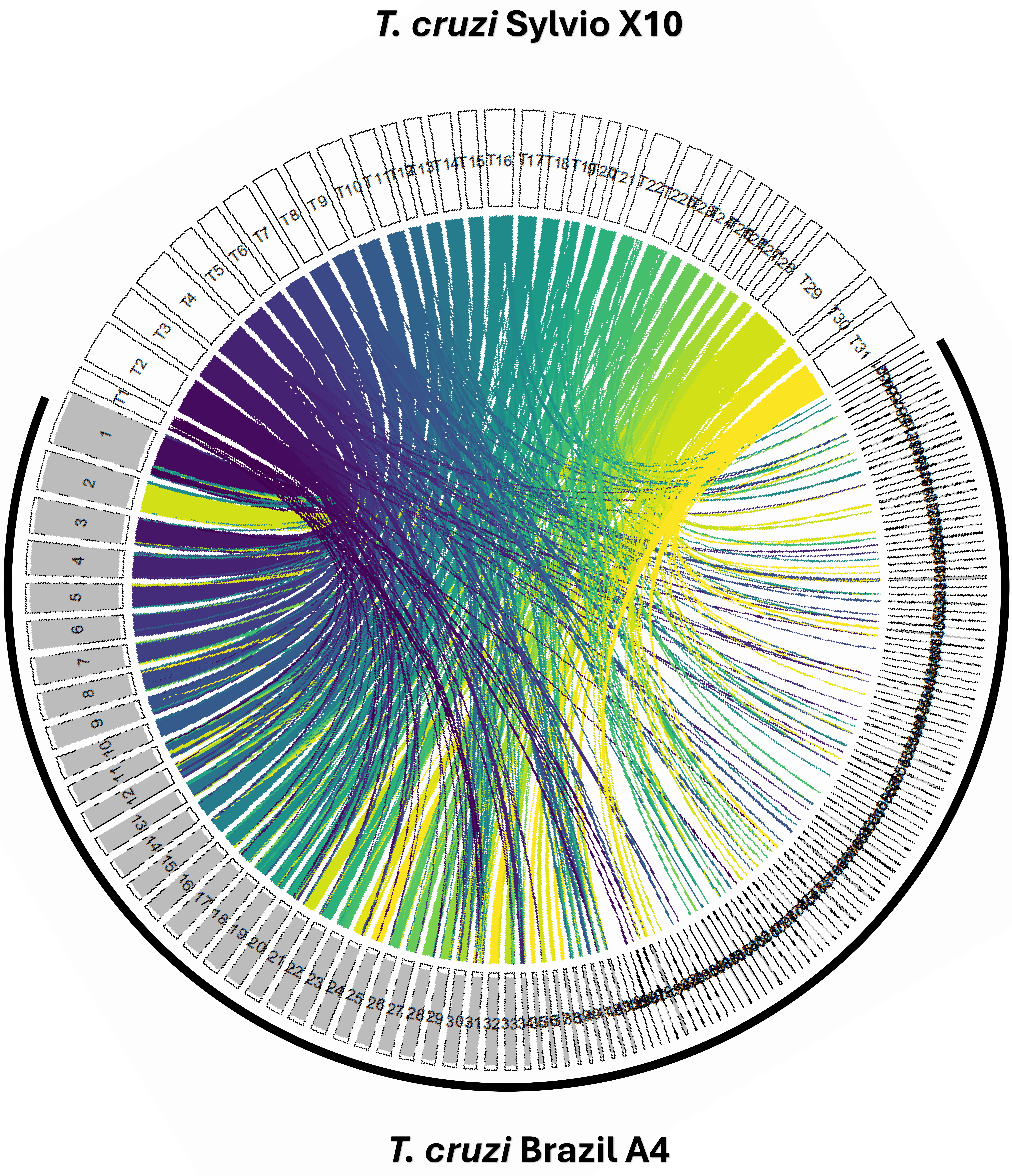

*T. cruzi* Sylvio X10
*T. cruzi* Brazil A4

***T. cruzi* Sylvio X10**

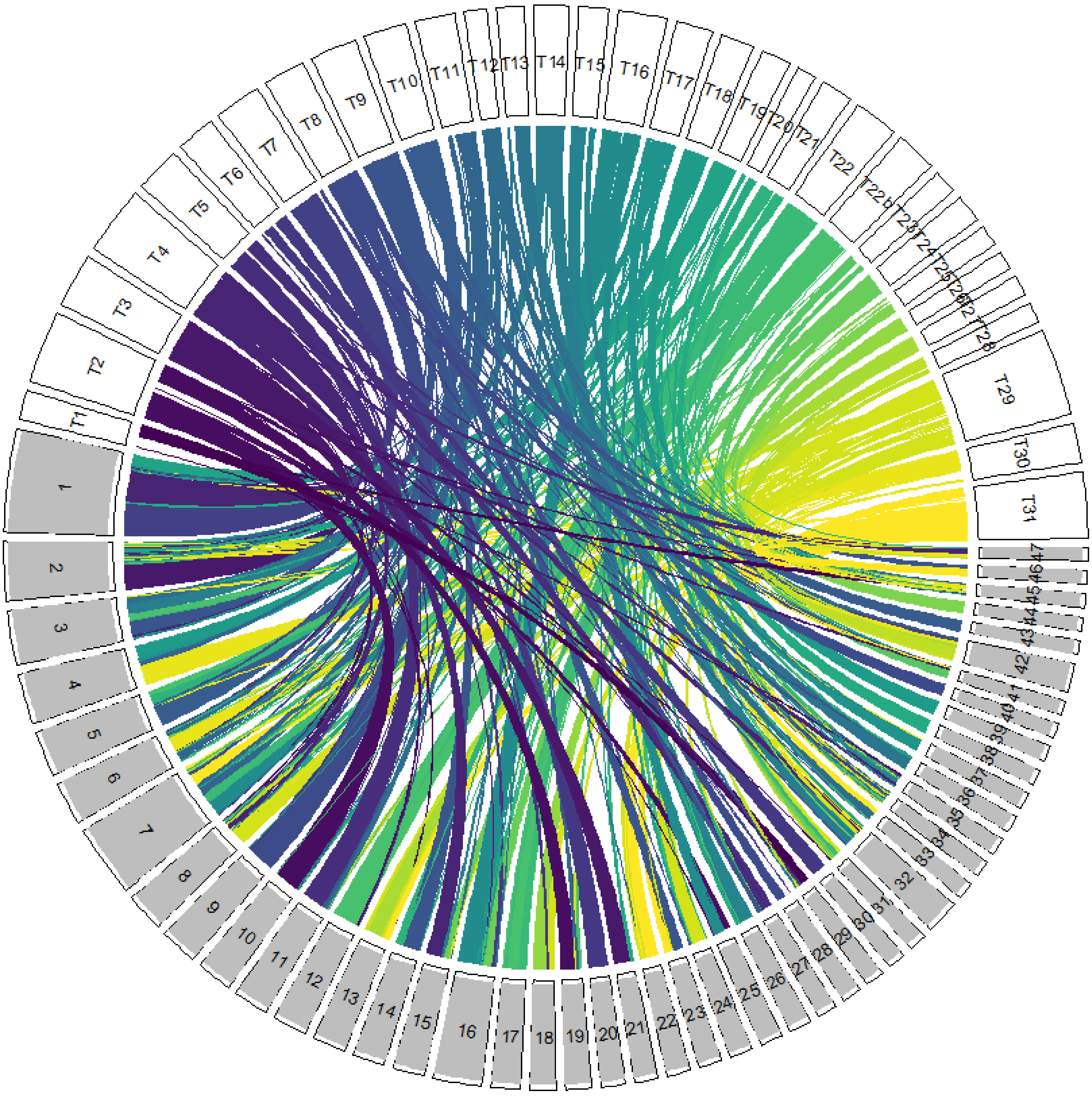


***T. cruzi* Sylvio X10 - 2018**

### *T. cruzi* Sylvio X10

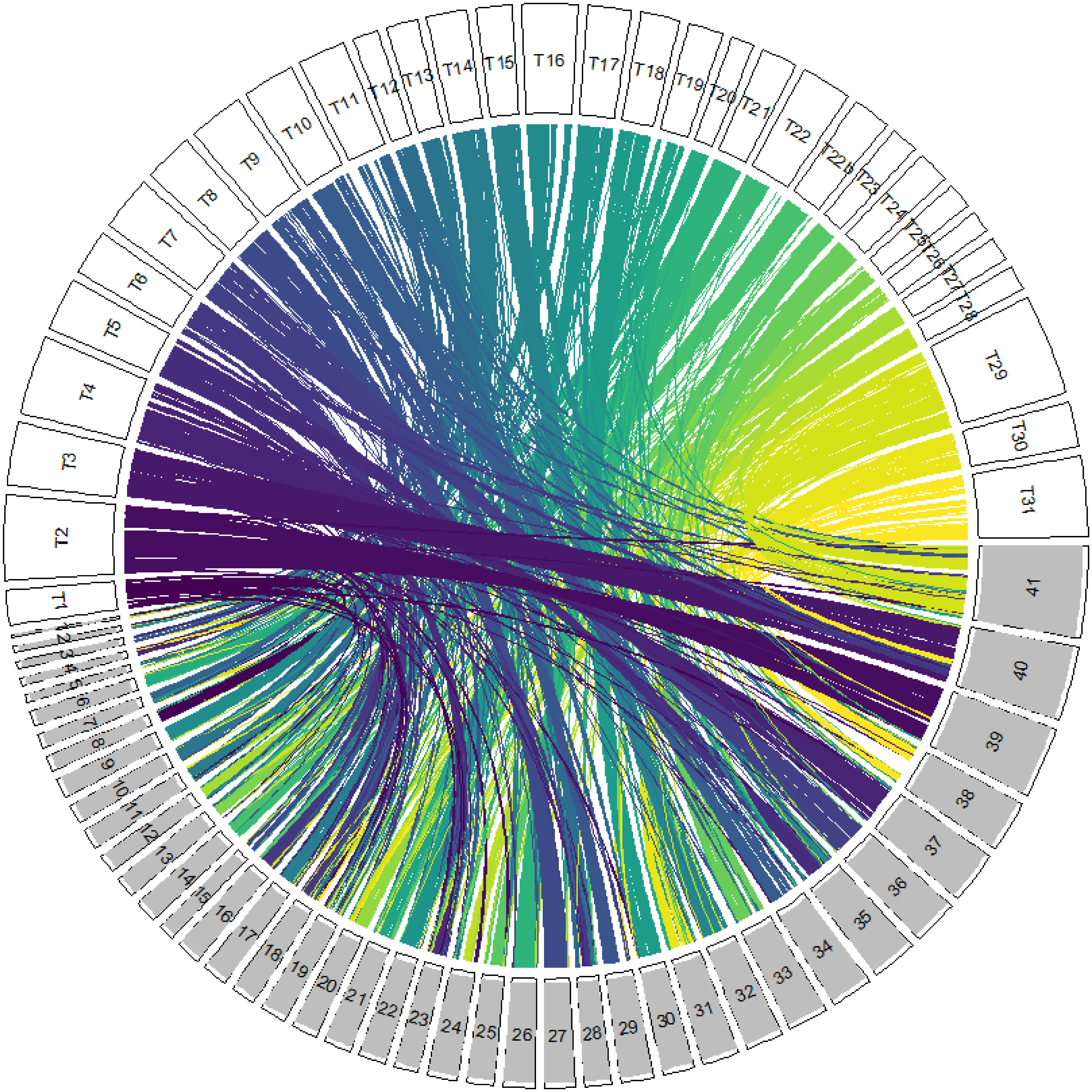


### *T. cruzi* Sylvio Cl-Brenner Esmeraldo like
