## Supplementary material for "Stochastic variation in surface protein expression diversifies *Trypanosoma cruzi* infection": Fig. S7

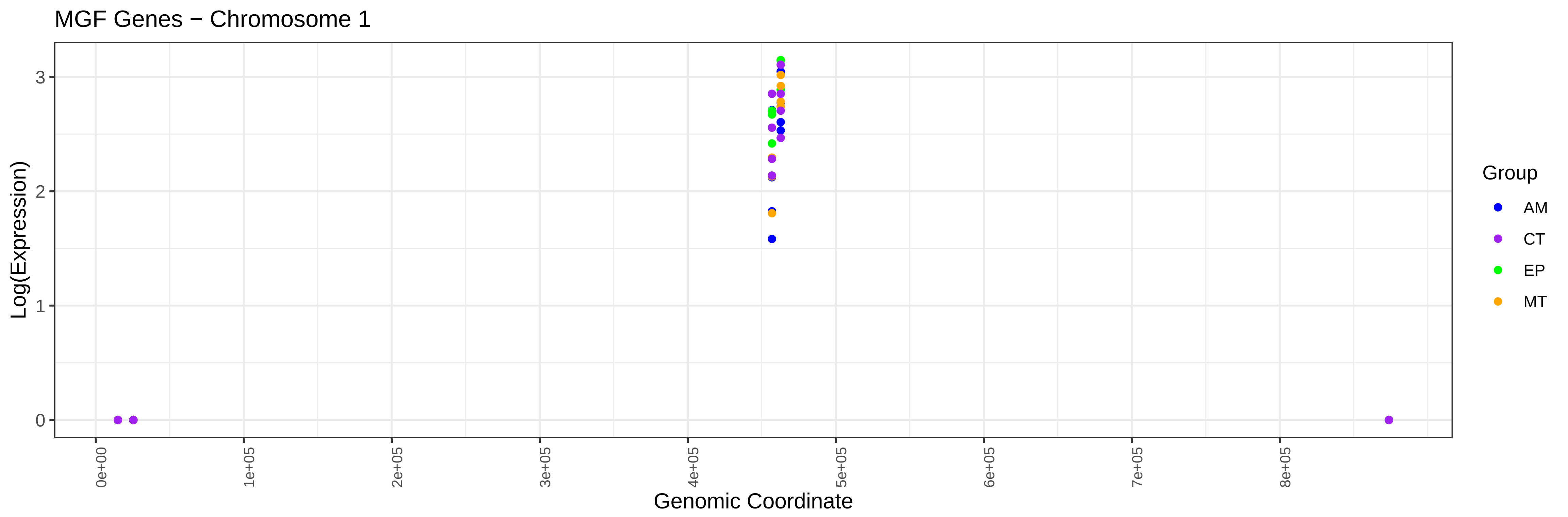

MGF Genes − Chromosome 1
Log(Expression)
0
1
2
3
0e+00
1e+05
2e+05
3e+05
4e+05
5e+05
6e+05
7e+05
8e+05
Genomic Coordinate
Group
AM
CT
EP
MT


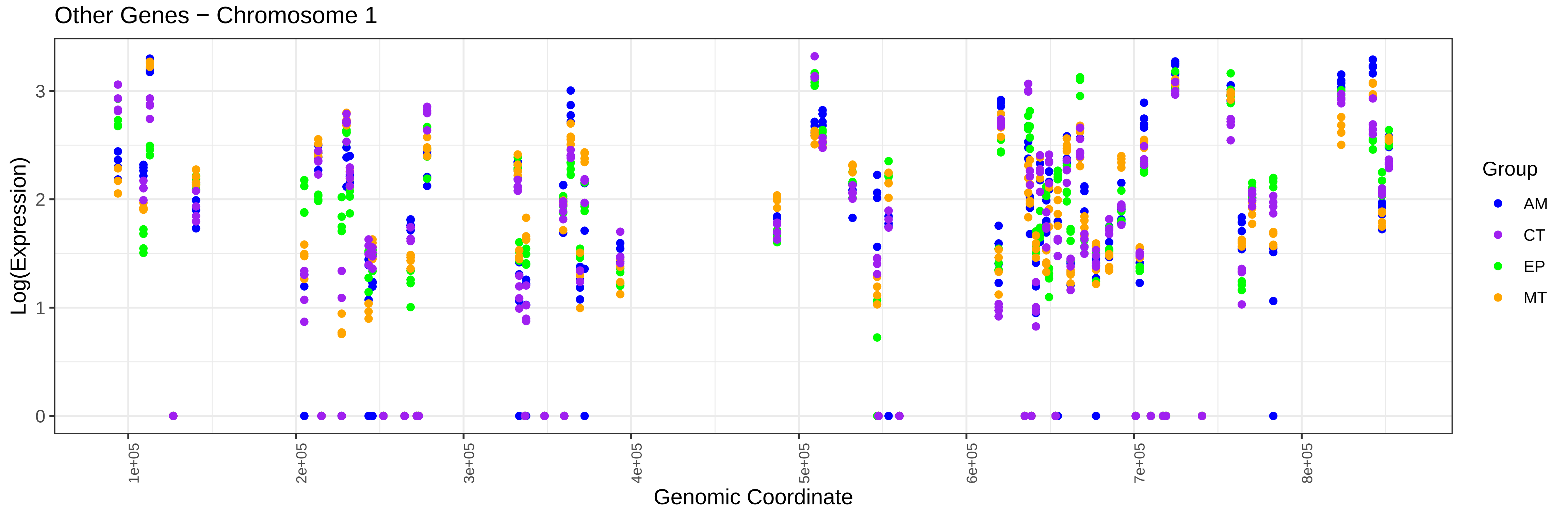

Other Genes − Chromosome 1
Log(Expression)
0
1
2
3
1e+05
2e+05
3e+05
4e+05
5e+05
6e+05
7e+05
8e+05
Genomic Coordinate
Group
AM
CT
EP
MT

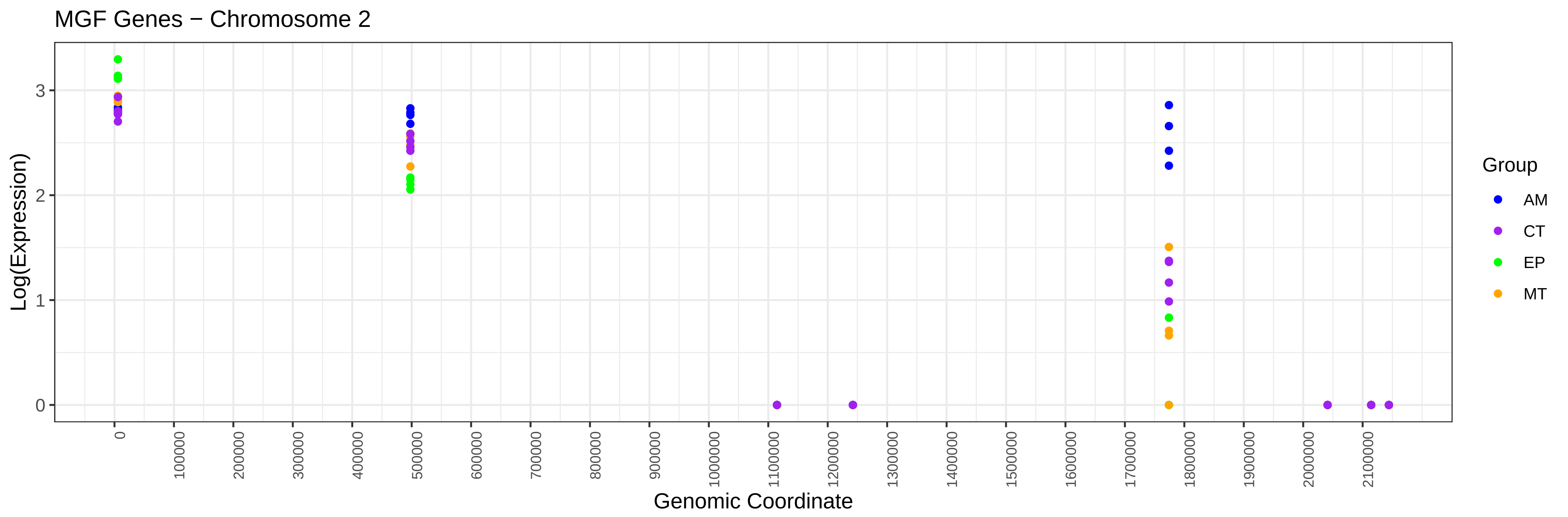
MGF Genes − Chromosome 2
Log(Expression)
Genomic Coordinate
Group
AM
CT
EP
MT

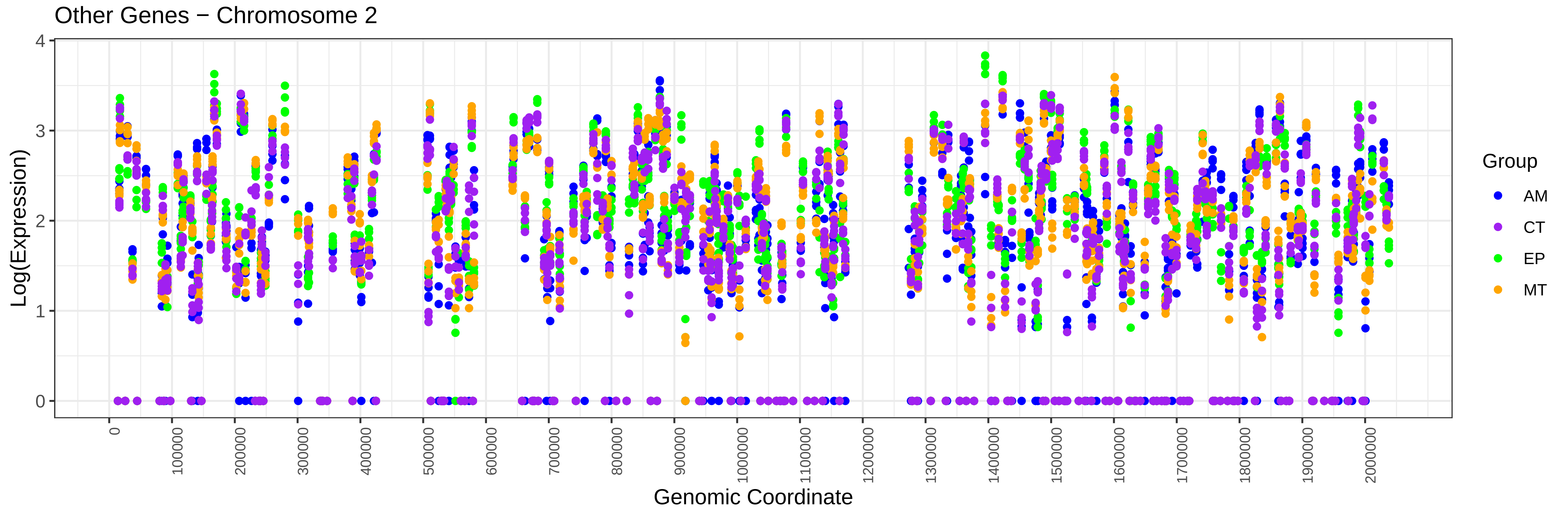
Other Genes − Chromosome 2
Log(Expression)
Genomic Coordinate
Group
AM
CT
EP
MT

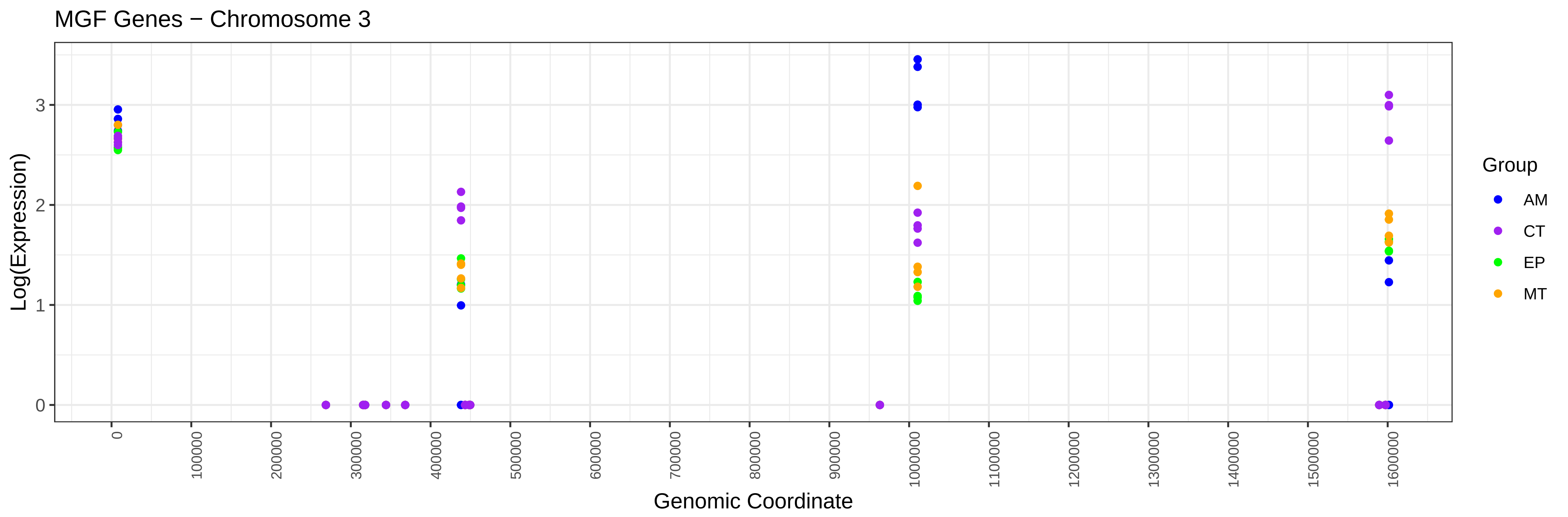
MGF Genes − Chromosome 3
Log(Expression)
0
1
2
3
0
100000
200000
300000
400000
500000
600000
700000
800000
900000
1000000
1100000
1200000
1300000
1400000
1500000
1600000
Genomic Coordinate
Group
AM
CT
EP
MT

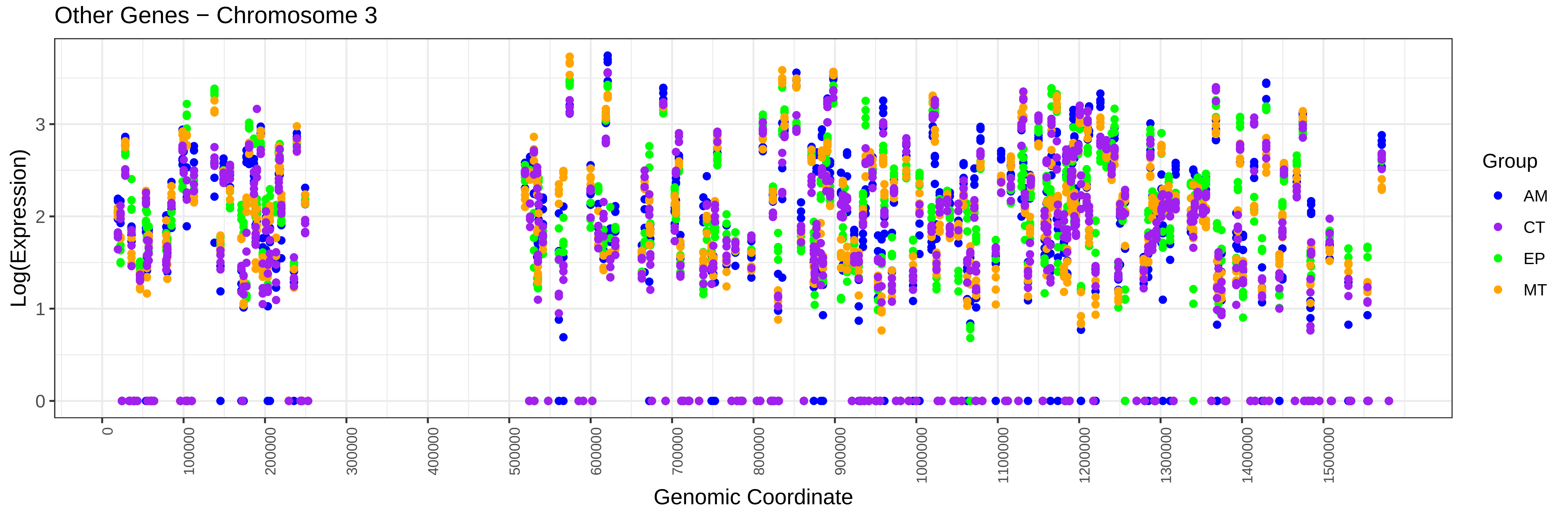
Other Genes − Chromosome 3
Log(Expression)
0
1
2
3
0
100000
200000
300000
400000
500000
600000
700000
800000
900000
1000000
1100000
1200000
1300000
1400000
1500000
Genomic Coordinate
Group
AM
CT
EP
MT

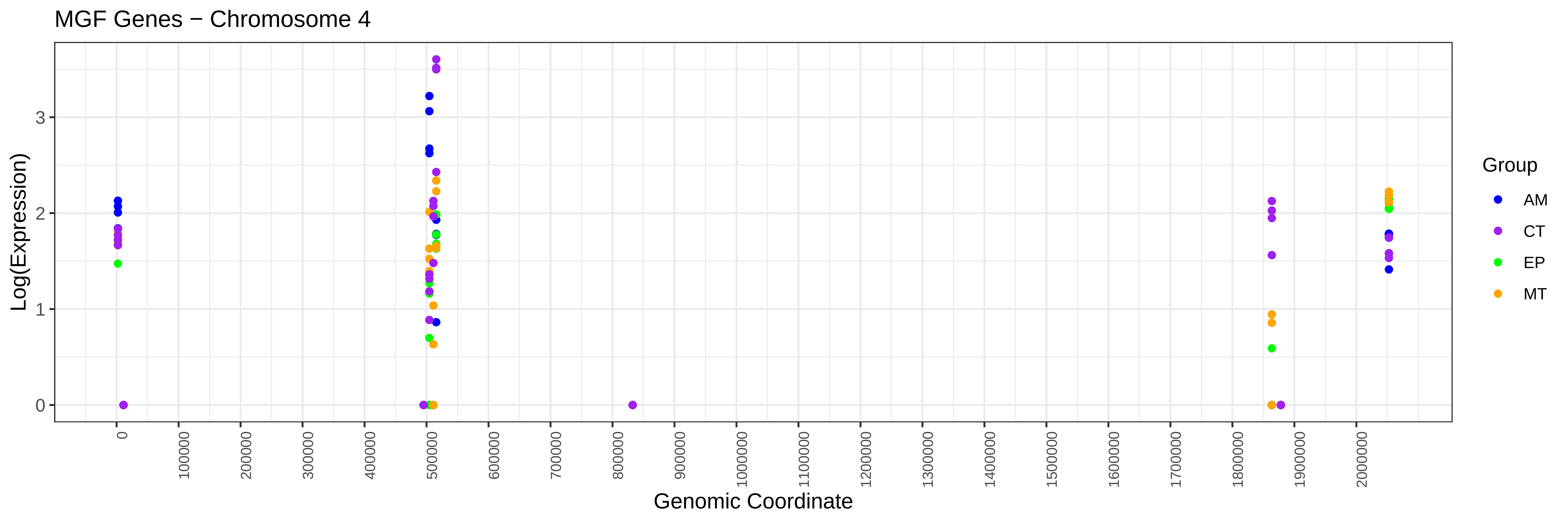
MGF Genes − Chromosome 4
Log(Expression)
0
1
2
3
0
100000
200000
300000
400000
500000
600000
700000
800000
900000
1000000
1100000
1200000
1300000
1400000
1500000
1600000
1700000
1800000
1900000
2000000
Genomic Coordinate
Group
AM
CT
EP
MT

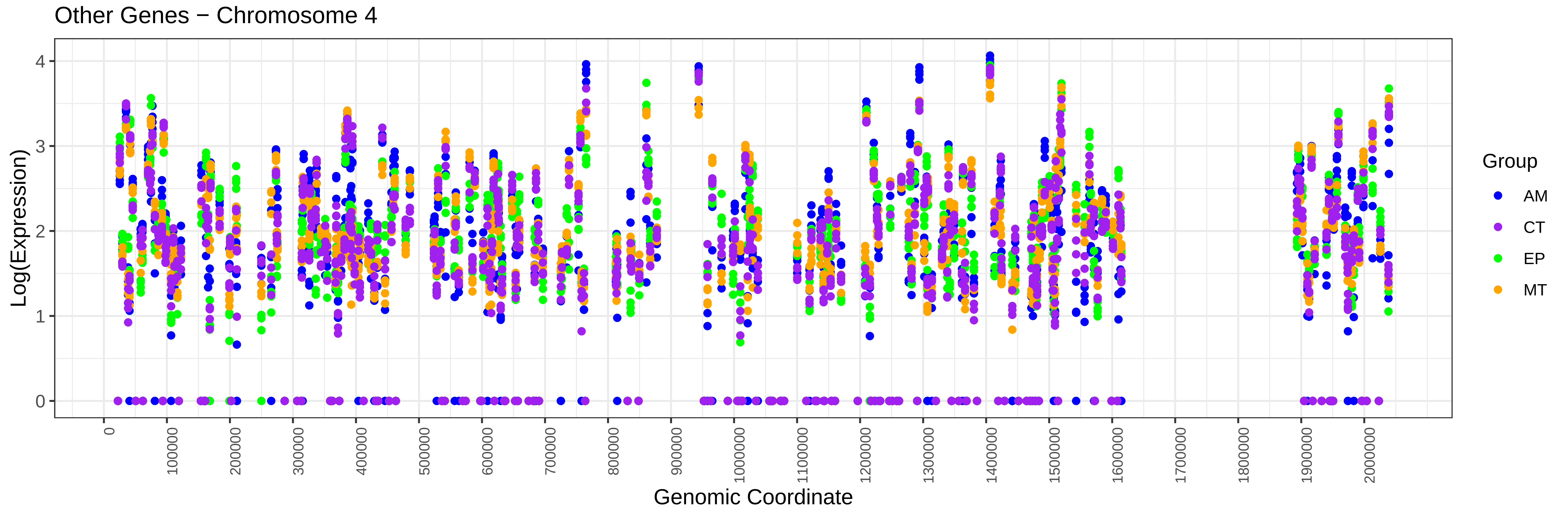
Other Genes − Chromosome 4
Log(Expression)
0
1
2
3
4
0
100000
200000
300000
400000
500000
600000
700000
800000
900000
1000000
1100000
1200000
1300000
1400000
1500000
1600000
1700000
1800000
1900000
2000000
Genomic Coordinate
Group
AM
CT
EP
MT

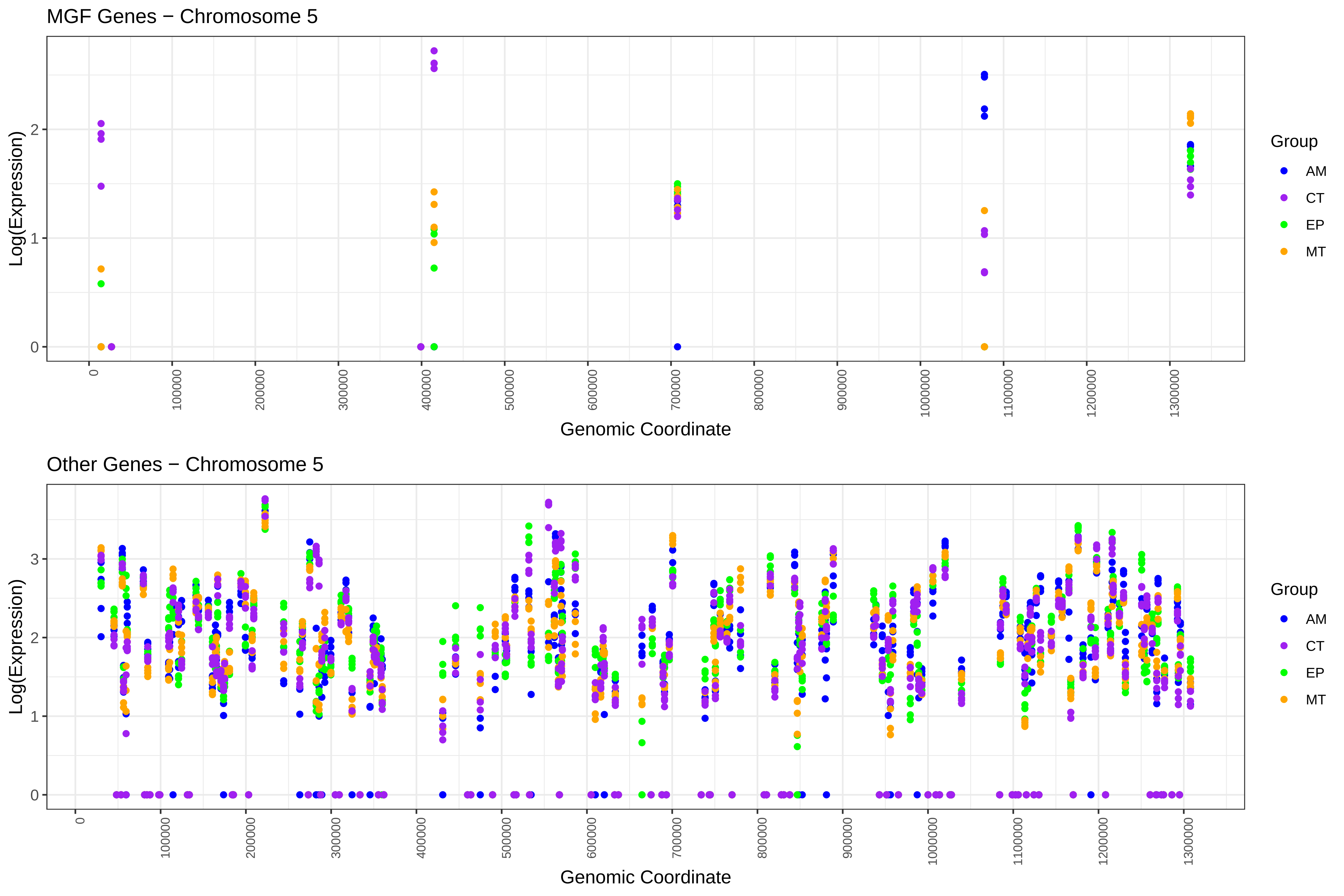
MGF Genes − Chromosome 5
Log(Expression)
0
1
2
0
100000
200000
300000
400000
500000
600000
700000
800000
900000
1000000
1100000
1200000
1300000
Genomic Coordinate
Group
AM
CT
EP
MT
Other Genes − Chromosome 5
Log(Expression)
0
1
2
3
0
100000
200000
300000
400000
500000
600000
700000
800000
900000
1000000
1100000
1200000
1300000
Genomic Coordinate
Group
AM
CT
EP
MT

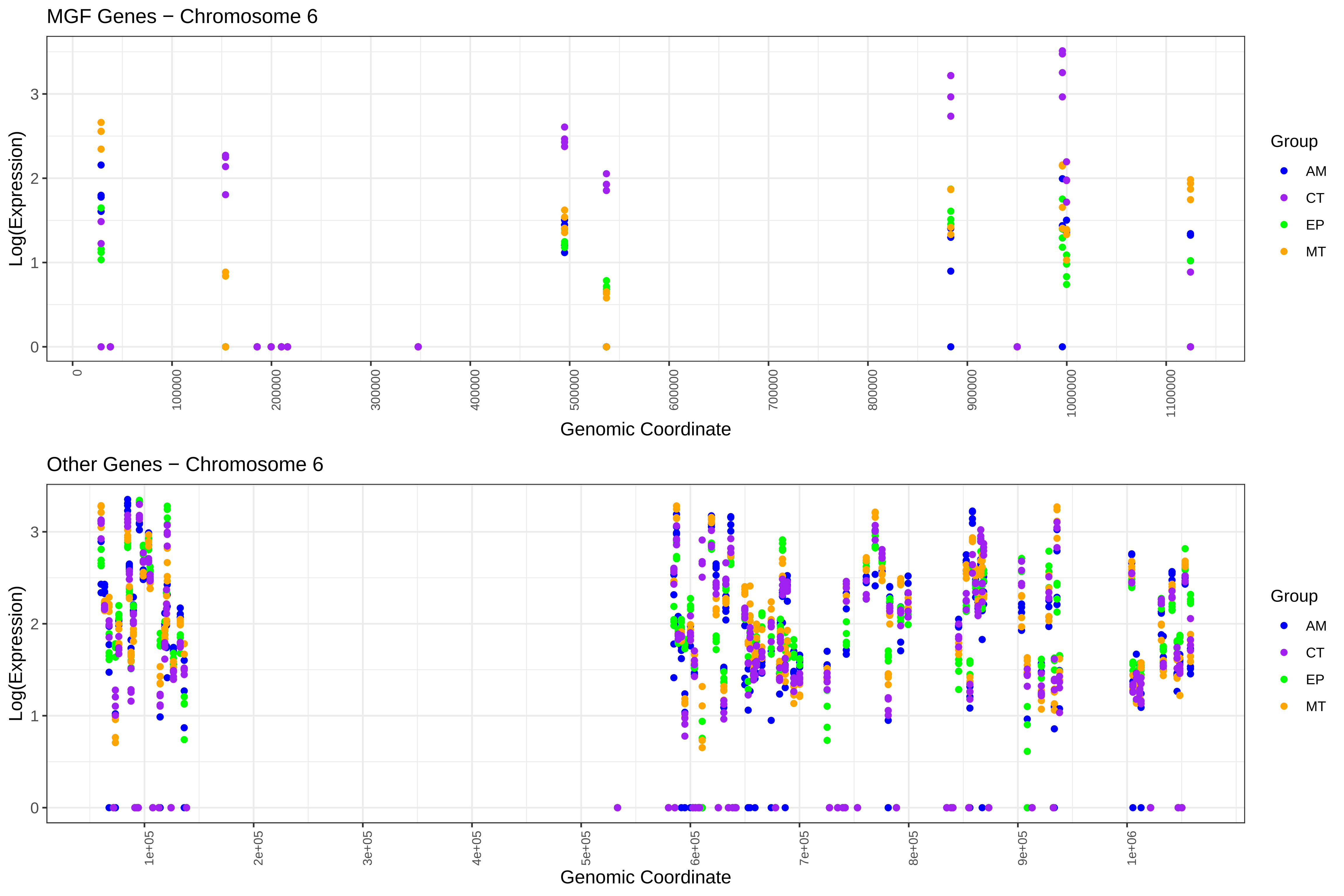

MGF Genes − Chromosome 6
Log(Expression)
0
1
2
3
0
100000
200000
300000
400000
500000
600000
700000
800000
900000
1000000
1100000
Genomic Coordinate
Group
AM
CT
EP
MT
Other Genes − Chromosome 6
Log(Expression)
0
1
2
3
1e+05
2e+05
3e+05
4e+05
5e+05
6e+05
7e+05
8e+05
9e+05
1e+06
Genomic Coordinate
Group
AM
CT
EP
MT

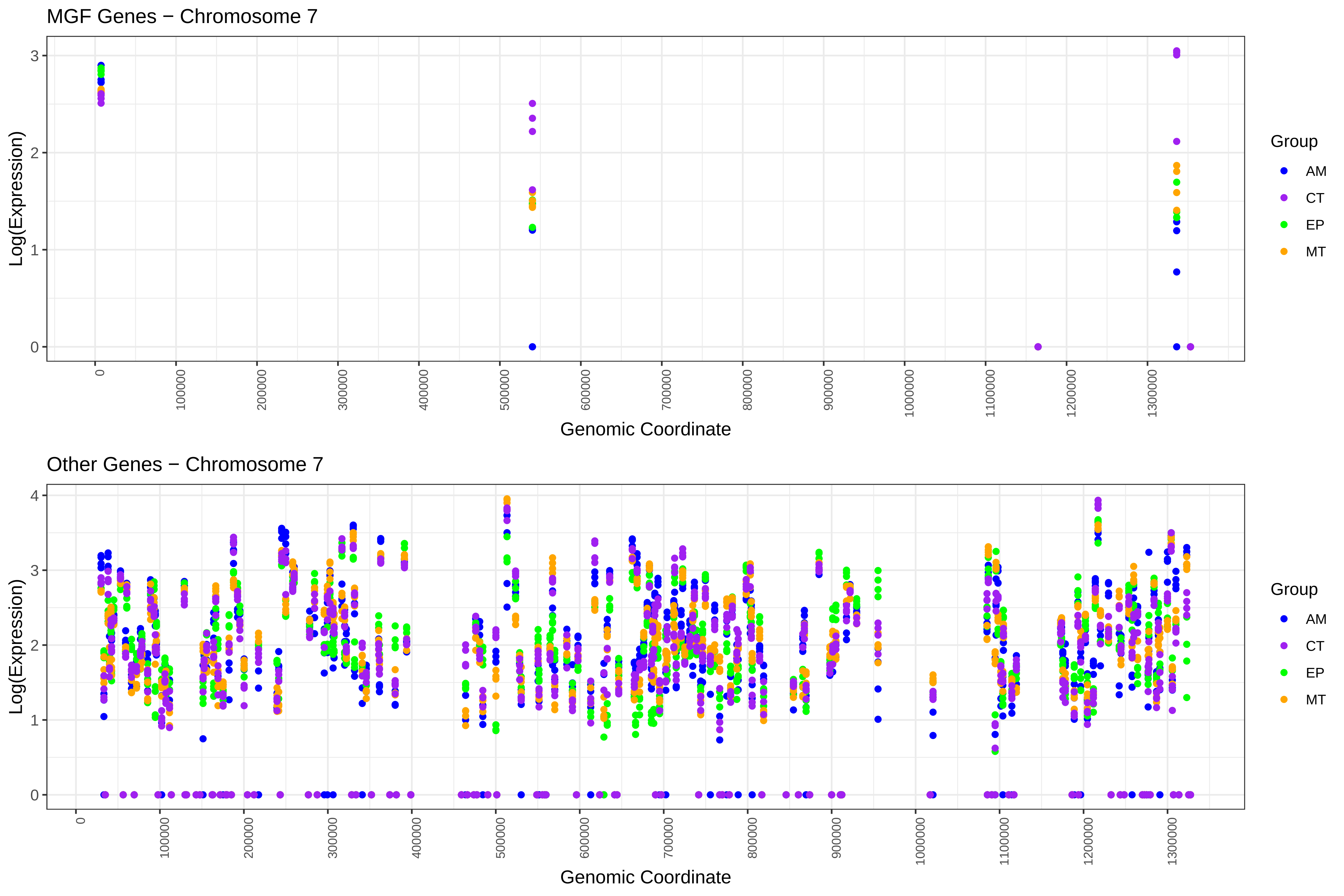

MGF Genes − Chromosome 7
Log(Expression)
Genomic Coordinate
Group
AM
CT
EP
MT
0
1
2
3
0
100000
200000
300000
400000
500000
600000
700000
800000
900000
1000000
1100000
1200000
1300000
Other Genes − Chromosome 7
Log(Expression)
Genomic Coordinate
Group
AM
CT
EP
MT
0
1
2
3
4

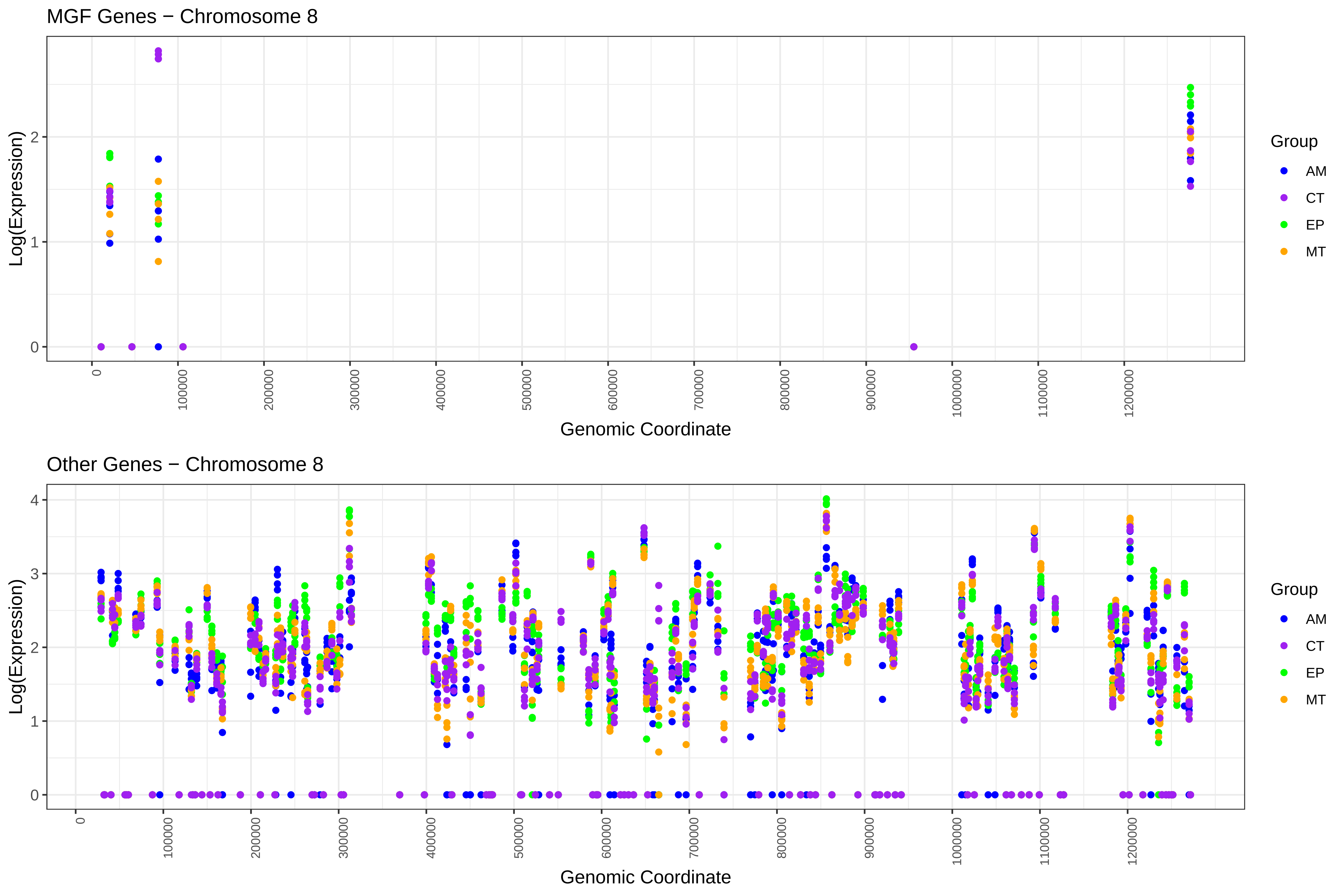

MGF Genes − Chromosome 8
Log(Expression)
0
1
2
0
100000
200000
300000
400000
500000
600000
700000
800000
900000
1000000
1100000
1200000
Genomic Coordinate
Group
AM
CT
EP
MT
Other Genes − Chromosome 8
Log(Expression)
0
1
2
3
4
0
100000
200000
300000
400000
500000
600000
700000
800000
900000
1000000
1100000
1200000
Genomic Coordinate
Group
AM
CT
EP
MT

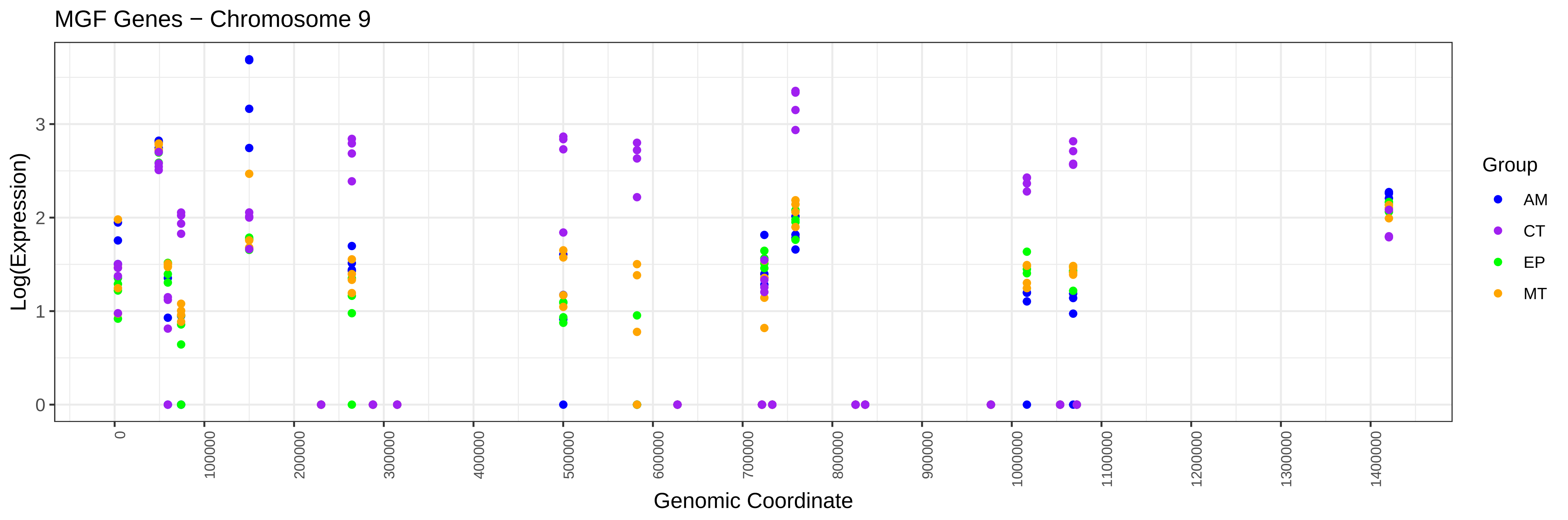
MGF Genes − Chromosome 9
Log(Expression)
0
1
2
3
Genomic Coordinate
0
100000
200000
300000
400000
500000
600000
700000
800000
900000
1000000
1100000
1200000
1300000
1400000
Group
AM
CT
EP
MT

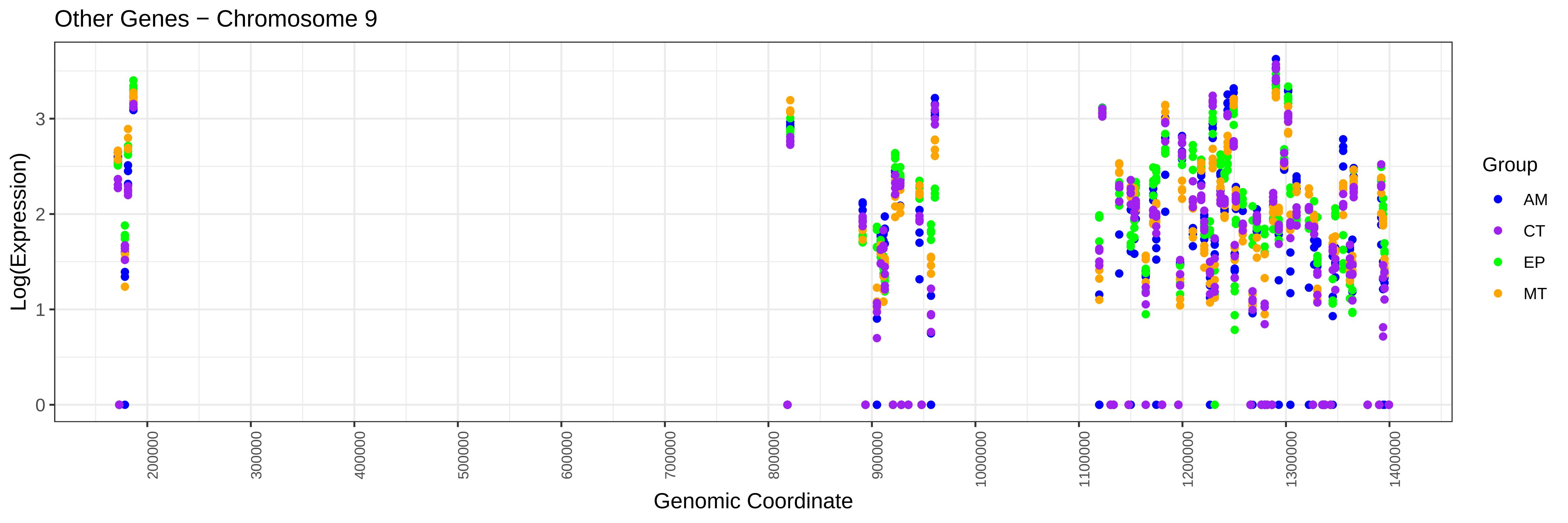
Other Genes − Chromosome 9
Log(Expression)
0
1
2
3
Genomic Coordinate
200000
300000
400000
500000
600000
700000
800000
900000
1000000
1100000
1200000
1300000
1400000
Group
AM
CT
EP
MT

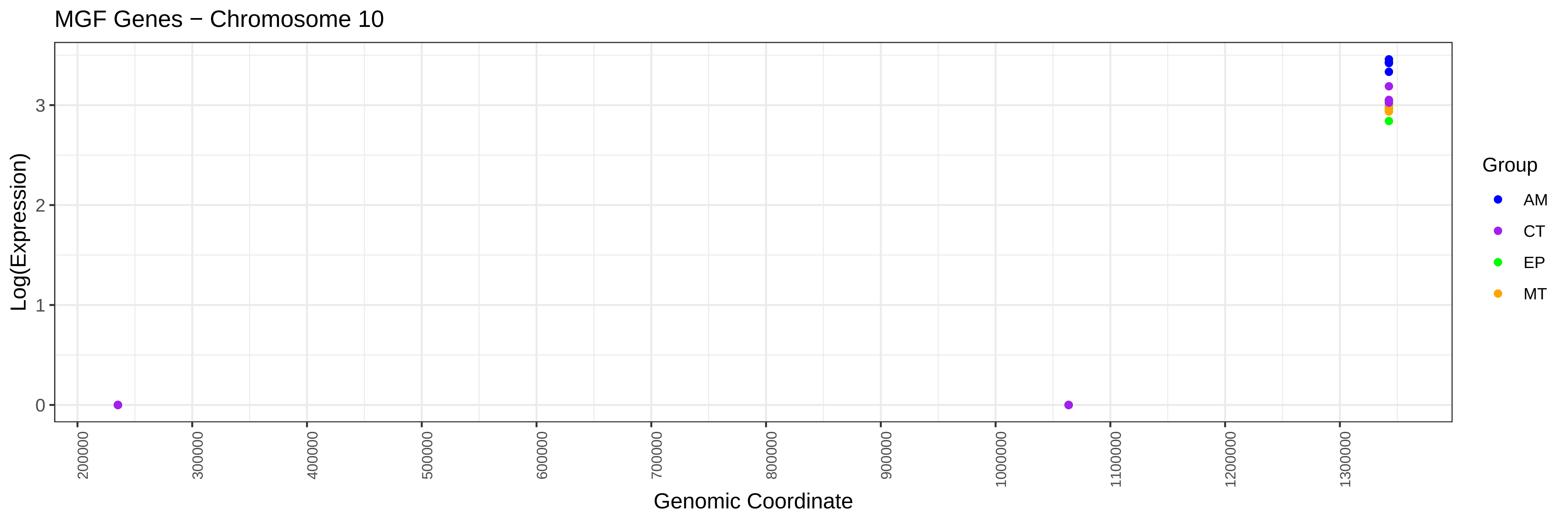
MGF Genes − Chromosome 10
Log(Expression)
0
1
2
3
200000
300000
400000
500000
600000
700000
800000
900000
1000000
1100000
1200000
1300000
Genomic Coordinate
Group
AM
CT
EP
MT

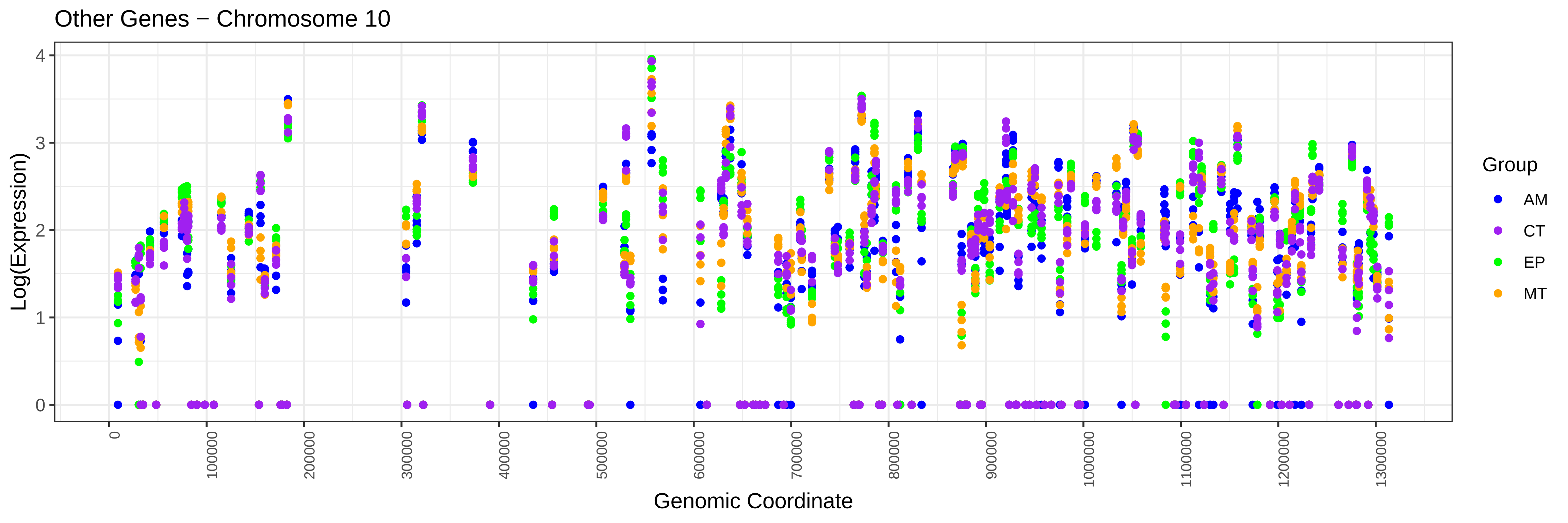
Other Genes − Chromosome 10
Log(Expression)
0
1
2
3
4
0
100000
200000
300000
400000
500000
600000
700000
800000
900000
1000000
1100000
1200000
1300000
Genomic Coordinate
Group
AM
CT
EP
MT

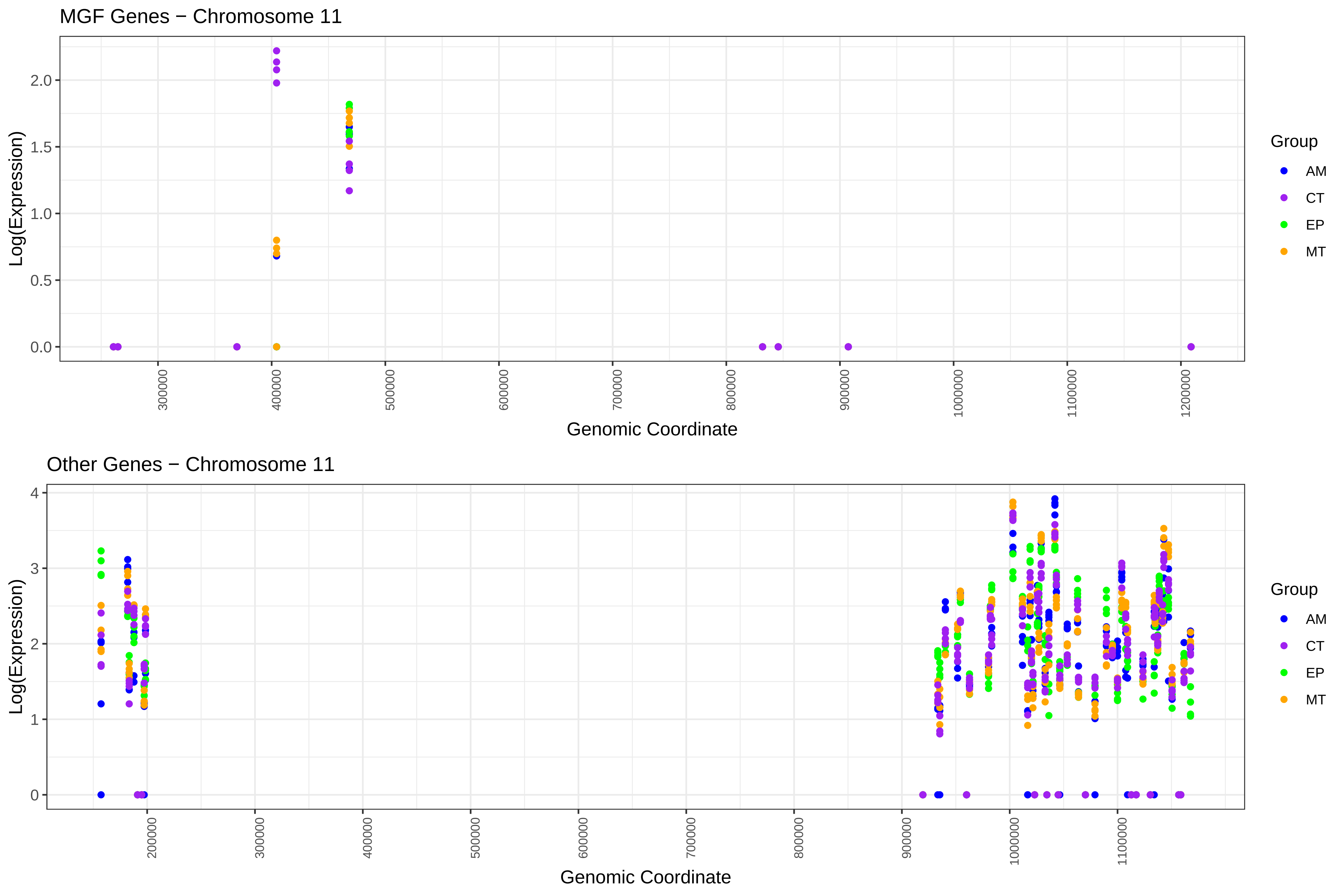
MGF Genes − Chromosome 11
Log(Expression)
2.0
1.5
1.0
0.5
0.0
300000
400000
500000
600000
700000
800000
900000
1000000
1100000
1200000
Genomic Coordinate
Group
AM
CT
EP
MT
Other Genes − Chromosome 11
Log(Expression)
4
3
2
1
0
200000
300000
400000
500000
600000
700000
800000
900000
1000000
1100000
Genomic Coordinate
Group
AM
CT
EP
MT

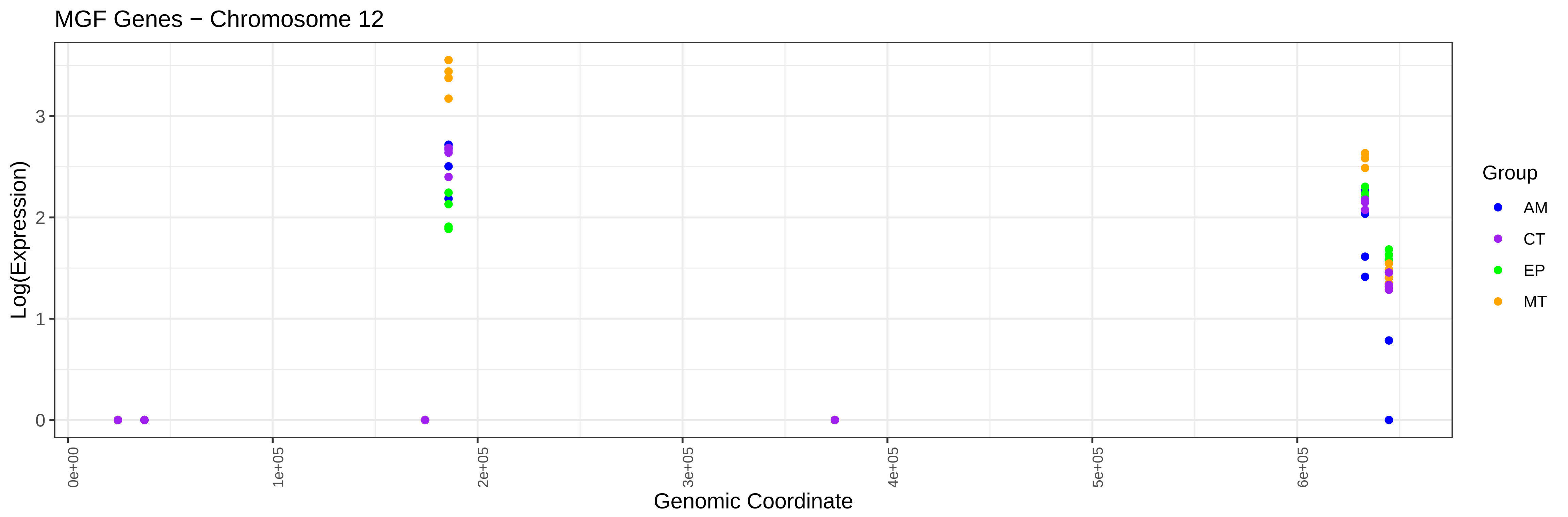
MGF Genes − Chromosome 12
Log(Expression)
0
1
2
3
0e+00
1e+05
2e+05
3e+05
4e+05
5e+05
6e+05
Genomic Coordinate
Group
AM
CT
EP
MT

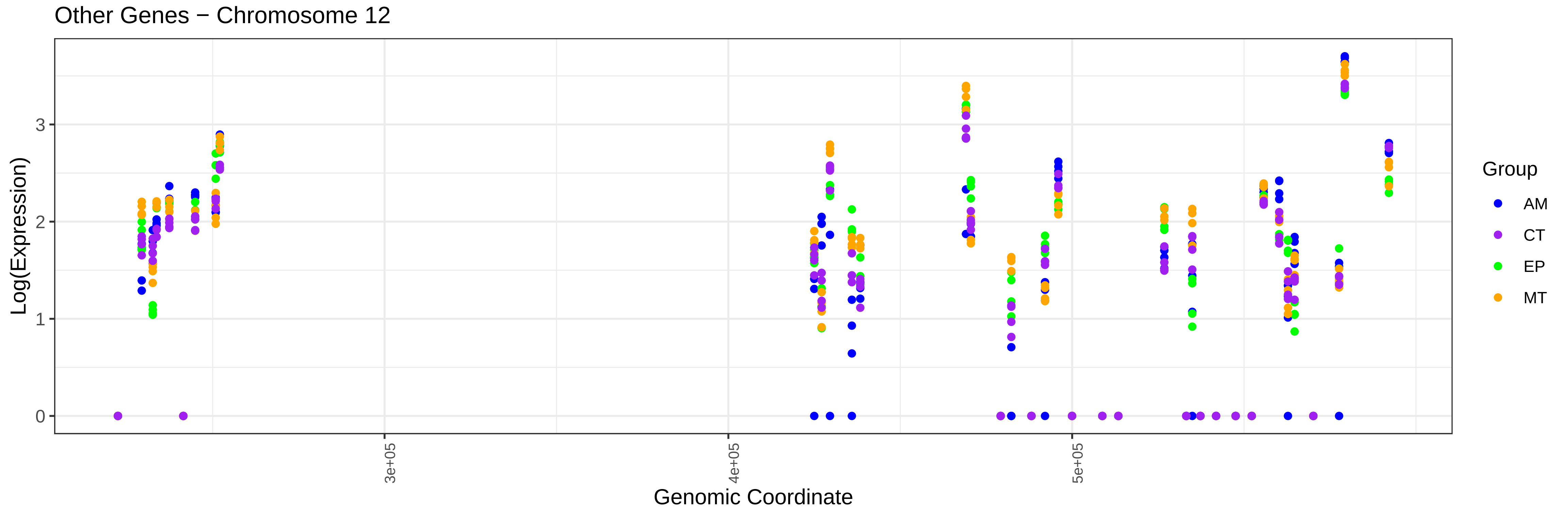
Other Genes − Chromosome 12
Log(Expression)
0
1
2
3
3e+05
4e+05
5e+05
Genomic Coordinate
Group
AM
CT
EP
MT

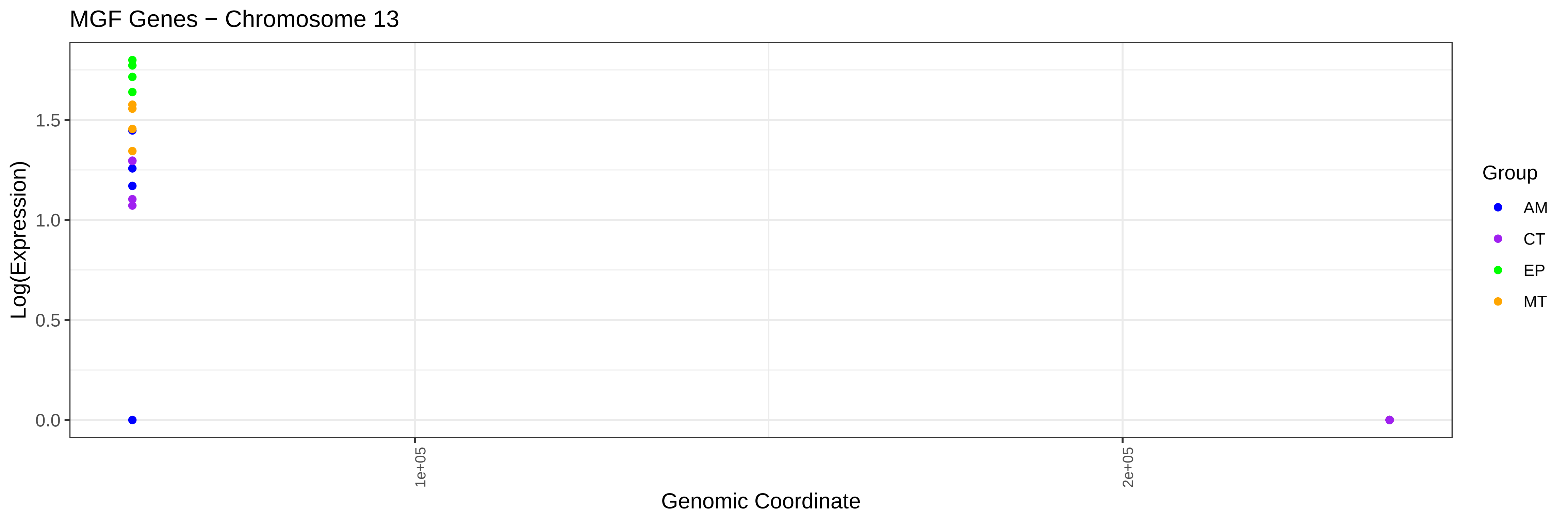

MGF Genes − Chromosome 13
Log(Expression)
0.0
0.5
1.0
1.5
1e+05
2e+05
Genomic Coordinate
Group
AM
CT
EP
MT


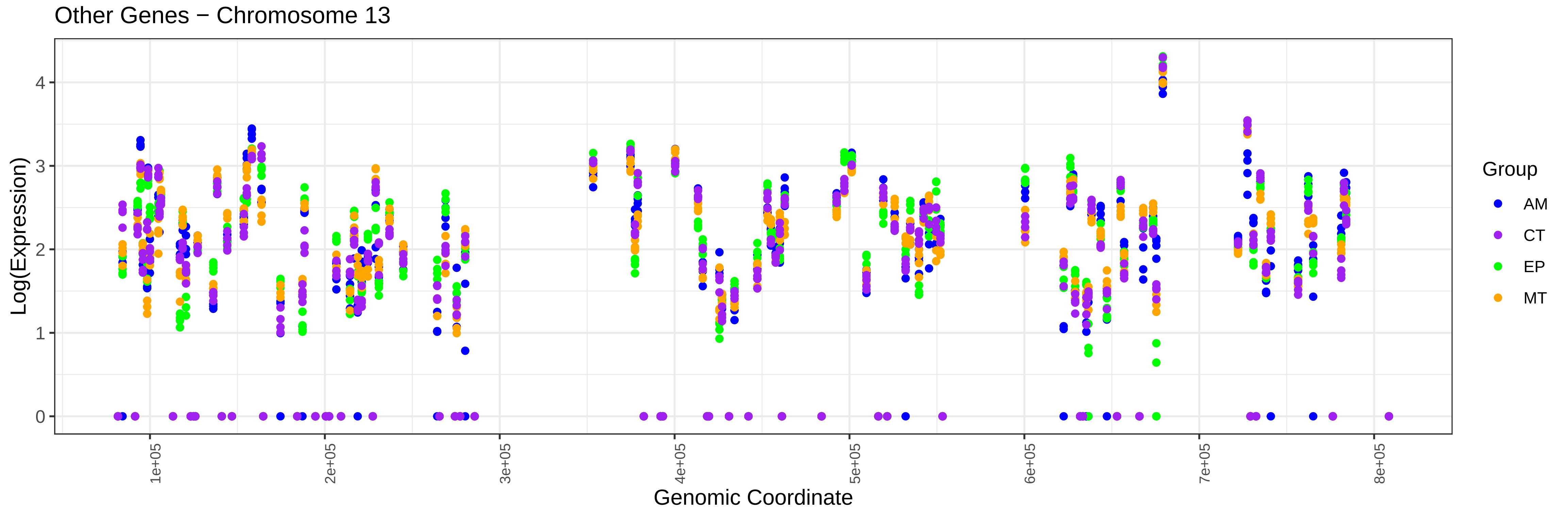

Other Genes − Chromosome 13
Log(Expression)
0
1
2
3
4
1e+05
2e+05
3e+05
4e+05
5e+05
6e+05
7e+05
8e+05
Genomic Coordinate
Group
AM
CT
EP
MT

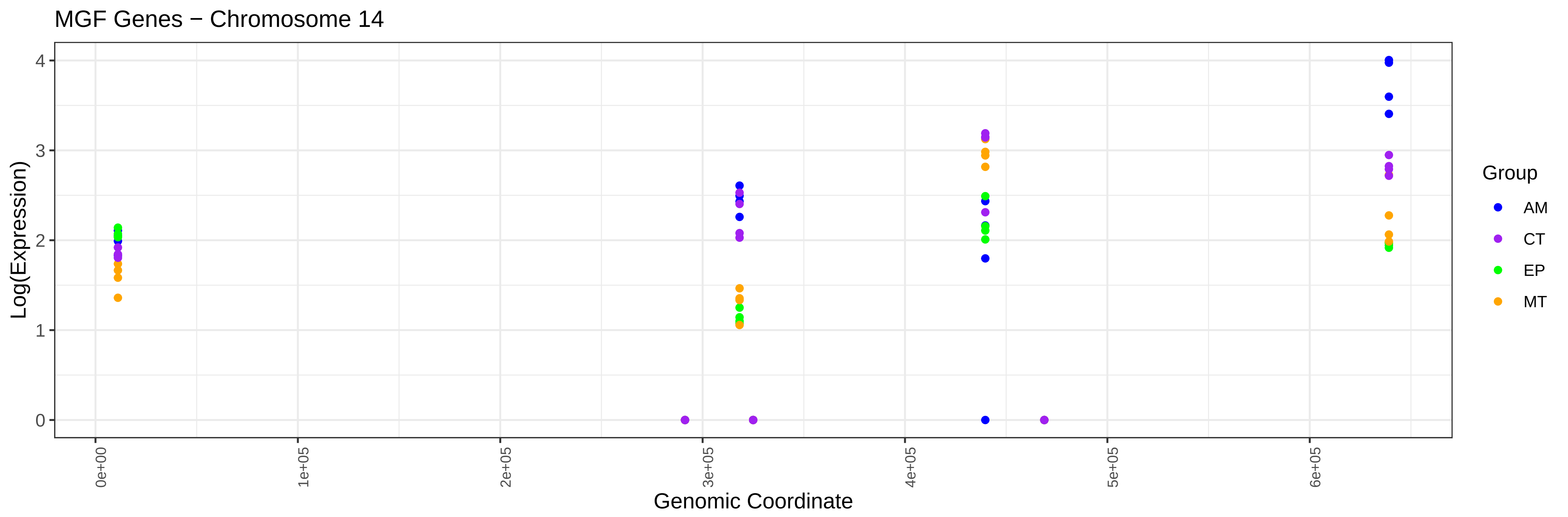
MGF Genes − Chromosome 14
Log(Expression)
0
1
2
3
4
0e+00
1e+05
2e+05
3e+05
4e+05
5e+05
6e+05
Genomic Coordinate
Group
AM
CT
EP
MT

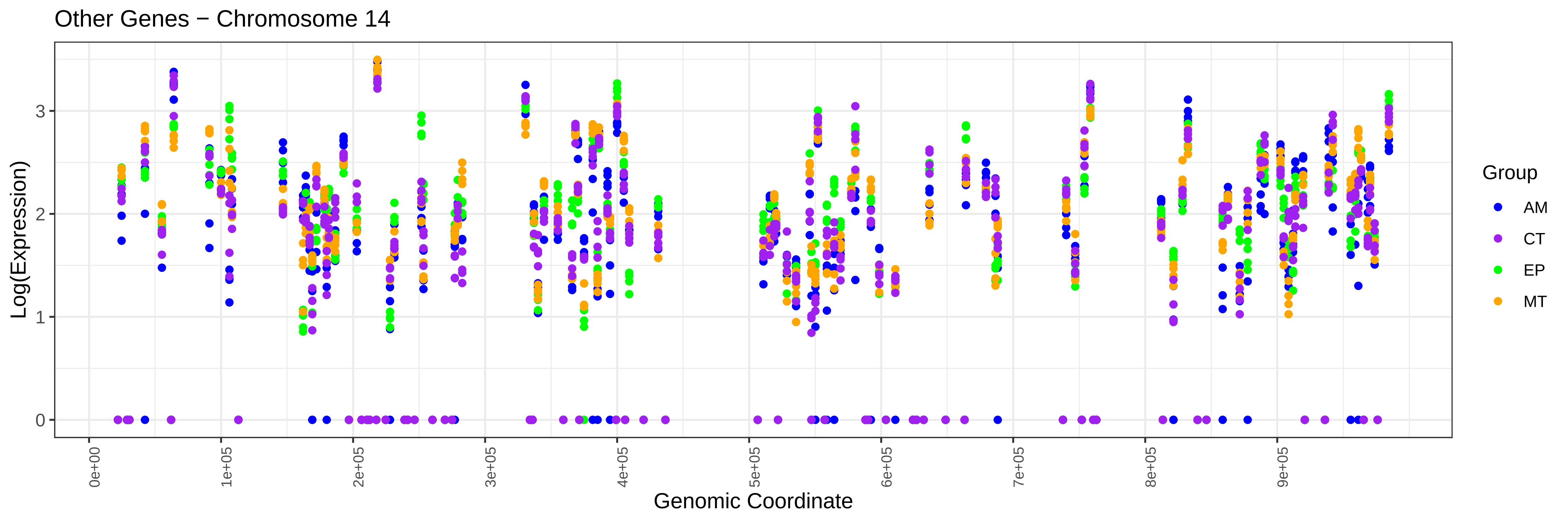
Other Genes − Chromosome 14
Log(Expression)
0
1
2
3
0e+00
1e+05
2e+05
3e+05
4e+05
5e+05
6e+05
7e+05
8e+05
9e+05
Genomic Coordinate
Group
AM
CT
EP
MT

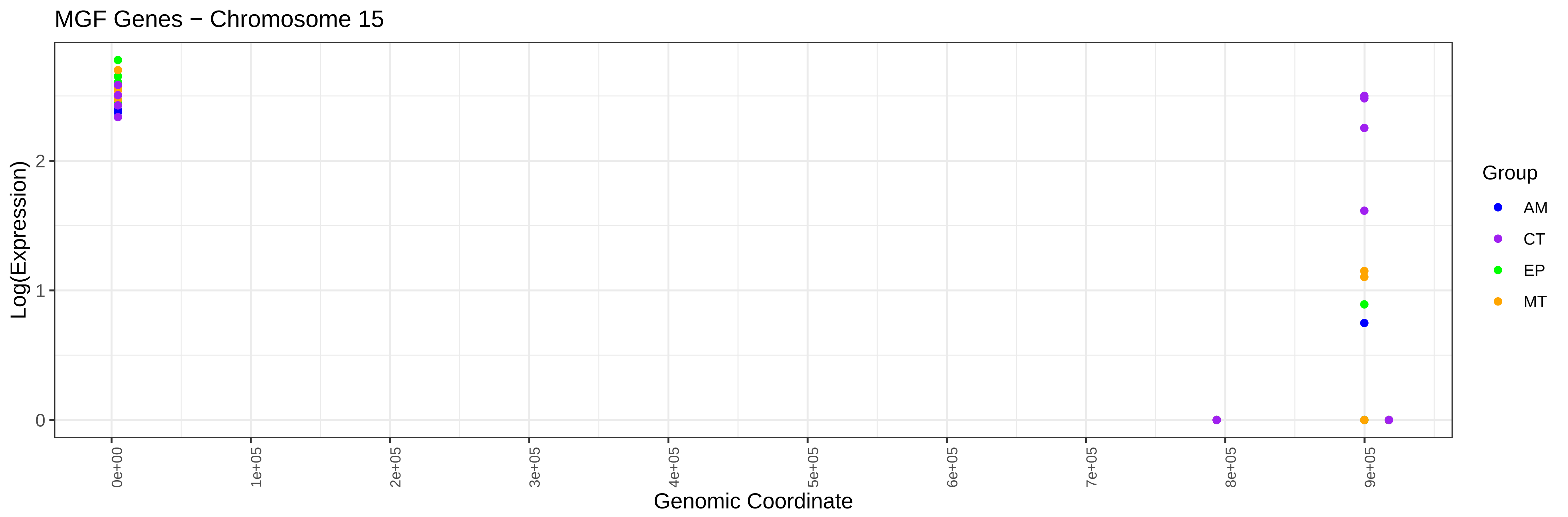
MGF Genes − Chromosome 15
Log(Expression)
0
1
2
0e+00
1e+05
2e+05
3e+05
4e+05
5e+05
6e+05
7e+05
8e+05
9e+05
Genomic Coordinate
Group
AM
CT
EP
MT

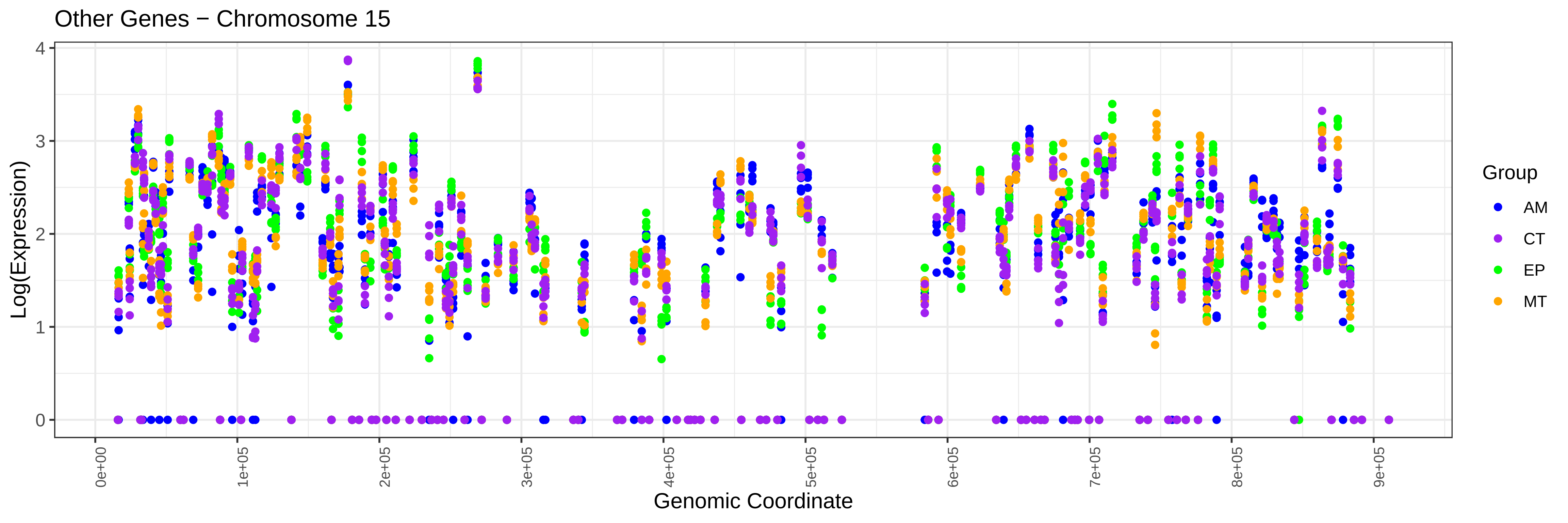
Other Genes − Chromosome 15
Log(Expression)
0
1
2
3
4
0e+00
1e+05
2e+05
3e+05
4e+05
5e+05
6e+05
7e+05
8e+05
9e+05
Genomic Coordinate
Group
AM
CT
EP
MT

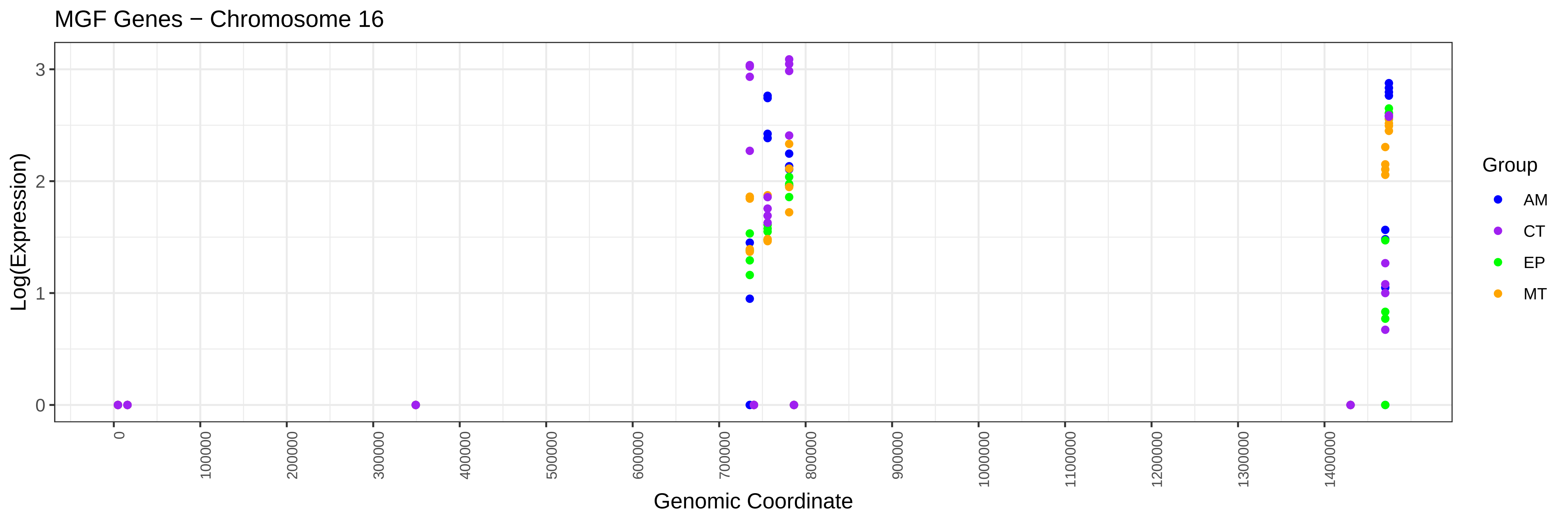
MGF Genes − Chromosome 16
Log(Expression)
0
1
2
3
0
100000
200000
300000
400000
500000
600000
700000
800000
900000
1000000
1100000
1200000
1300000
1400000
Genomic Coordinate
Group
AM
CT
EP
MT

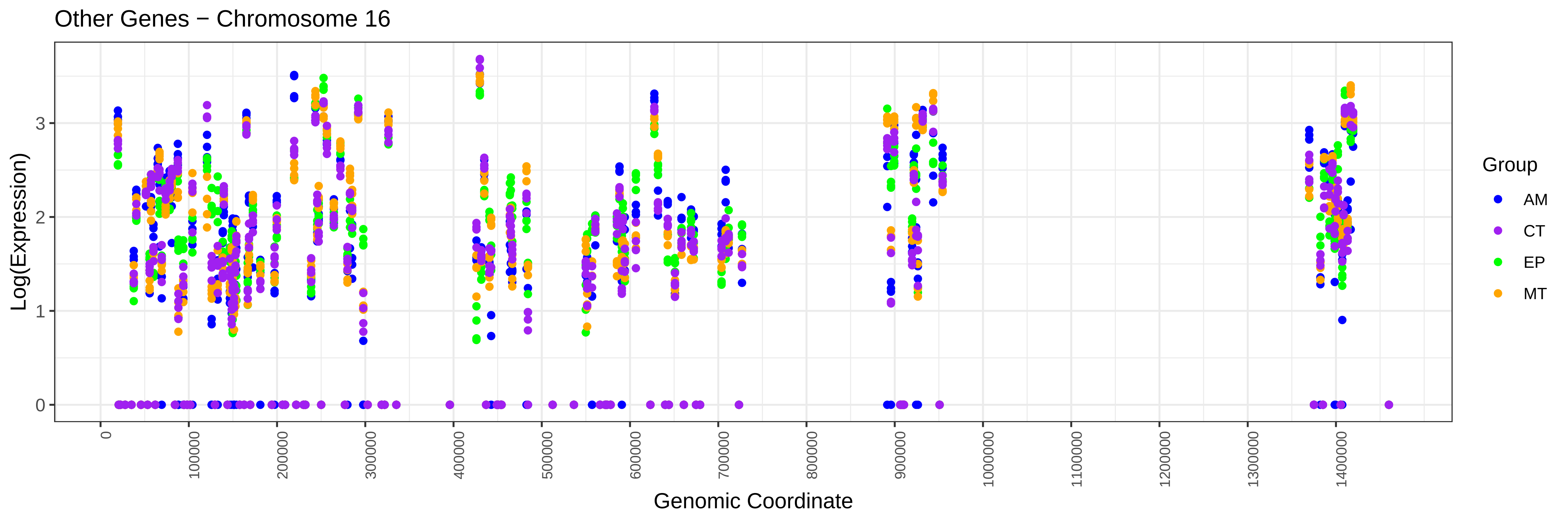
Other Genes − Chromosome 16
Log(Expression)
0
1
2
3
0
100000
200000
300000
400000
500000
600000
700000
800000
900000
1000000
1100000
1200000
1300000
1400000
Genomic Coordinate
Group
AM
CT
EP
MT

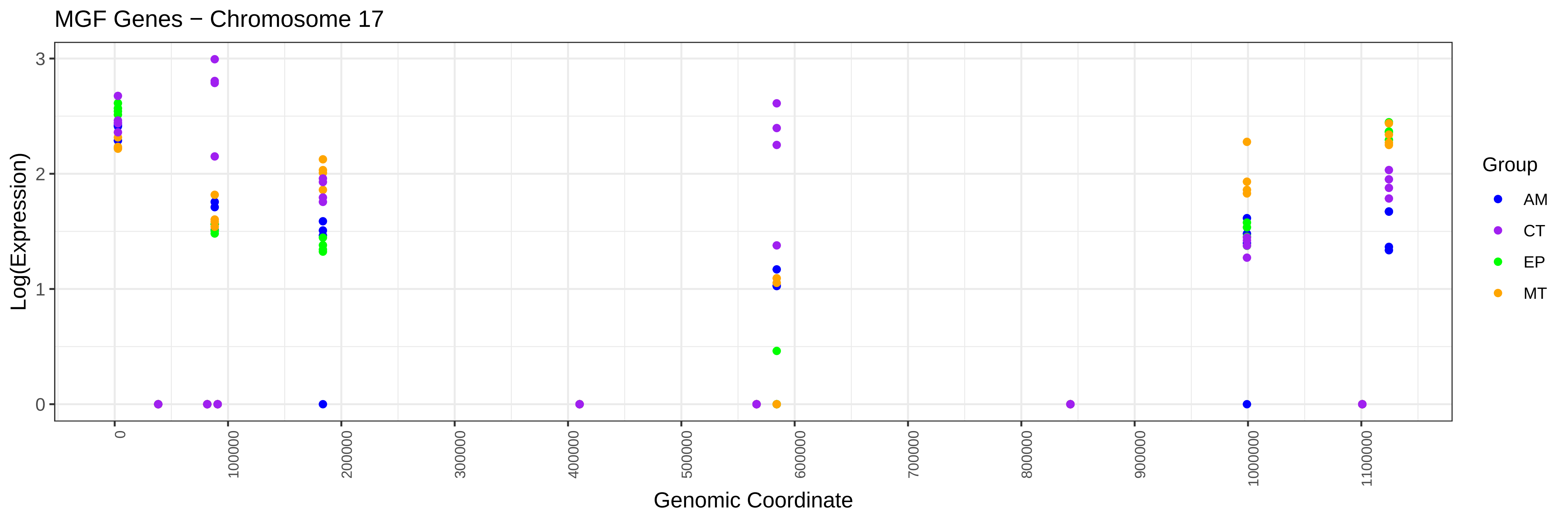
MGF Genes − Chromosome 17
Log(Expression)
0
1
2
3
0
100000
200000
300000
400000
500000
600000
700000
800000
900000
1000000
1100000
Genomic Coordinate
Group
AM
CT
EP
MT

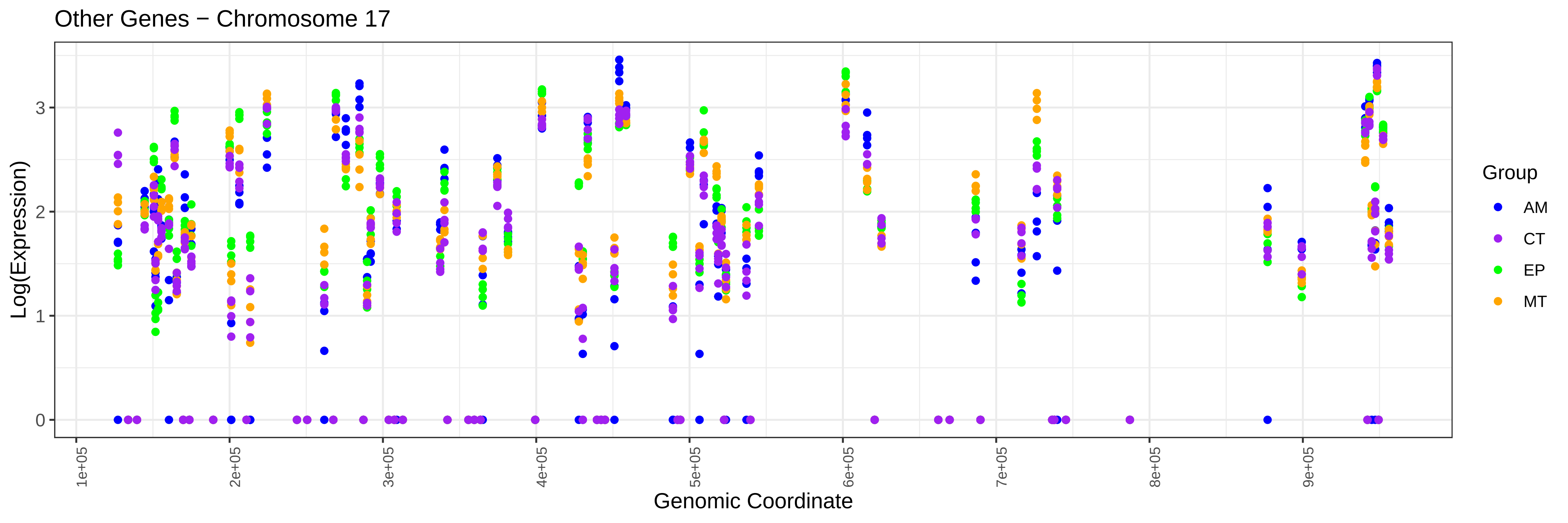
Other Genes − Chromosome 17
Log(Expression)
0
1
2
3
1e+05
2e+05
3e+05
4e+05
5e+05
6e+05
7e+05
8e+05
9e+05
Genomic Coordinate
Group
AM
CT
EP
MT

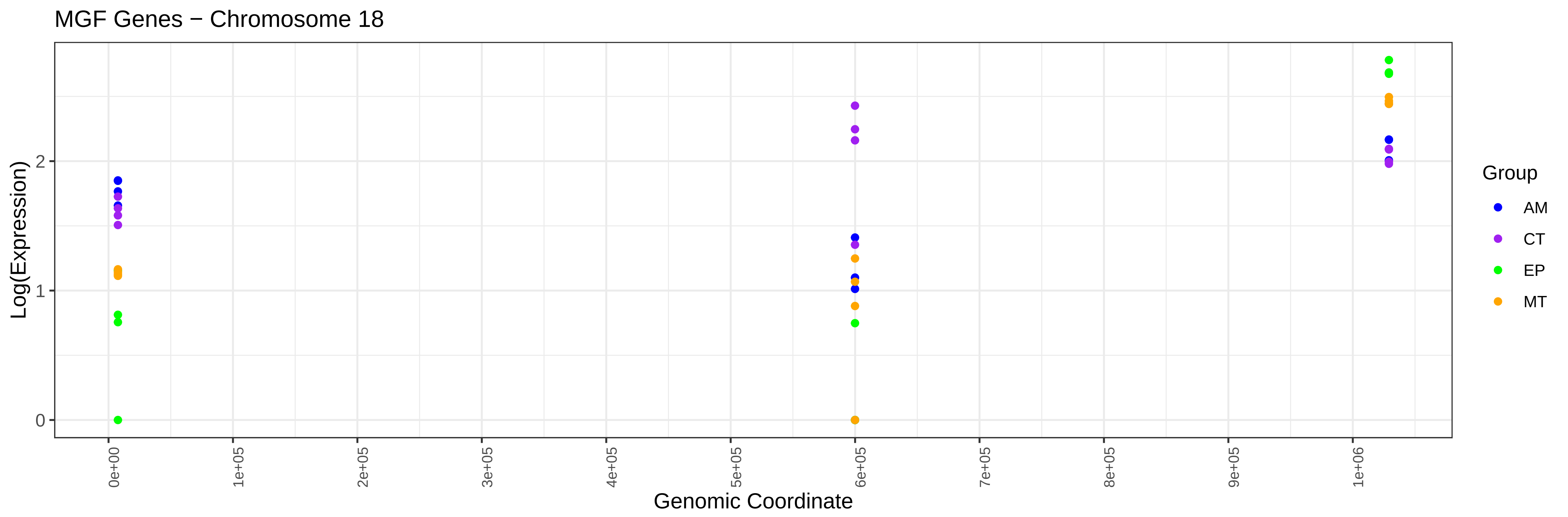
MGF Genes − Chromosome 18
Log(Expression)
0
1
2
0e+00
1e+05
2e+05
3e+05
4e+05
5e+05
6e+05
7e+05
8e+05
9e+05
1e+06
Genomic Coordinate
Group
AM
CT
EP
MT

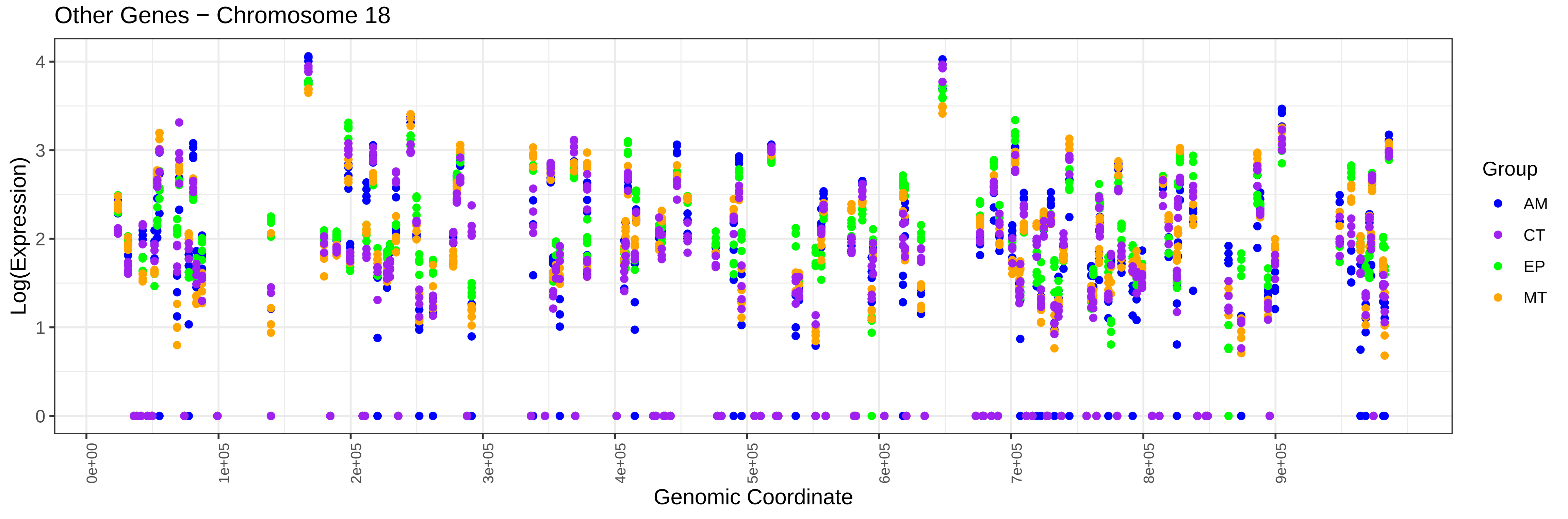
Other Genes − Chromosome 18
Log(Expression)
0
1
2
3
4
0e+00
1e+05
2e+05
3e+05
4e+05
5e+05
6e+05
7e+05
8e+05
9e+05
Genomic Coordinate
Group
AM
CT
EP
MT

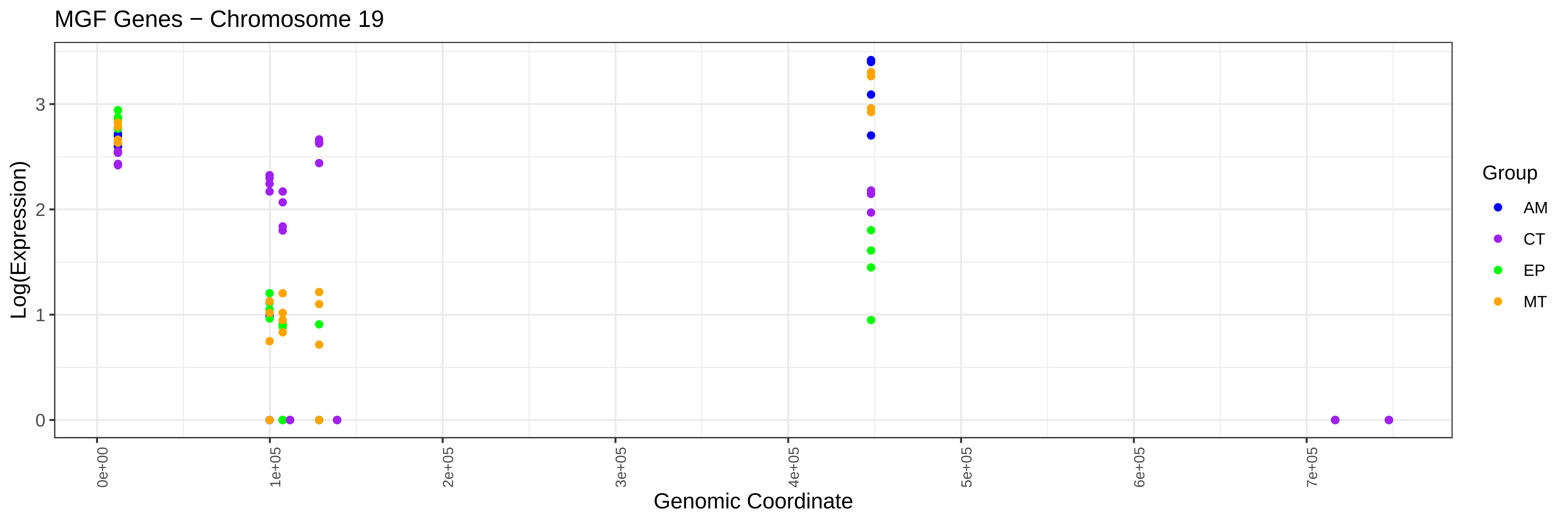
MGF Genes − Chromosome 19
Log(Expression)
Genomic Coordinate
0
1
2
3
0e+00
1e+05
2e+05
3e+05
4e+05
5e+05
6e+05
7e+05
Group
AM
CT
EP
MT

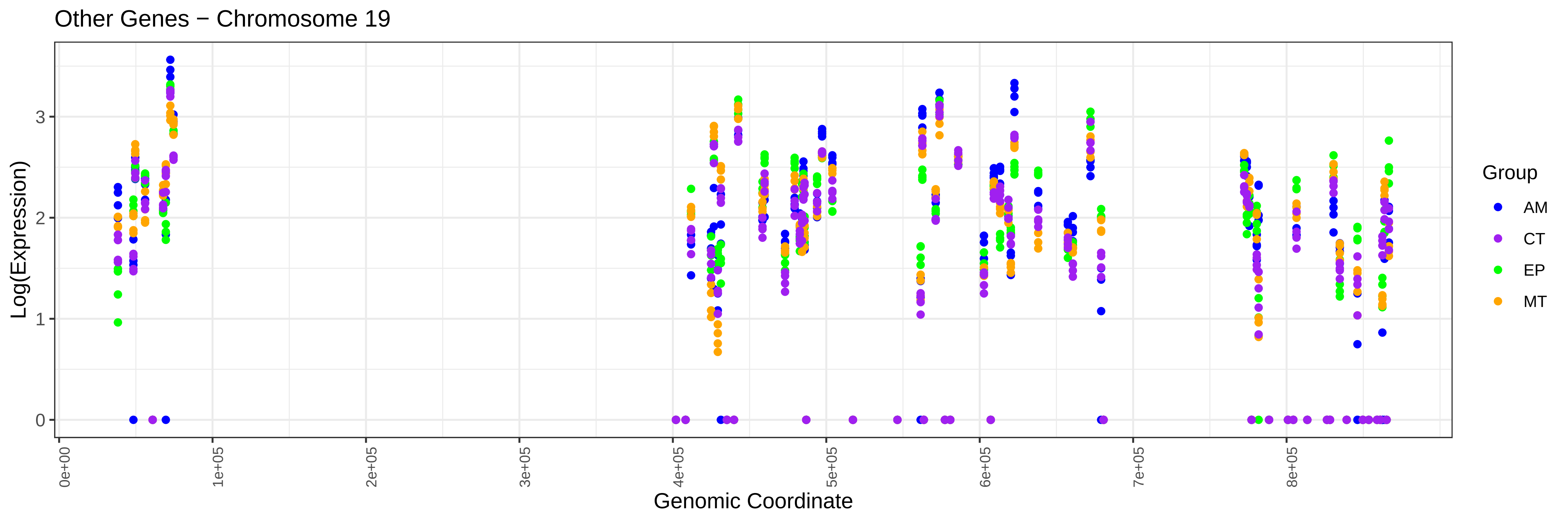
Other Genes − Chromosome 19
Log(Expression)
Genomic Coordinate
0
1
2
3
0e+00
1e+05
2e+05
3e+05
4e+05
5e+05
6e+05
7e+05
8e+05
Group
AM
CT
EP
MT

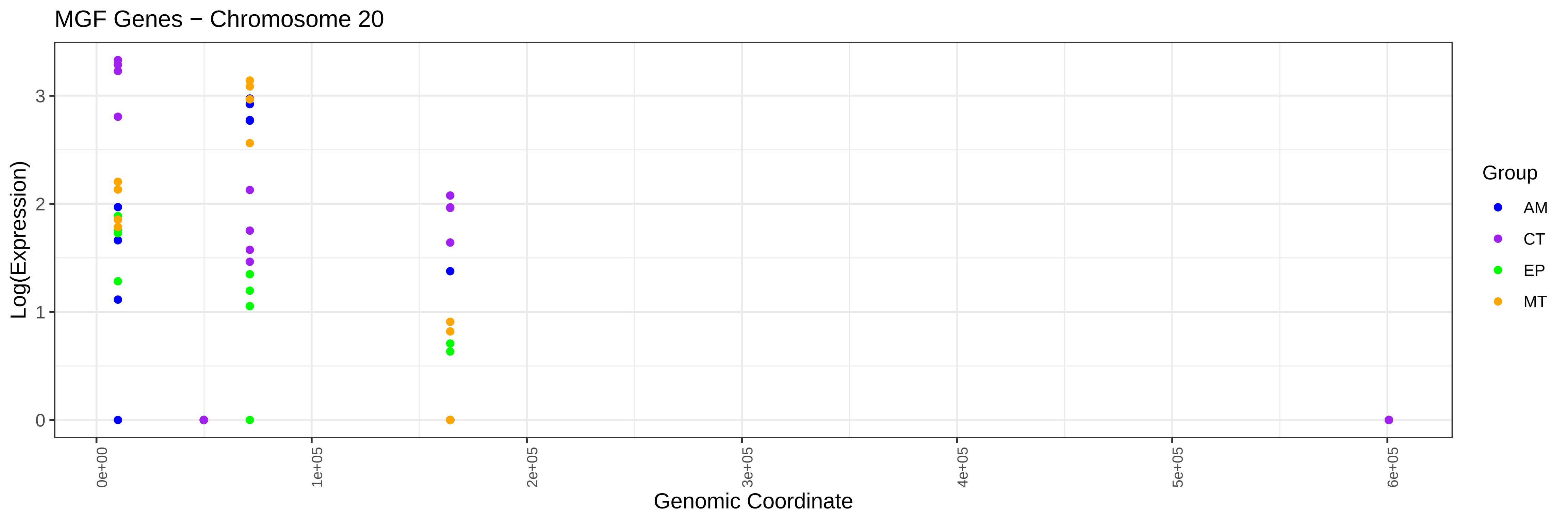
MGF Genes − Chromosome 20
Log(Expression)
0
1
2
3
0e+00
1e+05
2e+05
3e+05
4e+05
5e+05
6e+05
Genomic Coordinate
Group
AM
CT
EP
MT

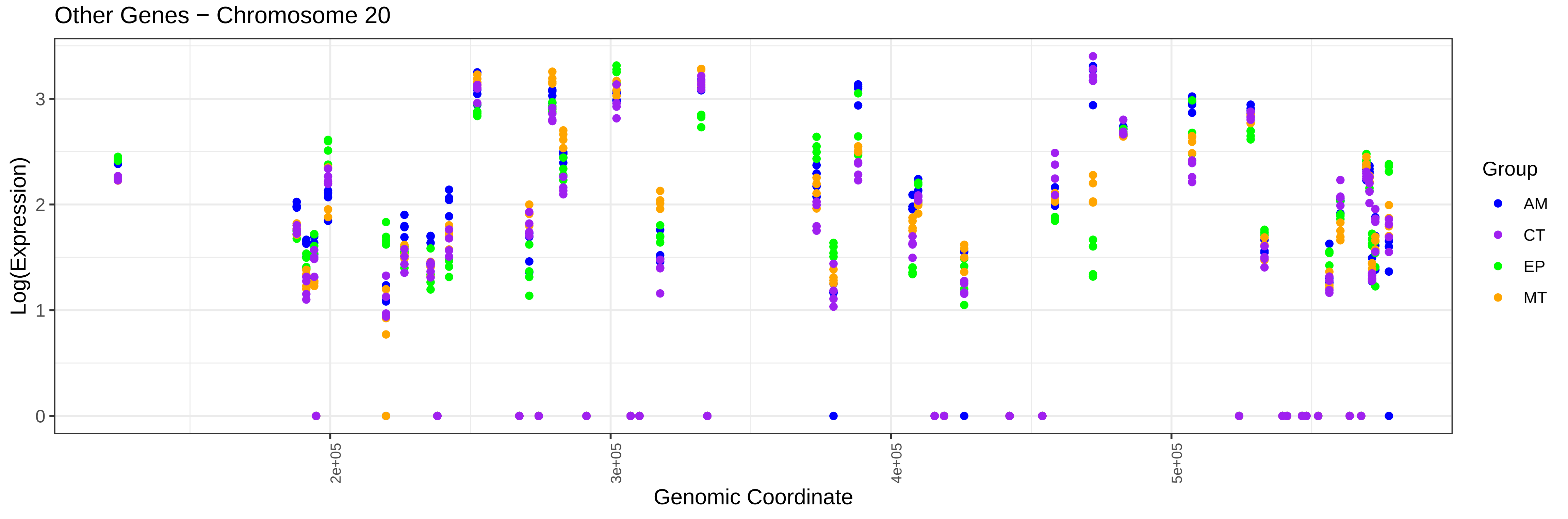
Other Genes − Chromosome 20
Log(Expression)
0
1
2
3
2e+05
3e+05
4e+05
5e+05
Genomic Coordinate
Group
AM
CT
EP
MT

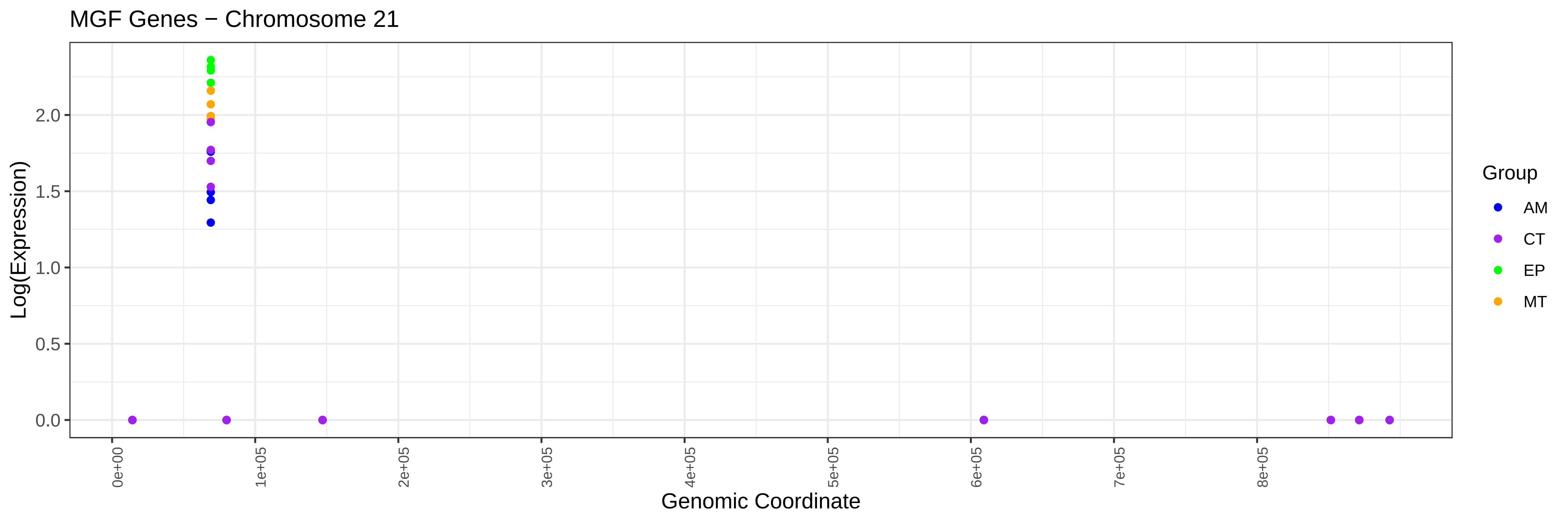
MGF Genes − Chromosome 21
Log(Expression)
Genomic Coordinate
0.0
0.5
1.0
1.5
2.0
0e+00
1e+05
2e+05
3e+05
4e+05
5e+05
6e+05
7e+05
8e+05
Group
AM
CT
EP
MT

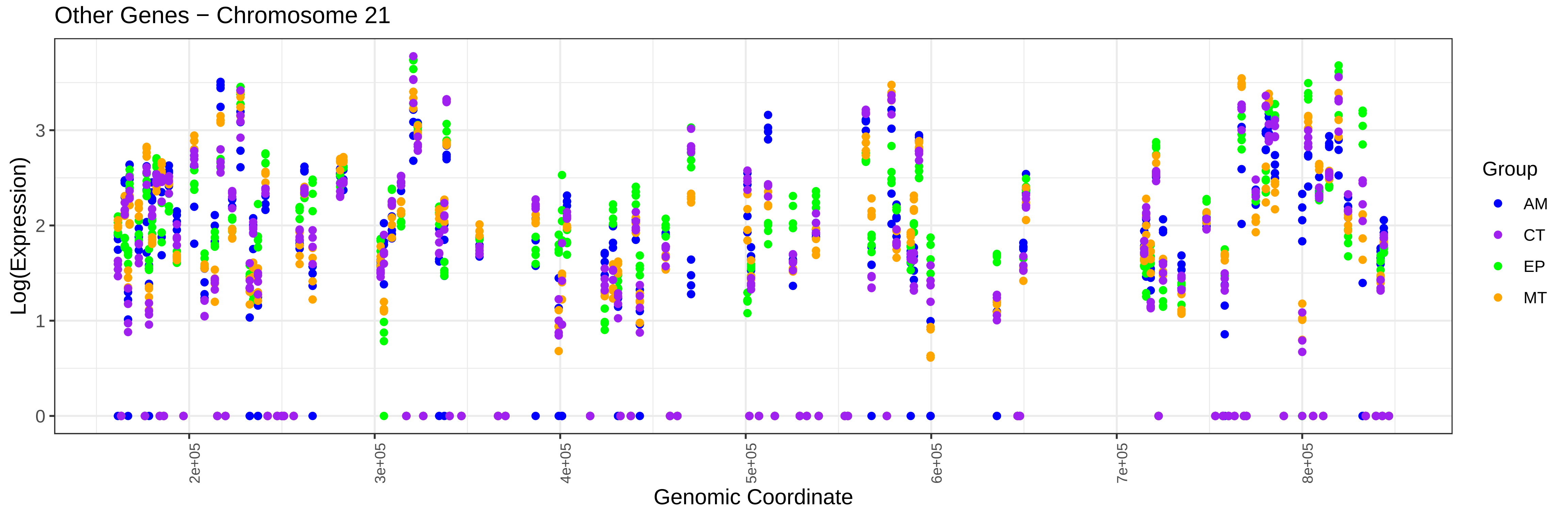
Other Genes − Chromosome 21
Log(Expression)
Genomic Coordinate
0
1
2
3
2e+05
3e+05
4e+05
5e+05
6e+05
7e+05
8e+05
Group
AM
CT
EP
MT

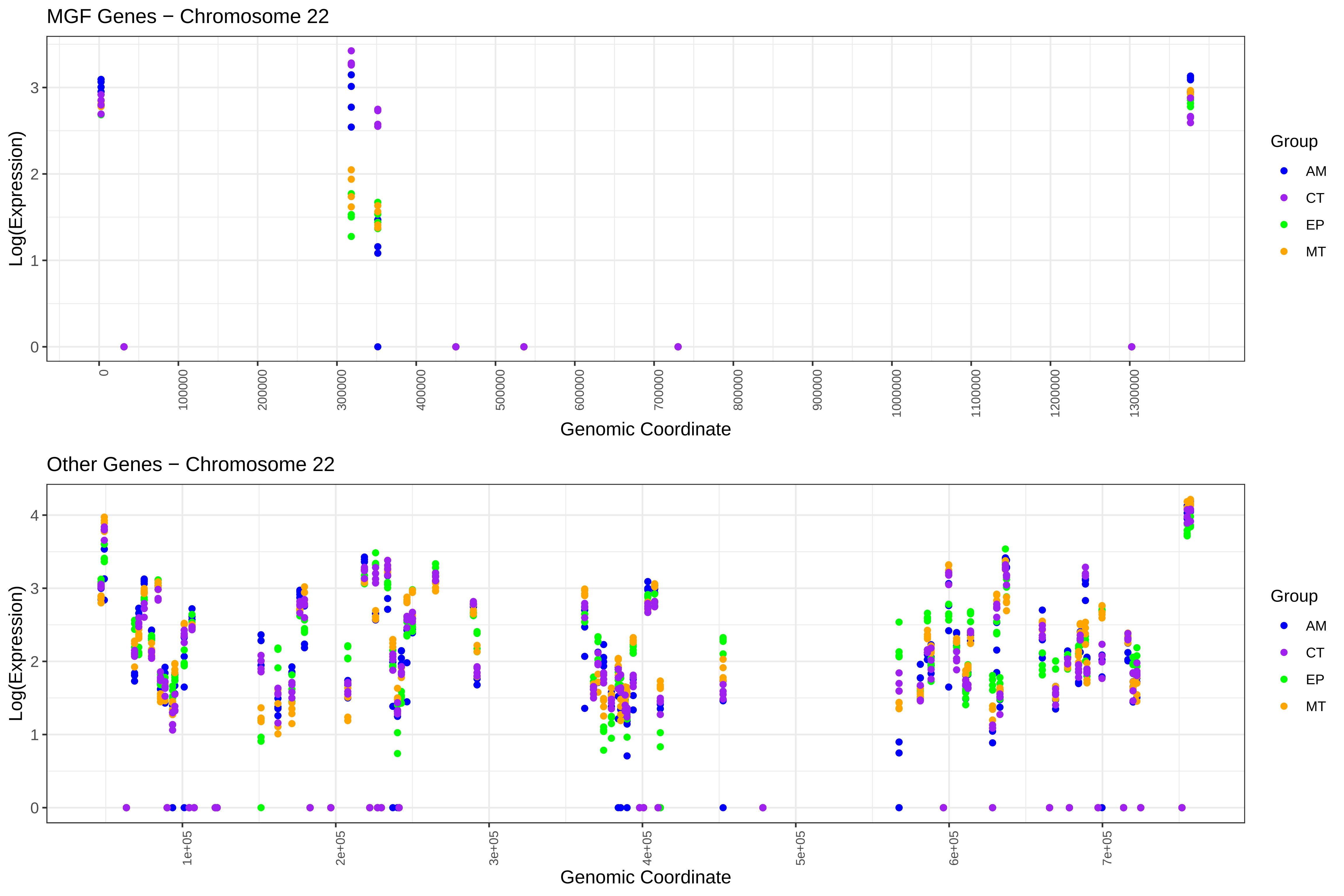

MGF Genes − Chromosome 22
Log(Expression)
Genomic Coordinate
Group
AM
CT
EP
MT
Other Genes − Chromosome 22
Log(Expression)
Genomic Coordinate
Group
AM
CT
EP
MT

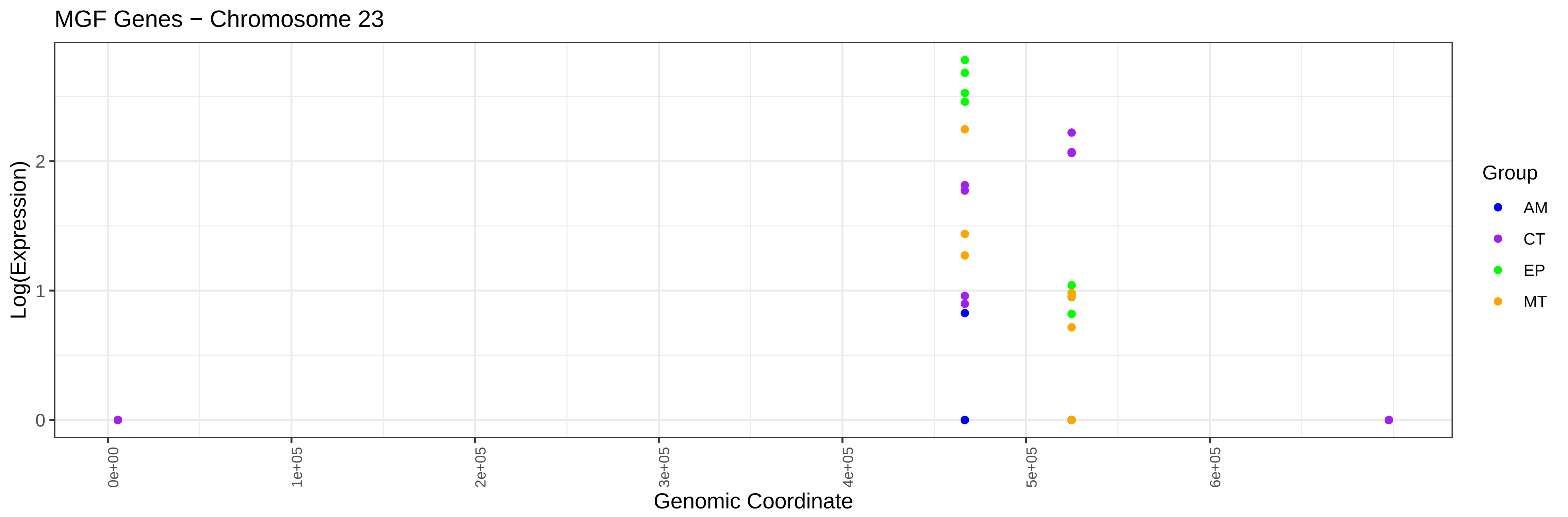
MGF Genes − Chromosome 23
Log(Expression)
Genomic Coordinate
0e+00
1e+05
2e+05
3e+05
4e+05
5e+05
6e+05
0
1
2
Group
AM
CT
EP
MT

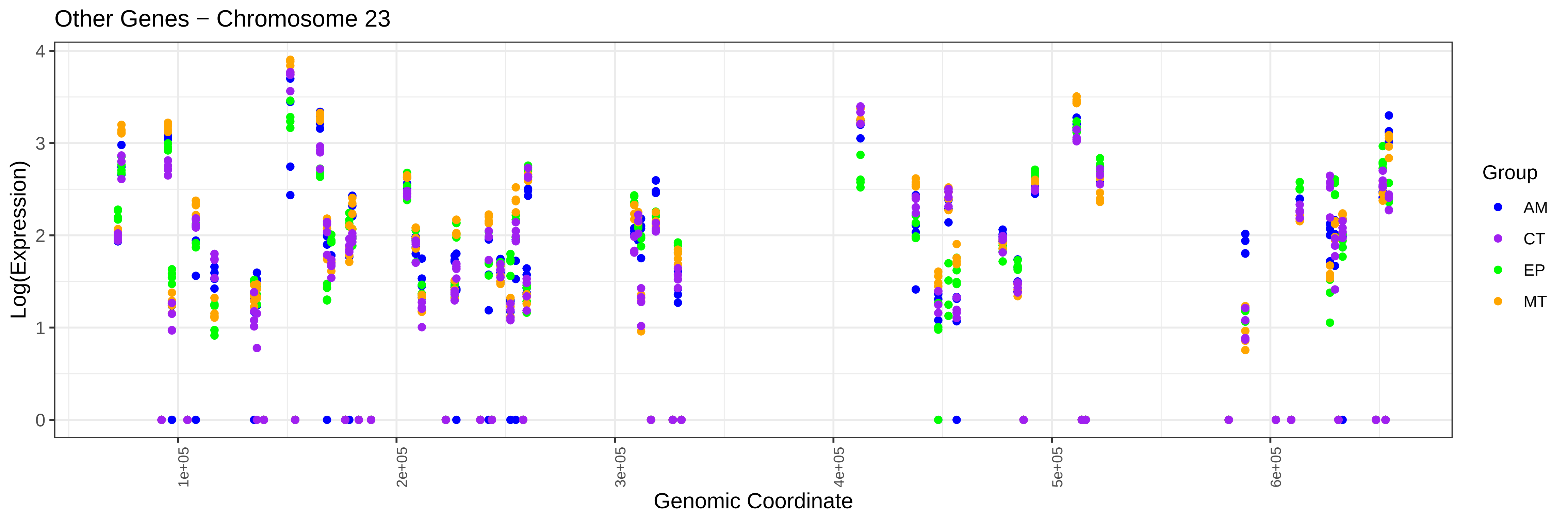
Other Genes − Chromosome 23
Log(Expression)
Genomic Coordinate
1e+05
2e+05
3e+05
4e+05
5e+05
6e+05
0
1
2
3
4
Group
AM
CT
EP
MT

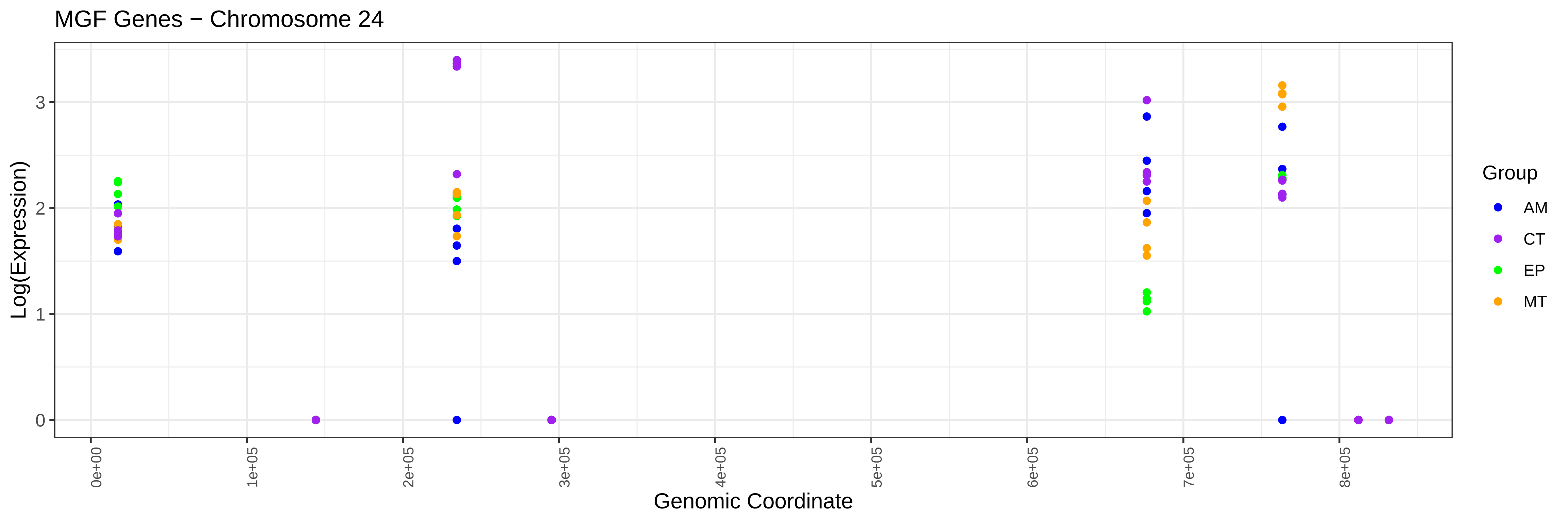
MGF Genes − Chromosome 24
Log(Expression)
Genomic Coordinate
0e+00
1e+05
2e+05
3e+05
4e+05
5e+05
6e+05
7e+05
8e+05
0
1
2
3
Group
AM
CT
EP
MT

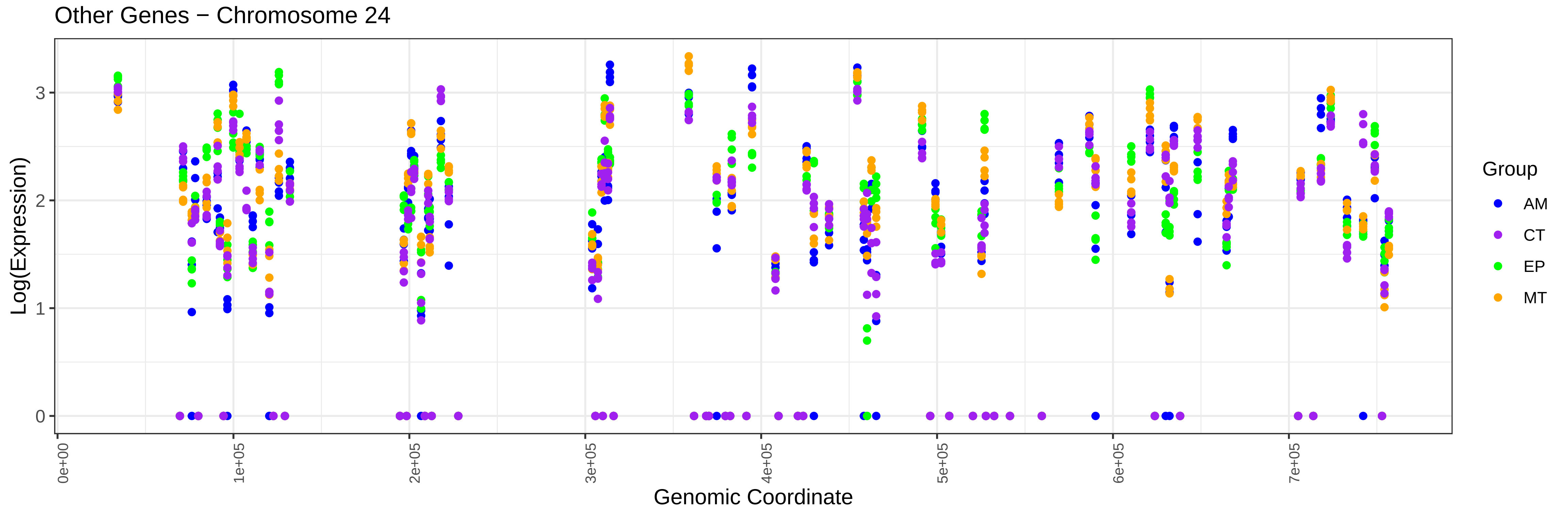
Other Genes − Chromosome 24
Log(Expression)
Genomic Coordinate
0e+00
1e+05
2e+05
3e+05
4e+05
5e+05
6e+05
7e+05
0
1
2
3
Group
AM
CT
EP
MT

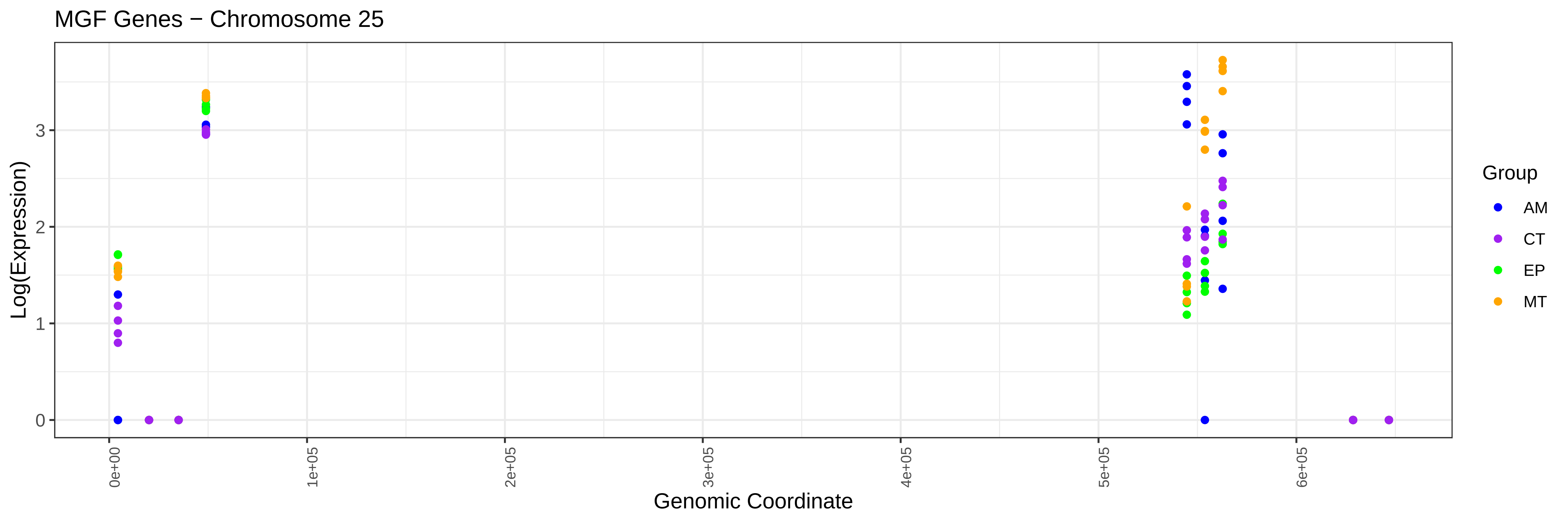
MGF Genes − Chromosome 25
Log(Expression)
0
1
2
3
0e+00
1e+05
2e+05
3e+05
4e+05
5e+05
6e+05
Genomic Coordinate
Group
AM
CT
EP
MT

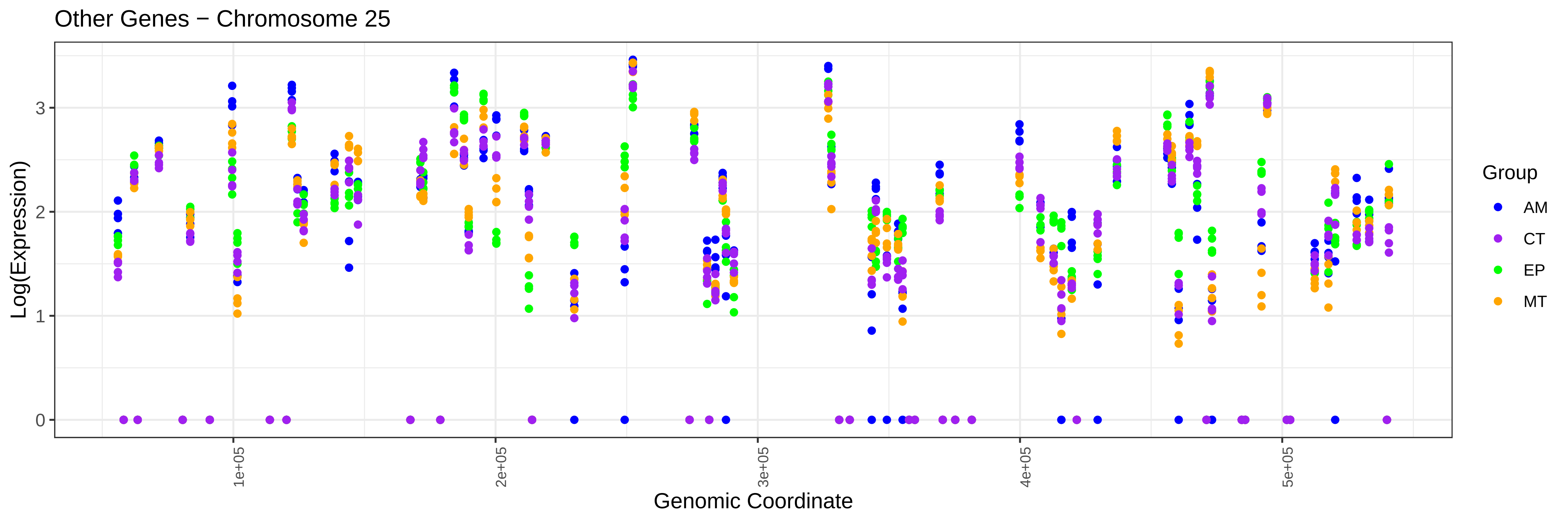
Other Genes − Chromosome 25
Log(Expression)
0
1
2
3
1e+05
2e+05
3e+05
4e+05
5e+05
Genomic Coordinate
Group
AM
CT
EP
MT

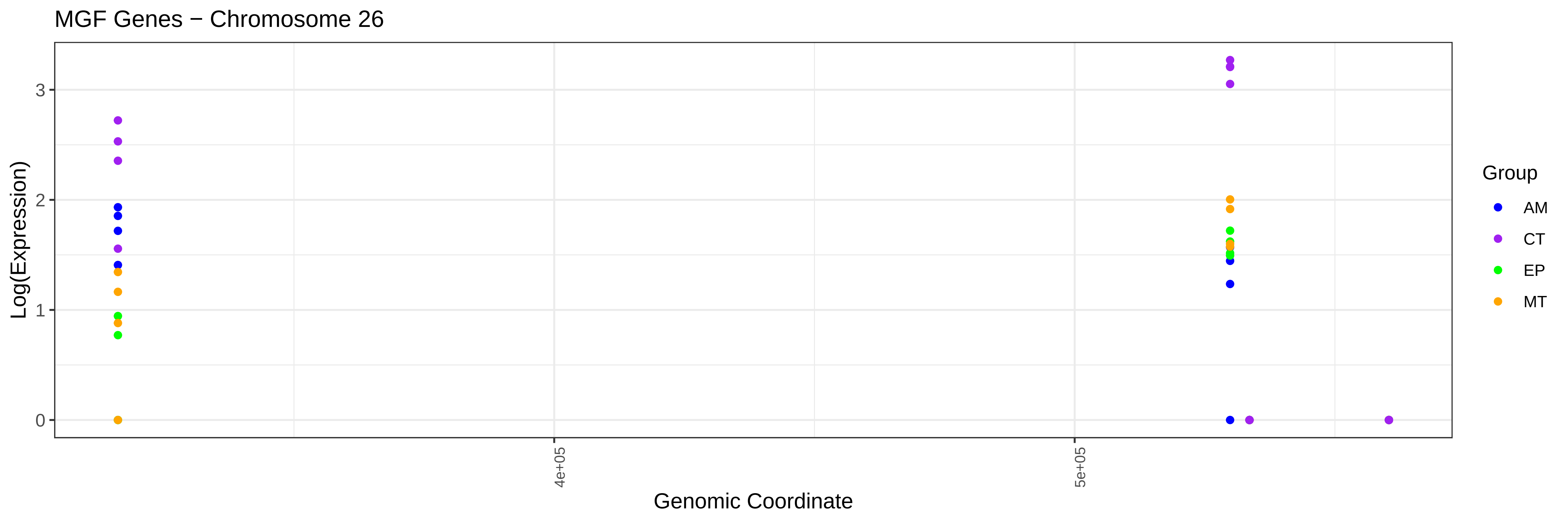
MGF Genes − Chromosome 26
Log(Expression)
Genomic Coordinate
Group
AM
CT
EP
MT

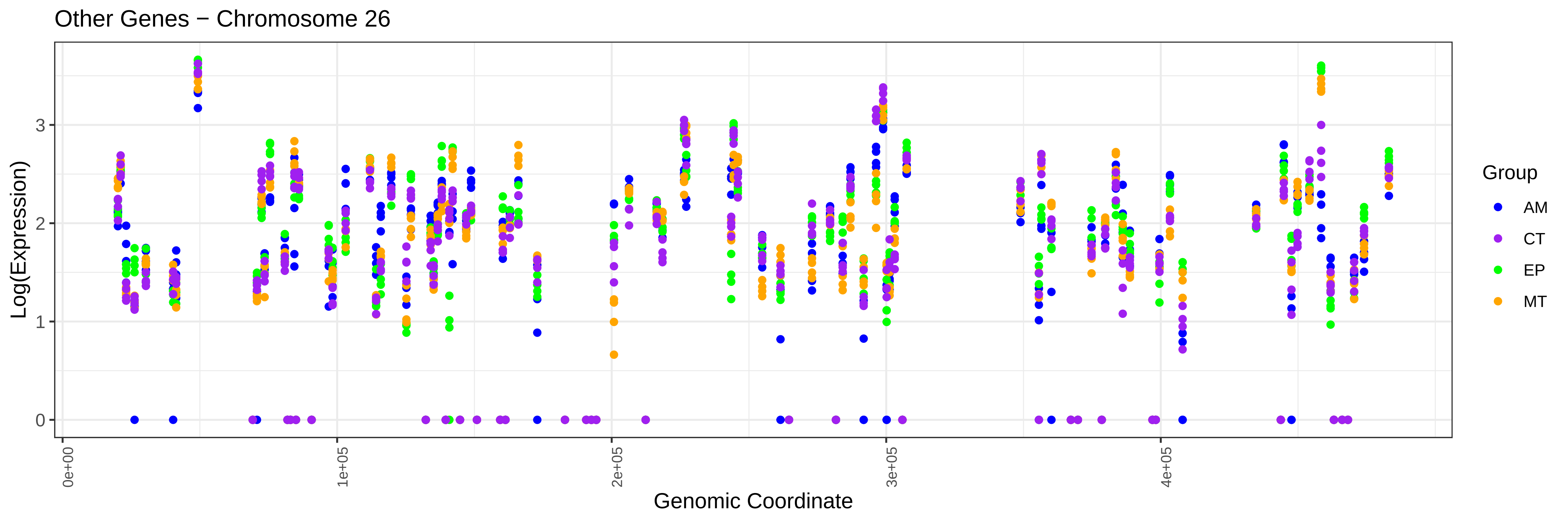
Other Genes − Chromosome 26
Log(Expression)
Genomic Coordinate
Group
AM
CT
EP
MT

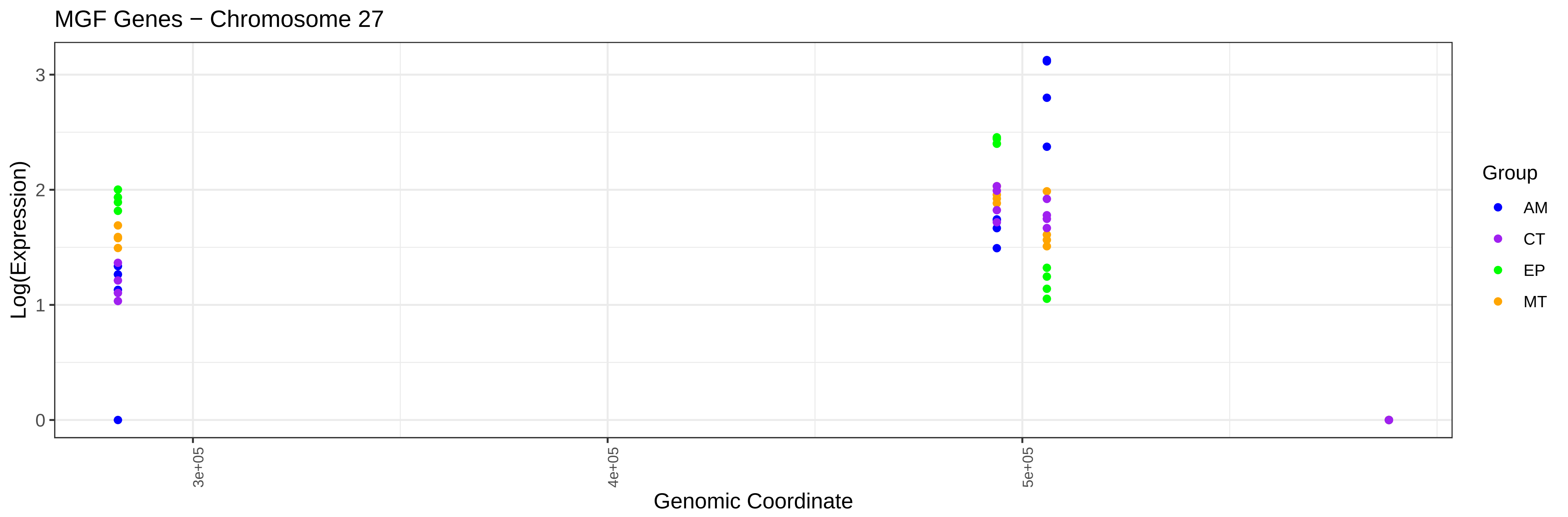

MGF Genes − Chromosome 27
Log(Expression)
0
1
2
3
3e+05
4e+05
5e+05
Genomic Coordinate
Group
AM
CT
EP
MT


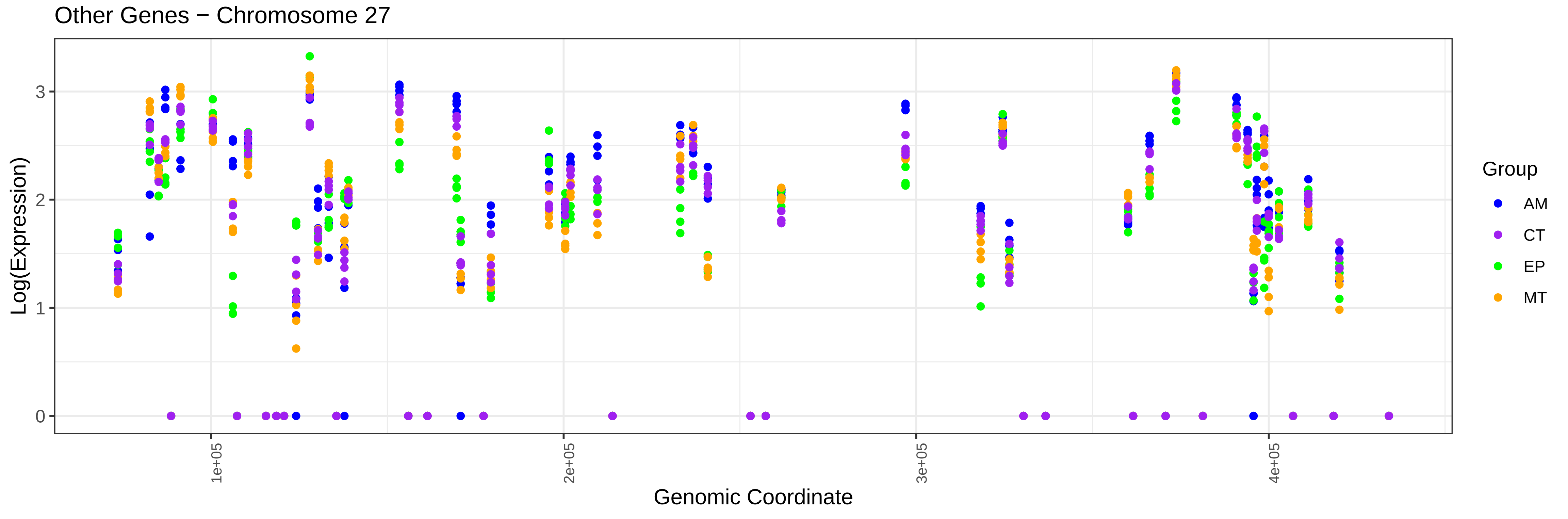

Other Genes − Chromosome 27
Log(Expression)
0
1
2
3
1e+05
2e+05
3e+05
4e+05
Genomic Coordinate
Group
AM
CT
EP
MT

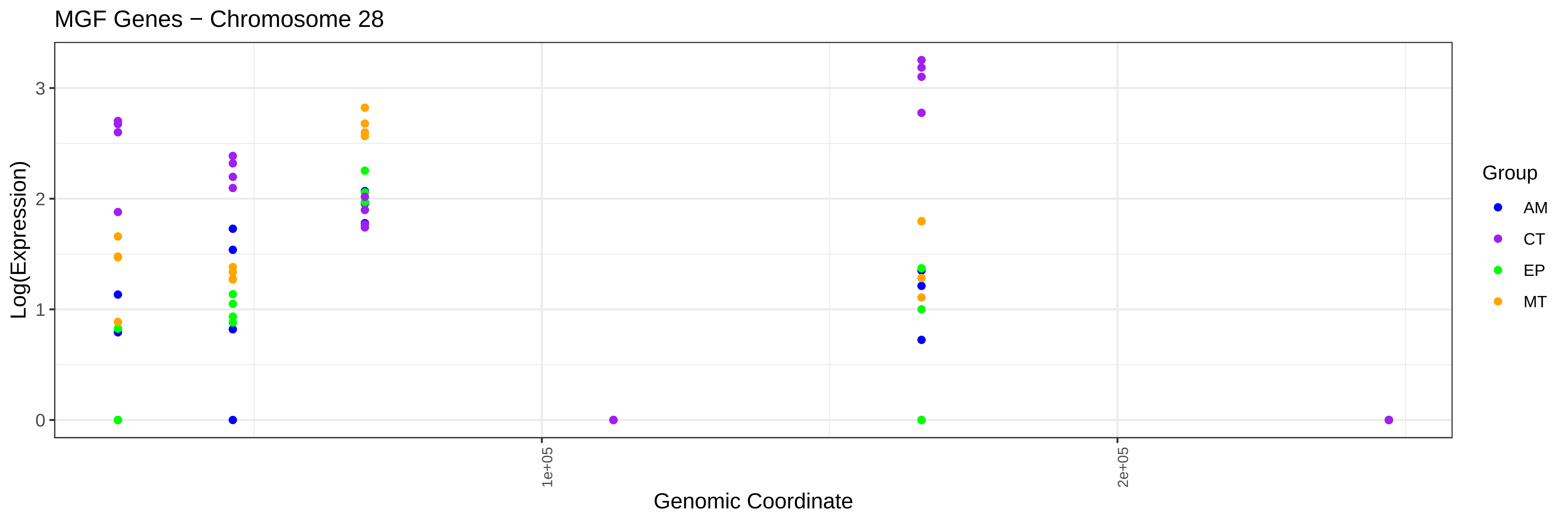
MGF Genes − Chromosome 28
Log(Expression)
0
1
2
3
1e+05
2e+05
Genomic Coordinate
Group
AM
CT
EP
MT

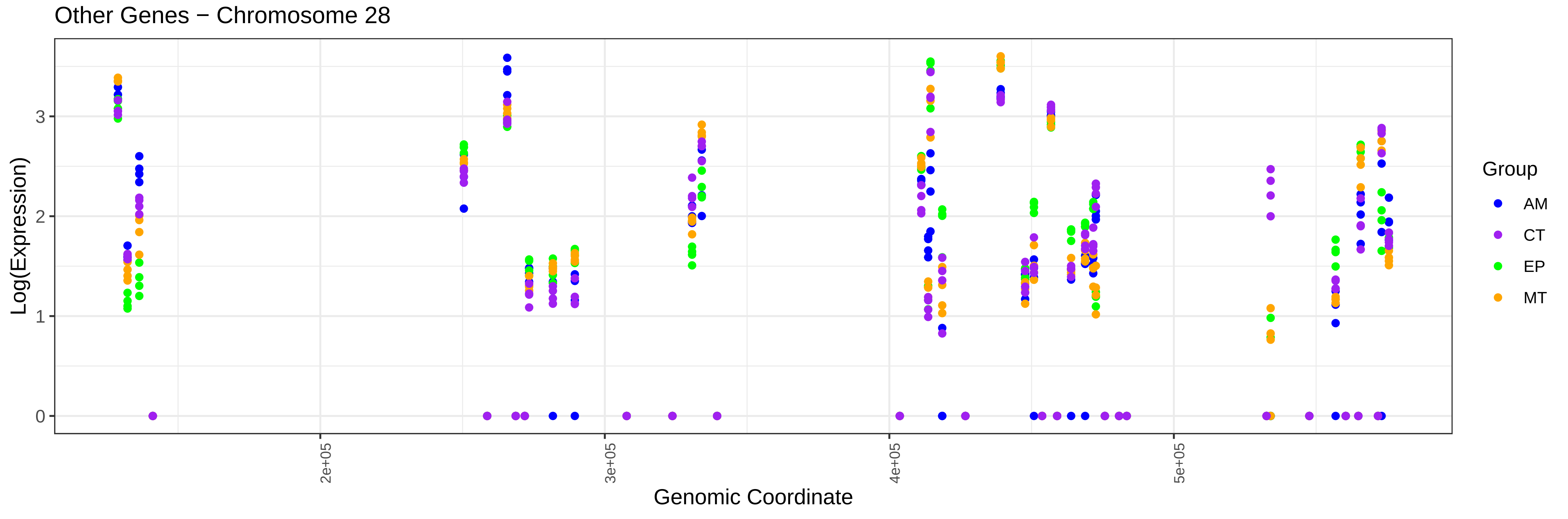
Other Genes − Chromosome 28
Log(Expression)
0
1
2
3
2e+05
3e+05
4e+05
5e+05
Genomic Coordinate
Group
AM
CT
EP
MT

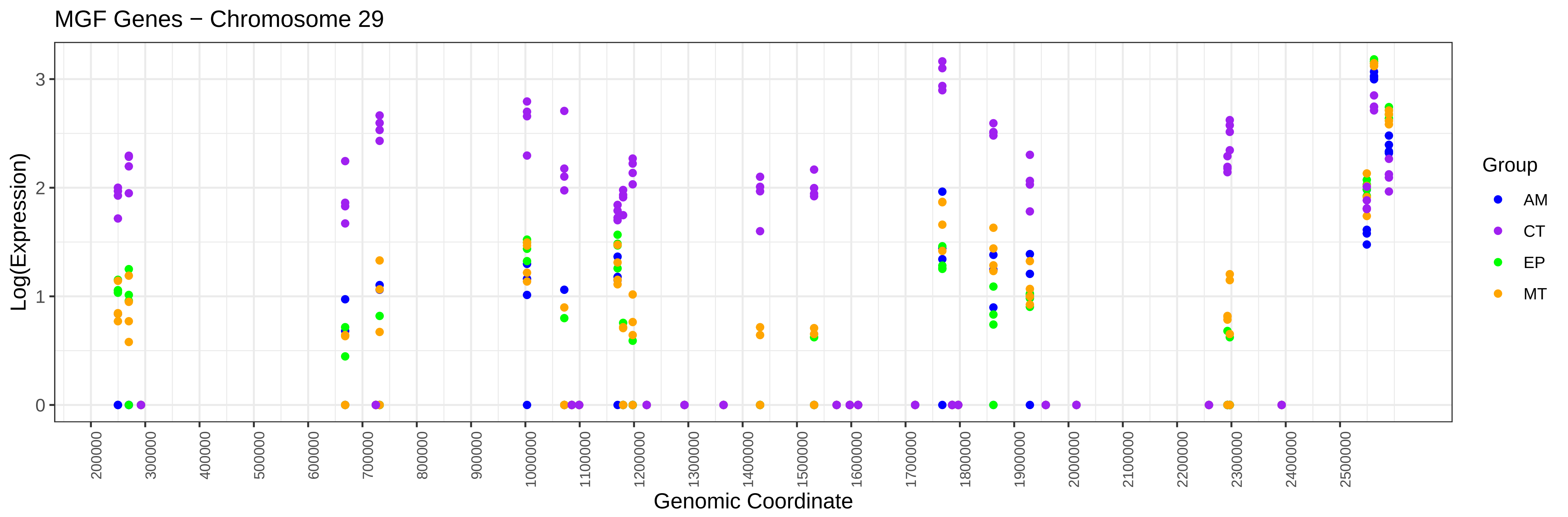
MGF Genes − Chromosome 29
Log(Expression)
0
1
2
3
200000
300000
400000
500000
600000
700000
800000
900000
1000000
1100000
1200000
1300000
1400000
1500000
1600000
1700000
1800000
1900000
2000000
2100000
2200000
2300000
2400000
2500000
Genomic Coordinate
Group
AM
CT
EP
MT

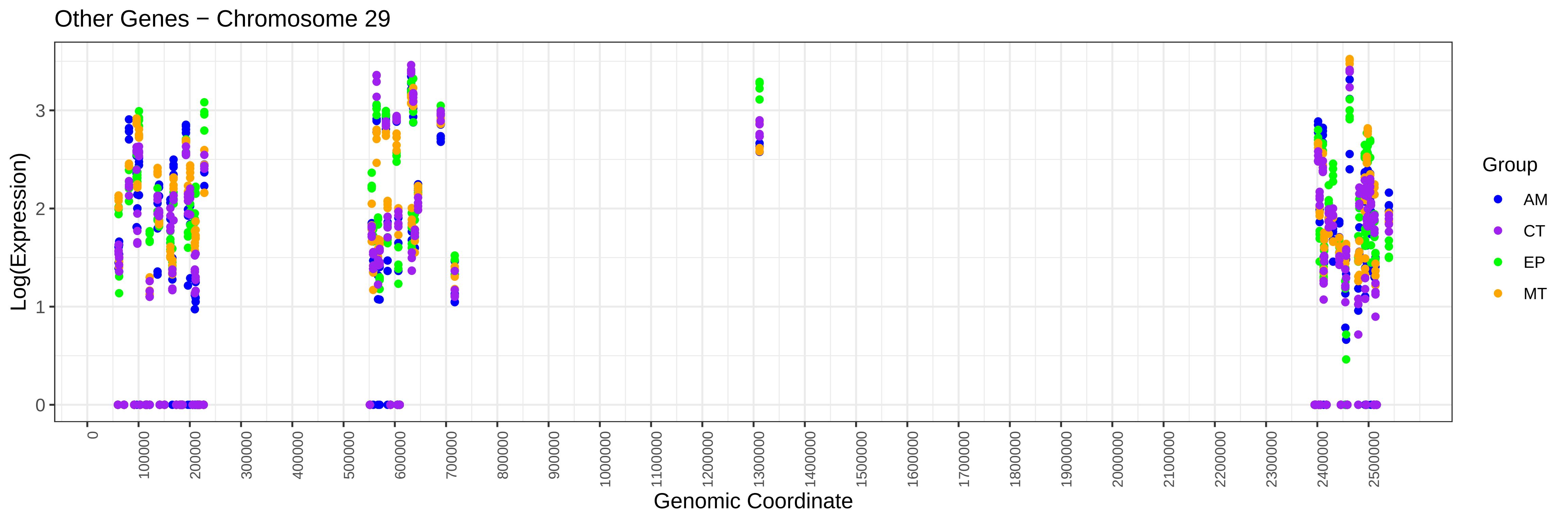
Other Genes − Chromosome 29
Log(Expression)
0
1
2
3
0
100000
200000
300000
400000
500000
600000
700000
800000
900000
1000000
1100000
1200000
1300000
1400000
1500000
1600000
1700000
1800000
1900000
2000000
2100000
2200000
2300000
2400000
2500000
Genomic Coordinate
Group
AM
CT
EP
MT

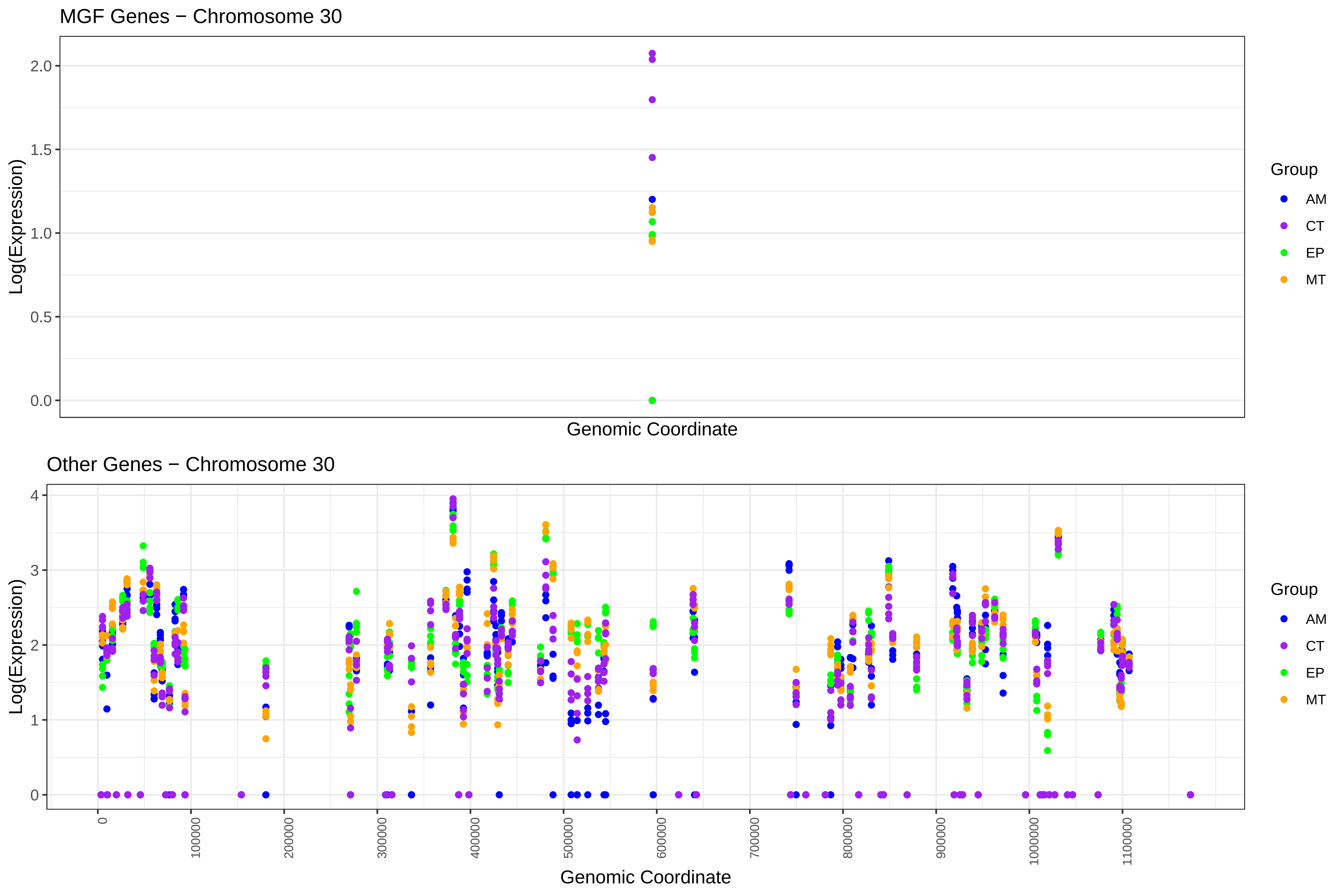
MGF Genes − Chromosome 30
Log(Expression)
2.0
1.5
1.0
0.5
0.0
Genomic Coordinate
Group
AM
CT
EP
MT
Other Genes − Chromosome 30
Log(Expression)
4
3
2
1
0
0
100000
200000
300000
400000
500000
600000
700000
800000
900000
1000000
1100000
Genomic Coordinate
Group
AM
CT
EP
MT

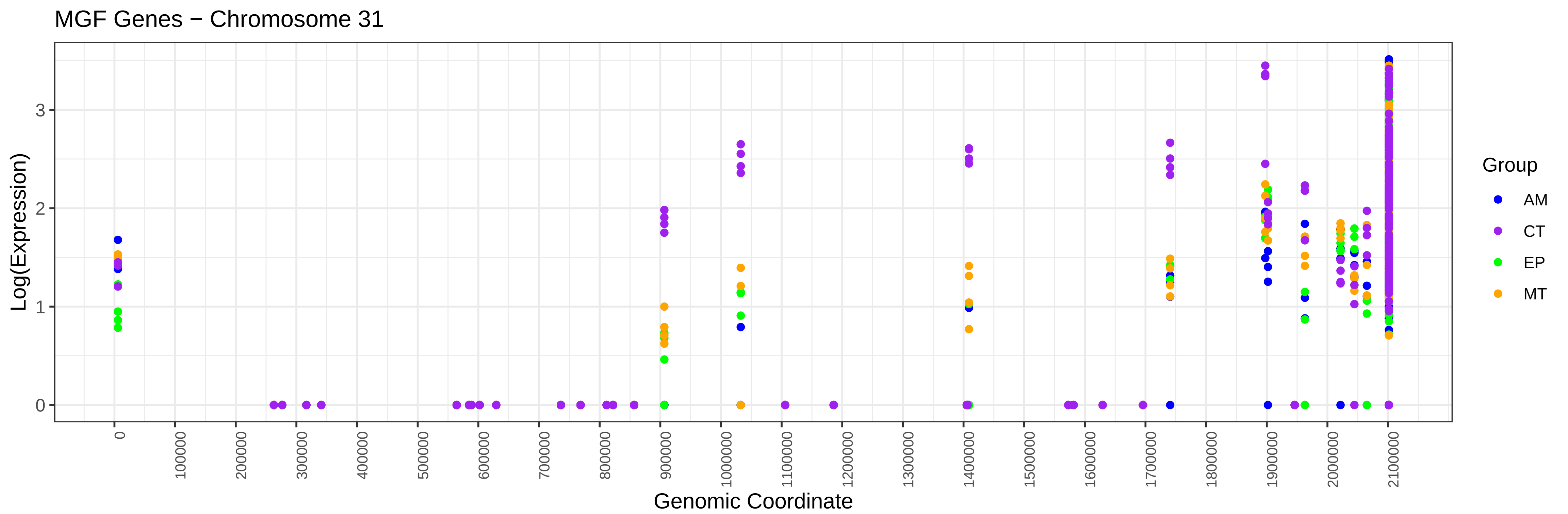
MGF Genes − Chromosome 31
Log(Expression)
0
1
2
3
Genomic Coordinate
0
100000
200000
300000
400000
500000
600000
700000
800000
900000
1000000
1100000
1200000
1300000
1400000
1500000
1600000
1700000
1800000
1900000
2000000
2100000
Group
AM
CT
EP
MT

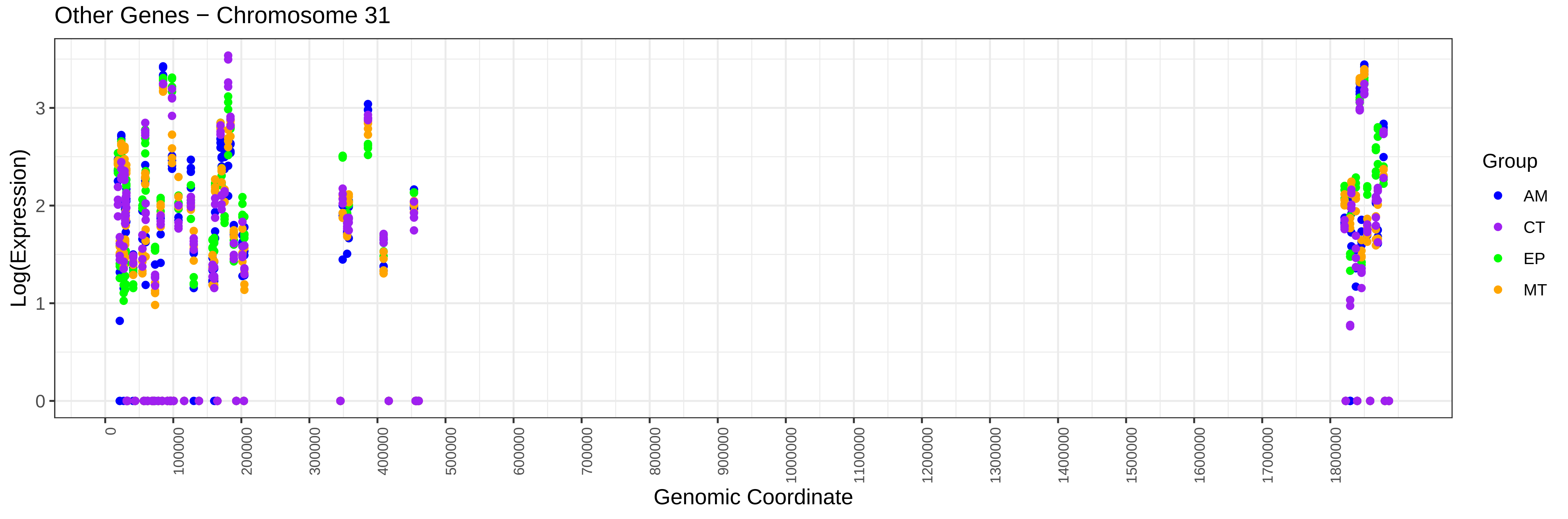
Other Genes − Chromosome 31
Log(Expression)
0
1
2
3
Genomic Coordinate
0
100000
200000
300000
400000
500000
600000
700000
800000
900000
1000000
1100000
1200000
1300000
1400000
1500000
1600000
1700000
1800000
Group
AM
CT
EP
MT
